# Caging-group-free photoactivatable fluorophores with far-red emission

**DOI:** 10.1101/2025.02.24.639818

**Authors:** Richard Lincoln, Alexey N. Butkevich, Clara-Marie Gürth, Alena Fischer, Anna R. Emmerich, Maximilian Fidlin, Mariano L. Bossi, Stefan W. Hell

## Abstract

State of the art super-resolution microscopy methods profit greatly when combined with photoactivatable or photoswitchable fluorophores with far-red emission, but the majority of these fluorophores require specialized buffers or rely on photolabile protecting groups that limit their biocompatibility. We report here a caging-group-free strategy for photoactivatable dyes based on 1-vinyl-10-silaxanthone derivatives containing 9-acylimino or 9-(alkoxycarbonyl)imino groups, enabling minimally-sized photoactivatable dyes with >680 nm emission. The 9-(alkoxycarbonyl)imino silaxanthones are particularly suitable for live-cell labelling, undergoing byproduct-free photoactivation to yield bright and photostable fluorophores that can be readily imaged by STED, PALM, or MINFLUX nanoscopy techniques. The labels derived from these photoactivatable dyes open up new possibilities for multiplexed imaging.

## INTRODUCTION

Red and far-red fluorescent dyes (with excitation maxima, λ_ex._ > 600 nm) are highly valuable tools in super-resolution fluorescent microscopy (optical nanoscopy) techniques, on account of the reduced cellular background, high penetration depth, and low phototoxicity of the incident light.^1^ In single-molecule-based nanoscopy methods,^2,3^ which require addressing sporadic subsets of fluorophores over time, widely used far-red dyes rely heavily on dedicated buffers and additives to achieve stochastic ‘blinking’ processes. These include the inherent on/off behavior of fluorophores (particularly cyanines) under reducing conditions,^4,5^ the sporadic interconversion between fluorescent and non-fluorescent structures of dye scaffolds (including rhodamines^6-9^ and cyanines^10^), or the stochastic binding/unbinding of fluorophores (e.g., PAINT,^11^ DNA-PAINT,^12,13^ or other low-affinity binders^14,15^).

Photoactivatable or caged dyes—in which the off–on transition is irreversible and triggered by light—provide a powerful alternative to stochastic ‘blinking’ systems, with demonstrated performance in quantitative imaging,^16^ color multiplexing, and live-cell tracking.^17^ However, current approaches to photoactivatable far-red fluorophores rely heavily on the synthetic incorporation of photolabile protecting groups that increases molecular weight labels and can limit applications—the few exceptions include rhodamines NN^18-20^ and PA-JF dyes^21^ (both suffering from non-negligible amounts of non-fluorescent byproducts formed during the photoactivation), and SNile Red (thio-caged analogue of Nile Red^22^). Most recently, a photoactivatable PA-SiR dye was proposed,^23^ for which light promoted protonation yielded the fluorescent 9-alkyl-Si-pyronine with limited chemical stability.

We previously described a general strategy for caging-group free photoactivatable dyes through a light-promoted radical cascade reaction (Figure 1A).^24^ This process relies on the photochemical reaction of a 3,6-diaminoxanthone analogue—possessing an excited triplet state with biradical character—and a pendant alkenyl moiety. Following a 6-*endo*-trig cyclization and subsequent protonation by the solvent, this photoreaction yields 9-alkoxy-pyronine fluorophores with blue to orange emission, depending on the bridging atom and auxochromic fragments^16^ (Figure 1B). The resulting dyes and labels derived therefrom performed remarkably well in a number of super-resolution microscopy techniques including STED (stimulated emission depletion),^26^ PALM (photoactivated localization microscopy),^27^ and MINFLUX (minimal photon fluxes)^28^ on account of their live-cell compatibility, high brightness and photostability. Furthermore, the photoactivatable xanthone (PaX) strategy could also be extended to 9-iminoxanthone analogues,^29^ to construct photoactivatable dyes and labels with large Stokes shifts (Figure 1C).^25^ Following photoactivation, Si-xanthone imines bearing an electron-donating acyl amide (**LS1**) or urea (**LS2**) *N*-substituent photoconverted into pyronines with a large (∼100 nm) Stokes shift, on account of a significant hypsochromic shift of their absorption maxima. Yet, despite these efforts, a caging group free photoactivatable fluorophore suitable for imaging with far-red excitation (e.g., 640 nm) still remained elusive.

**Figure 1.**
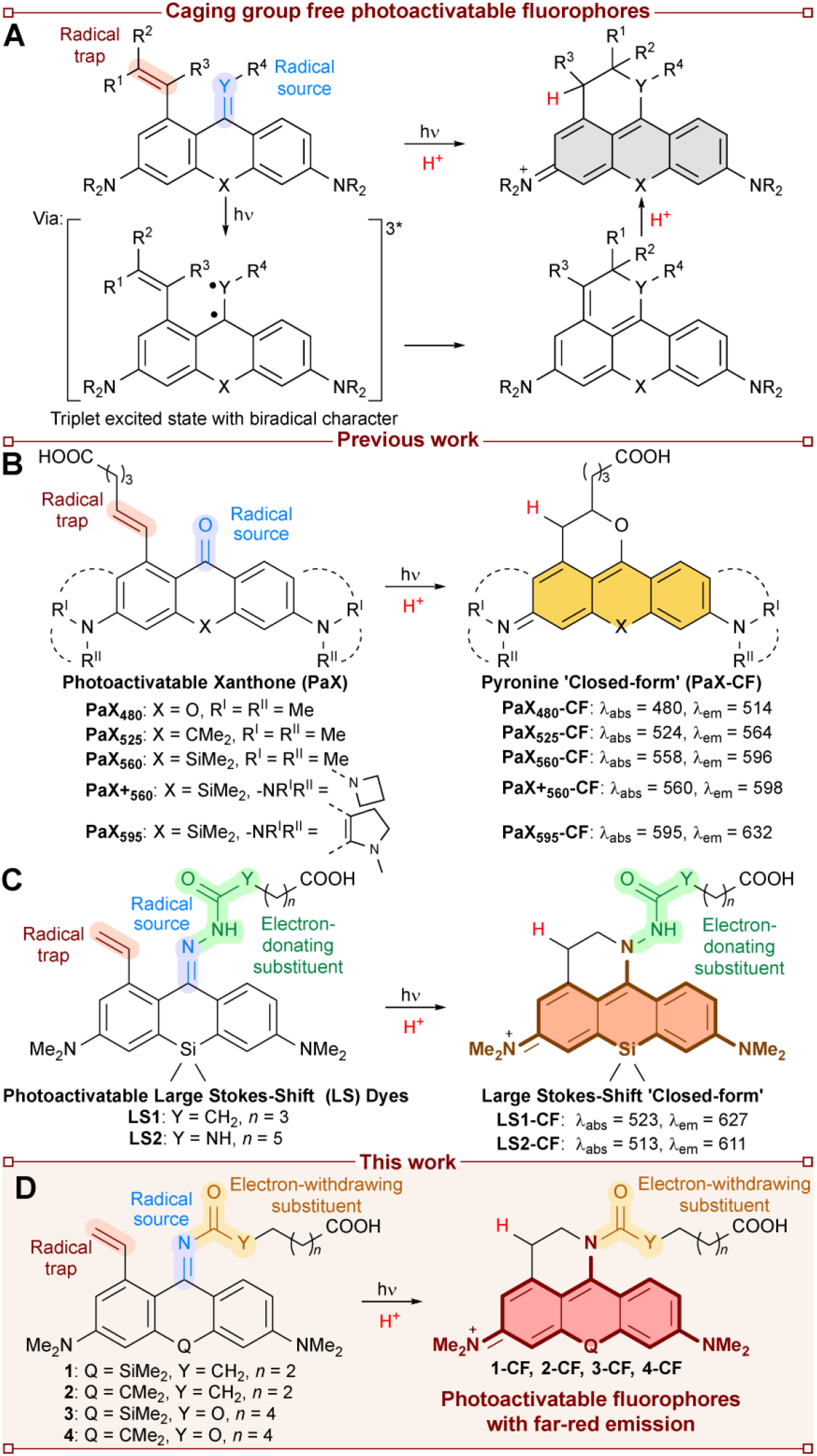
Caging group free photoactivatable dyes for bioimaging. (A) Proposed mechanism of photoactivation based on a radical cascade reaction.^24^ (B) Current examples of photoactivatable xanthone (PaX) dyes.^16,24^ (C) Photoactivatable large Stokes shift dyes.^25^ (D) This work: Photoactivatable fluorophores with far-red emission.

9-Iminopyronines with electron-withdrawing substituents on the imine nitrogen have been shown to enable pH-sensitive fluorophores with far-red emission.^30-33^ Combining these principles with our PaX strategy, we first introduced a 9-acylimino moiety to the PaX scaffold (Figure 1D), which resulted in photoactivatable dyes with red-shifted spectral properties but showing undesirable pH-dependent behavior, leading to lower biocompatibility and chemical stability, and modest photochemical efficiency. These shortcomings were overcome by utilizing a 9-(alkoxycarbonyl)imino moiety, providing a far-red emissive photoactivatable fluorophore with high brightness and photostability. This optimized fluorophore is suitable for a number of live- and fixed-cell labelling strategies, and its photoactivation properties are ideal for super-resolution imaging including STED, PALM, and 3D-MINFLUX.

## RESULTS AND DISCUSSION

### Design and synthesis of photoactivatable dyes

Various strategies were considered to impose a bathochromic shift on the closed-form (CF) of the photoactivatable PaX scaffold, including extending the conjugated system or changing the bridging moiety. As we sought to preserve the minimal size and live-cell compatibility of the photoactivated dye by limiting its electrophilic reactivity, we selected the electron-rich 9-iminopyronines as a starting point for further modifications. Since 9-iminopyronines with electron-donating *N*-substituents demonstrated large hypsochromic shift of absorption maxima (Figure 1C), we opted for electron-withdrawing *N*-substituent knowing that under acidic conditions *N*-protonated *N*-(acylimino)- and *N*-(sulfonylimino)pyronines showed far-red absorption and NIR emission^32,33^ and reasoned that the photoactivation reaction of the corresponding PaX core (Figure 1D) would yield a structurally similar pyronine dye.

It was also essential to suppress any pH-sensitivity of the open-form (OF) structure, to maximize signal contrast upon photoactivation. We hypothesized that a milder electron-withdrawing group, *N*-alkoxycarbonyl,^34^ would sufficiently lower the OF p*K*_a_ below the biologically relevant pH window (4.5–8.5) while still yielding the desired bathochromic shift in the CF. To test this hypothesis, four 9-(acylimino)xanthones modified with dimethylsilylene (**1**) and isopropylidene (**2**) bridges were prepared along with the corresponding 9-(alkoxycarbonyl)imino analogues (**3** and **4**, respectively). In their OF structure, the prepared dyes **1**–**4** exist in an equilibrium between a blue-absorbing 9-iminoxanthone (λ_max_ < 400 nm) and a red-shifted 9-aminopyronine structure (λ_max_ > 600 nm; Figure S1A).

To determine the p*K*_a_ of this equilibrium, the absorption spectra of compounds **1**–**4** were recorded at various pH values (in phosphate-buffered saline adjusted to the target pH) across the biologically relevant range (Figures 2A and S1B–E). A clear transition from the iminoxanthone to aminopyronine structure was observed for all of the compounds screened. Consistent with previous reports of 9-(acylimino)xanthones,^33^ silicon-bridged compounds showed a lower p*K*_a_ than their carbon-bridged analogues (e.g., 4.97 vs 6.94 for **1** and **2** respectively). As inferred above, replacement of 9-acylimino with 9-(alkoxycarbonyl)imino group resulted in a significant reduction of the p*K*_a_ for compounds **3** and **4** compared to the corresponding *N*-acylimines. In the case of compound **3**, its p*K*_a_ value was sufficiently outside of the biological window, even for the most acidic cell compartments (lysosomes and late endosomes), making it an optimal candidate for live-cell applications.

**Figure 2.**
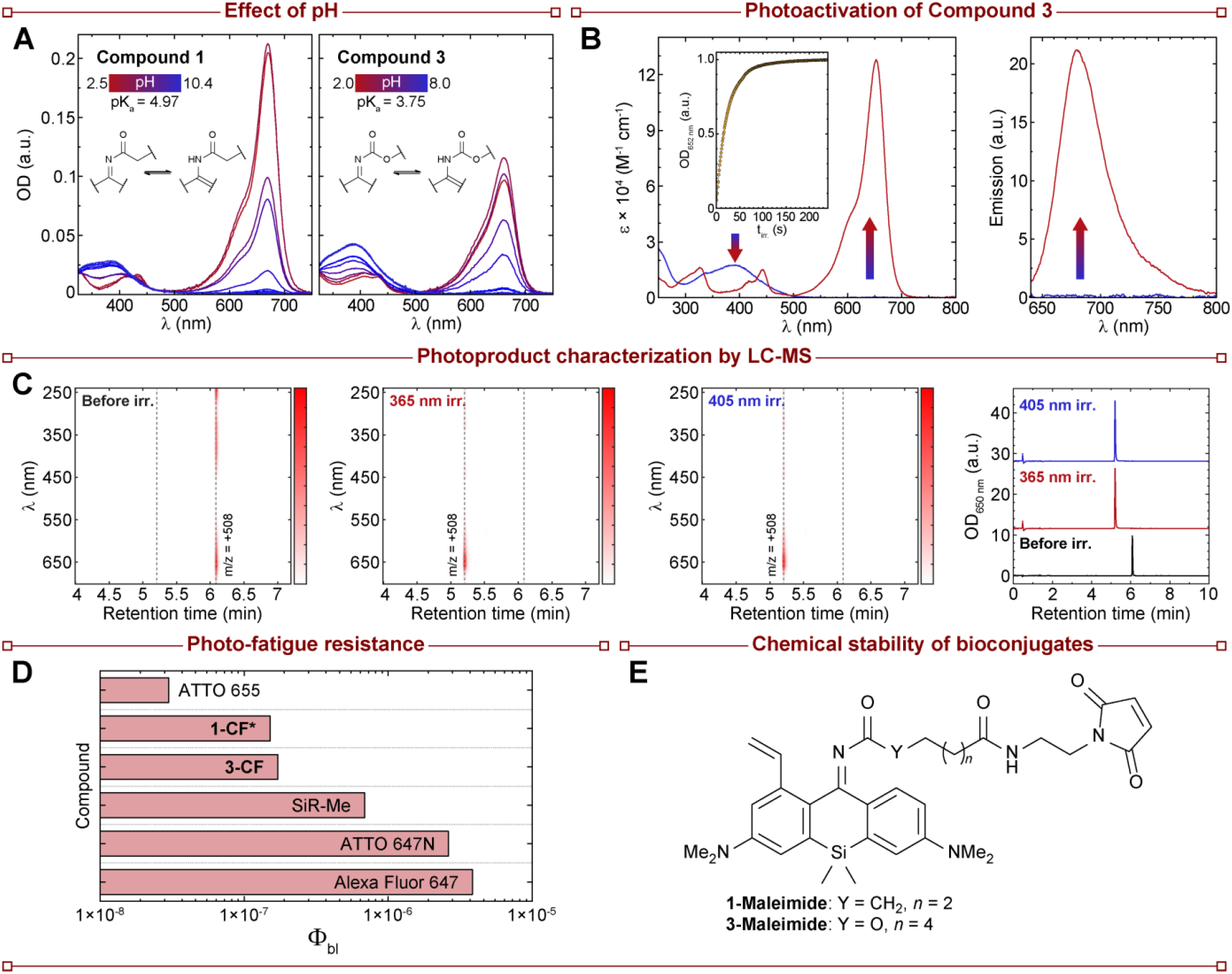
Characterization of far-red photoactivatable dyes. (A) Effect of pH on the open-form structures of **1** and **3**. (B) Temporal evolution of the absorption (left) and fluorescence (right) spectra of **3** (**3** → **3-CF**; 5 μM) irradiated in phosphate buffer (100 mM, pH = 7; λ_act_ = 365 nm). (C) LC-MS chromatograms of the reaction mixtures of compound **3** before (left) and after photoactivation with either 365 nm (center-left) or 405 nm (center-right) irradiation. The optical density at 650 nm is shown on the right, corresponding to either the protonated form of **3** (before irradiation) or **3-CF** after irradiation. (D) Quantum yield of photobleaching (Φ_bl_) of compounds **1-CF\*** and **3-CF** compared to commonly employed far-red fluorophores. (E) Structures of maleimide derivatives of **1** and **3** for assessment of chemical stability of bioconjugates.

We next studied the photoactivation behavior of the prepared compounds. Dilute (5 μM) solutions of compounds **1**–**4** in phosphate buffer (100 mM, pH = 7) were irradiated with either 365 nm or 405 nm light (Figures 2B and **S2**). For the *N*-acylimines **1** and **2**, a non-negligible initial fraction of the protonated form was present at the pH of the buffer. Irradiation resulted in the formation of the corresponding close-form structures (**1-CF** and **2-CF**) with complex reaction kinetics, suggesting possible formation of an intermediate product, which however proved too elusive for characterization by LCMS (only a single final product was detected, Figure S2). The carbamate-bearing compounds (**3, 4**) showed less of the protonated form at neutral pH than the corresponding acylimines, and photoconverted cleanly and completely to the corresponding close-form (**3-CF** and **4-CF**) upon irradiation, without evidence of intermediates on the observation timescale (Figure 2C for compound **3**). These results indicate that the carbamate modification results in a more robust and efficient photoswitching behavior, which is critical for microscopy applications. Due to the non-negligible amounts of aminopyronine form at physiological pH, the carbon-bridged compounds **2** and **4** are likely to be poor candidates for live-cell microscopy applications and were not studied further. We note however that compound **4** may find further practical use in fixed-cell multicolor imaging or material science applications where a photoactivatable dye with red (610 nm) absorption is needed to fill the gap between orange-red and far-red markers.

To determine the photofatigue resistance of the closed-form products **1-CF** and **3-CF**, we performed photobleaching studies benchmarking the dyes against a series of commercially available fluorophores with similar spectral properties (Figures 2D and S3). To avoid contributing errors from the intermediates formed along the photoactivation of **1**, we prepared and isolated the closed-form analogue **1-CF\*** with an ω-carboxyalkyl linker truncated to a methyl group (Figure S3A). Both **1-CF\*** and **3-CF** showed remarkable photostability when compared to the commonly-employed far-red fluorophores, including a cyanine (**AF647**), a carbopyronine (**ATTO 647N**), and a silicon rhodamine (**SiR-Me**) (Figure S3B). Only the oxazine dye **ATTO 655** demonstrated greater photostability under the studied conditions (i.e. in pH 7 phosphate buffer without additives).

Lastly, we assessed the chemical stability of the imine linkage critical for bioconjugate labelling. For this, we prepared the maleimide analogues **1-Maleimide** and **3-Maleimide** (Figures 2E) and conjugated them to commercially available nanobodies (single-domain camelid antibodies) carrying a single ectopic cysteine residue. The nanobodies were characterized by mass spectrometry before and after labelling, and again after one week of storage at 4°C (Figures S4 and S5). For the compound **1-Maleimide**, we observed two products formed during the labelling. The major fraction of detected mass was consistent with a single added fluorophore (M+600), however, another large fraction of labelled nanobodies carried a fragment with the mass (M+266), which we assigned to the primary amide resulting from the imine hydrolysis. Prolonged storage of the nanobodies further increased the amount of the hydrolyzed material. On the contrary, for the compound **3-Maleimide**, complete labelling was observed upon the conjugation, and very little hydrolysis was noticed after one week of storage.

The chemical and spectral properties for the studied compounds are summarized in Table 1. Taken together, these results highlight the significant improvements in biocompatibility, (photo-)chemical stability imparted by the alkoxycarbonyl modification, especially when incorporated in silicon-bridged xanthones. As such, compound **3** was chosen as the primary compound used in subsequent imaging experiments.

**Table 1.** Properties of Photoactivatable Dyes 1–5 before and after Activation Before Photoactivation After Photoactivation.

| Before Photoactivation |  |  |  | After Photoactivation |  |  |  |
| --- | --- | --- | --- | --- | --- | --- | --- |
| Dye | Abs. $\lambda_{\max}$, nm<br>($\epsilon$, M<sup>-1</sup> cm<sup>-1</sup>) | pK<sub>a</sub> | $\phi_{\text{Act}}$ | CF-Dye | Abs. $\lambda_{\max}$, nm<br>($\epsilon$, M<sup>-1</sup> cm<sup>-1</sup>) | Em. $\lambda_{\max}$, nm ($\Phi_f$) | Fl. lifetime $\tau$, ns |
| **1** | 389 (14500) | 4.97 | 7.1E-3<sup>b</sup> | 1-CF | 658 (72000)<sup>c</sup> | 690 (0.21) | 1.64 |
| **2** | 407 (14000) | 6.94 | 8.0E-3<sup>b</sup> | 2-CF | 617 (66000)<sup>c</sup> | 655 (n.d.) | n.d. |
| **3 (PaX<sub>650</sub>)** | 390 (19000) | 3.75 | 6.2E-3 | 3-CF | 652 (124000) | 680 (0.28) | 2.10 |
| **4** | 401 (21000) | 5.15 | 1.3E-2<sup>a</sup> | 4-CF | 610 (91000) | 648 (0.41) | 2.78 |
| **5 (PaX<sub>695</sub>)** | 356 (8500) | n.d. | 2.8E-3 | 5-CF | 696 (67000) | 724 (0.09) | 0.92 |

<sup>a</sup> Contributions from the protonated form were ignored.
<sup>b</sup> Transient at 365 nm because of a slow thermal reaction coupled as a second step.
<sup>c</sup> May be incomplete due to the slow thermal reaction (second step).

### Far-red emissive labels for fixed- and live-cell imaging

We next set out to prepare the targeted labels of compound **3** for live-cell fluorescence microscopy experiments (Figure 3). We focused our attention on constructing probes for self-labelling protein tags.^35^ The corresponding small-molecule ligands (Figure 3A) for the two most-commonly applied self-labelling proteins have been prepared: a linear ω-chloroalkane with PEG2 linker (**3-Halo)** specific for HaloTag^36^ (an engineered bacterial haloalkane dehalogenase) and a *O*^6^-benzylguanine derivative (**3-BG**) specific for the SNAP-tag^37^ (a modified *O*^6^-methylguanine-DNA methyltransferase).

**Figure 3.**
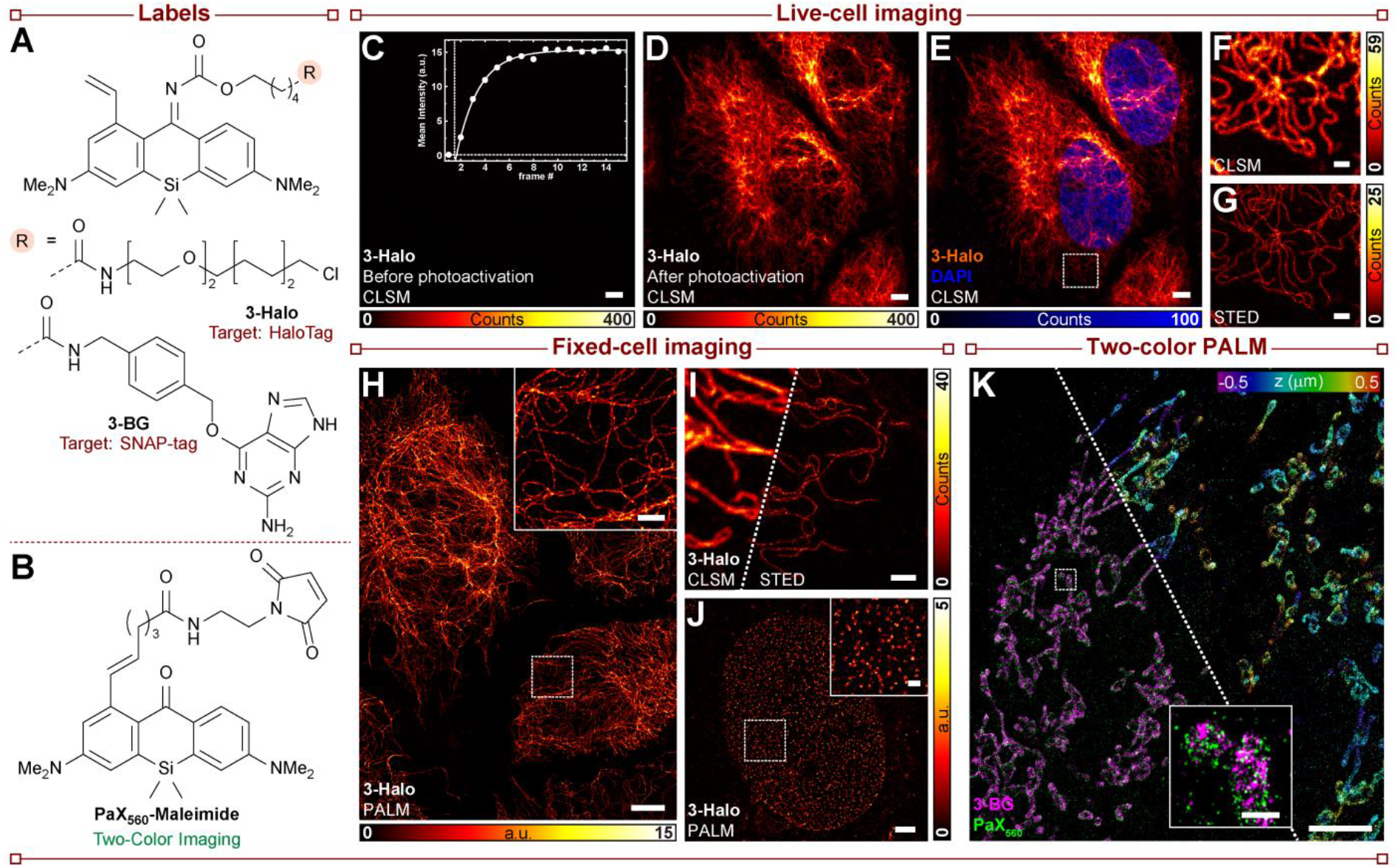
Photoactivatable far-red labels for live- and fixed-cell imaging. (A) Structures of derivatives of **3** for live cell labelling of HaloTag (**3-Halo**) or SNAP-tag (**3-BG**) self-labelling protein tags. (B) Structure of **PaX**_**560**_**-Maleimide** used in two-color PALM imaging. (C) Confocal image of live U-2 OS cells stably expressing a vimentin-HaloTag construct labelled with **3-Halo** (500 nM) before photoactivation. Inset: Change in intensity upon photoactivation with 405 nm laser light. Scale bar: 5 μm. (D–E) Confocal images of the same sample as (C) following photoactivation. Scale bars: 5 μm. (F–G) Confocal (F) and STED (G) images of the region marked in E. Scale bars: 1 μm. (H) PALM image of fixed U-2 OS cells stably expressing a vimentin-HaloTag construct labelled with **3-Halo**. Inset: Magnified region marked in the overview image. Scale bars: 5 μm (overview) and 1 μm (inset). (I) Confocal (left) and STED (right) images of fixed U-2 OS cells stably expressing a vimentin-HaloTag construct labelled with **3-Halo** (500 nM) following photoactivation with 405 nm laser light. Scale bar: 1 μm. (J) PALM image of nuclear pore complexes in fixed U-2 OS cells stably expressing a NUP96-HaloTag construct labelled with **3-Halo** (500 nM). Inset: Magnified region marked in the overview image. Scale bars: 2 μm (overview) and 500 nm (inset). (K) Two-color 3D-PALM imaging of fixed HeLa cells stably expressing a COX8A-SNAP-tag construct labelled with **3-BG** (1 μM) and co-labelled via indirect immunofluorescence with an anti-TOMM20 primary antibody and a secondary nanobody labelled with PaX_560_-Maleimide. Left: Localizations color coded by imaging channel for **3** (magenta) and PaX_560_ (green). Right: Combined localizations of **3** and **PaX**_**560**_ color coded by Z-position. Inset: Magnified region marked in the overview image. Scale bars: 5 μm (overview) and 500 nm (inset).

To test the live-cell permeability of **3-Halo**, U-2 OS cells stably expressing a vimentin-HaloTag construct^38^ were incubated with the chloroalkane derivative (500 nM, overnight). After washing, the cells were imaged by confocal microscopy. Initially, no fluorescence was observed in the far-red channel (Figure 3C), but upon photoactivation with 405 nm laser light, the vimentin filaments were revealed (Figure 3D–E). The imaging with sub-diffraction resolution in living cells was performed by STED (stimulated emission depletion) microscopy (Figure 3F–G) or PALM (photoactivated localization microscopy; Figure S6). Fixing the samples with paraformaldehyde (PFA) after labelling prevented cell dynamics during imaging, and further enhanced the number of detected photons in vimentin filaments imaged by PALM (Figure 3H) and STED (Figure 3H) as well as in PALM imaging of nuclear pore complexes (NPCs, Figure 3J) in the homozygous knock-in U-2 OS cell line endogenously expressing a fusion construct of nucleoporin Nup96 and HaloTag.^39^

To further demonstrate the utility of our labels, we performed two-color 3D-PALM imaging of fixed mitochondria by combining compound **3** with another spectrally discernable photoactivatable PaX label. Live-cell labelling of a COX8a-SNAP-tag fusion protein was first performed with **3-BG**, and following paraformaldehyde fixation, a secondary immunofluorescence labelling of the mitochondria outer membrane protein TOMM20 was performed using nanobodies carrying **PaX**_**560**_ dye (**Figure 3K**).^24^ As it was determined that 560 nm excitation was sufficient to photoactivate compound **3** (but not **PaX**_**560**_), we took advantage of sequential imaging to independently select the two fluorophores. First, compound **3** was imaged with 640 nm excitation (270 mW) using the 560 nm laser as the activation laser (<0.1 mW to 3 mW). Upon depleting the localizations of **3, PaX**_**560**_ was in turn imaged using the 560 nm laser for excitation (180 mW) and a 405 nm laser for photoactivation (<0.1 mW), see Figure S7.

### Applications in MINFLUX nanoscopy

The photoactivatable dye **3** was next tested in MINFLUX nanoscopy,^28,40^ a technique that utilizes a shaped (e.g., a donut) excitation beam with an intensity minimum to localize individual fluorophores. By exploiting the known position of the excitation zero, the fluorophores can be localized with single-digit nanometer precisions with fewer emitted photons compared to camera-based localization. In the case of photoactivatable fluorophores, once in the on-state, the position of the single molecule can often be repeatedly determined several times before its destruction via photobleaching, further enabling the statistical analysis of these combined localizations for molecular positioning down to angstrom precisions.^41^

To test the detection efficiency of compound **3** in 640 nm MINFLUX imaging, we utilized the Nup96-HaloTag fusion cell line—a robust reference standard for quantitative super-resolution imaging.^39^ 2D MINFLUX imaging of fixed samples labelled with **3-Halo** on a commercial MINFLUX microscope (Figure 4A and Figure S8B) revealed the expected NPC structures with circumscribed labelling, enabling us to characterize the fluorophore performance in a consistent manner. For compound **3-Halo**, we obtained an average localization precision of 3.2 nm (Figure S8C), with each single molecule localized an average (median) of 29.0 times with an average of 179 photons used per localization (Figure S8D–F). We evaluated the occupancy of HaloTag sites in the nuclear pore complexes as a measure of the labelling efficiency of **3-Halo** (Figure 4B), having obtained a value of 6, which is similar to the previously reported value of **PaX**_**560**_^16^ in MINFLUX, and of other chloroalkane-tagged dyes reported on PALM/STORM.^39^

**Figure 4.**
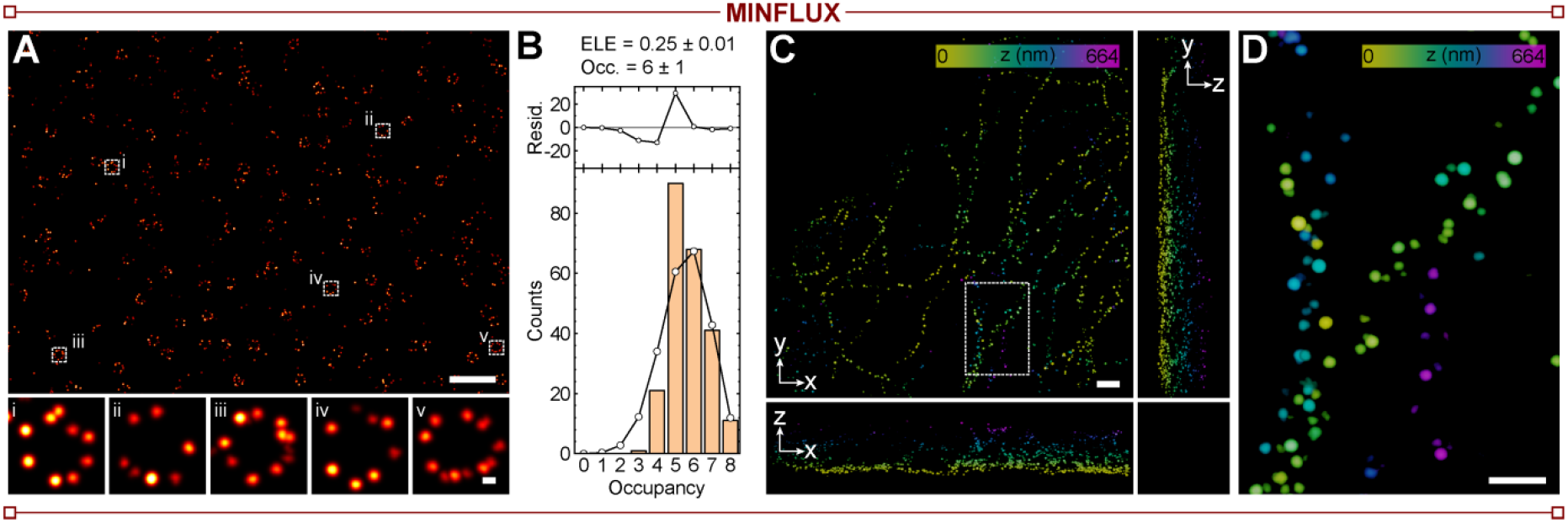
MINFLUX imaging. (A) 2D-MINFLUX image of U-2 OS cells stably expressing a NUP96-HaloTag construct labelled compound **3-Halo** (500 nM, overnight), fixed and mounted in Mowiol prior to imaging. Bottom row: individual NPCs marked in the overview image. Scale bars: 500 nm (overview) and 20 nm (bottom row). (B) Normalized occupancy histogram of NPCs labeled with **3-Halo**. A fit to a probabilistic model is included in the histogram (bars: experimental data, circles: fit) considering eight labeling sites and four protein copies at each one. Residuals are shown in the top plot (See Supporting Information). (C) Three-dimensional MINFLUX image of U-2 OS cells stably expressing a vimentin-HaloTag construct labelled compound **3-Halo** (500 nM, overnight), fixed and mounted in Mowiol prior to imaging. Scale bar: 200 nm. (D) Magnified region marked in C. Scale bar: 100 nm.

Finally, we used compound **3-Halo** for the 3D mapping of vimentin-HaloTag fusion proteins in the U-2 OS cell line with 3D-MINFLUX imaging, which allowed us to see the individual vimentin filaments and their helical bundling (Figure 4C–D). We envision that further adoption of these probes, especially when combined with MINFLUX imaging, will enable new opportunities for quantitative fluorescence imaging and structural biology.

### Further bathochromic shift enables dual color 640 nm MINFLUX

To push the absorption maxima of PaX fluorophores beyond the far-red range, for potential applications including multiplexing and deep tissue imaging, we rationalized that fusing indoline auxochromic fragments to the 10-silaxanthone core (similar to SiR700 silicon-rhodamine dye^42^) will provide a further bathochromically shifted analogue of **3**, a 9-(alkoxycarbonyl)imino derivative of the dye **PaX**_**595**_.^16^ With further optimization of the synthetic sequence reported for **PaX**_**595**_, compound **5** along with its ω-chloroalkane (**5-Halo**), *O*-benzylguanine (**5-BG**), NHS-ester (**5-NHS**), and maleimide (**5-Maleimide**) derivatives were prepared (Figure 5A). Photoactivation of **5** yields a closed-form (**5-CF**) with an emission maximum of 724 nm (Table 1 and Figure S9A) upon clean photoactivation (Figure S9B). The derived labels can be used in live-cell imaging (**5-Halo** and **5-BG**, Figure 5C–D) and fixed samples (**5-NHS** and **5-Maleimide**, Figure S9E–F) employing the same excitation wavelength as previously established for compound **3** (640 nm).

**Figure 5.**
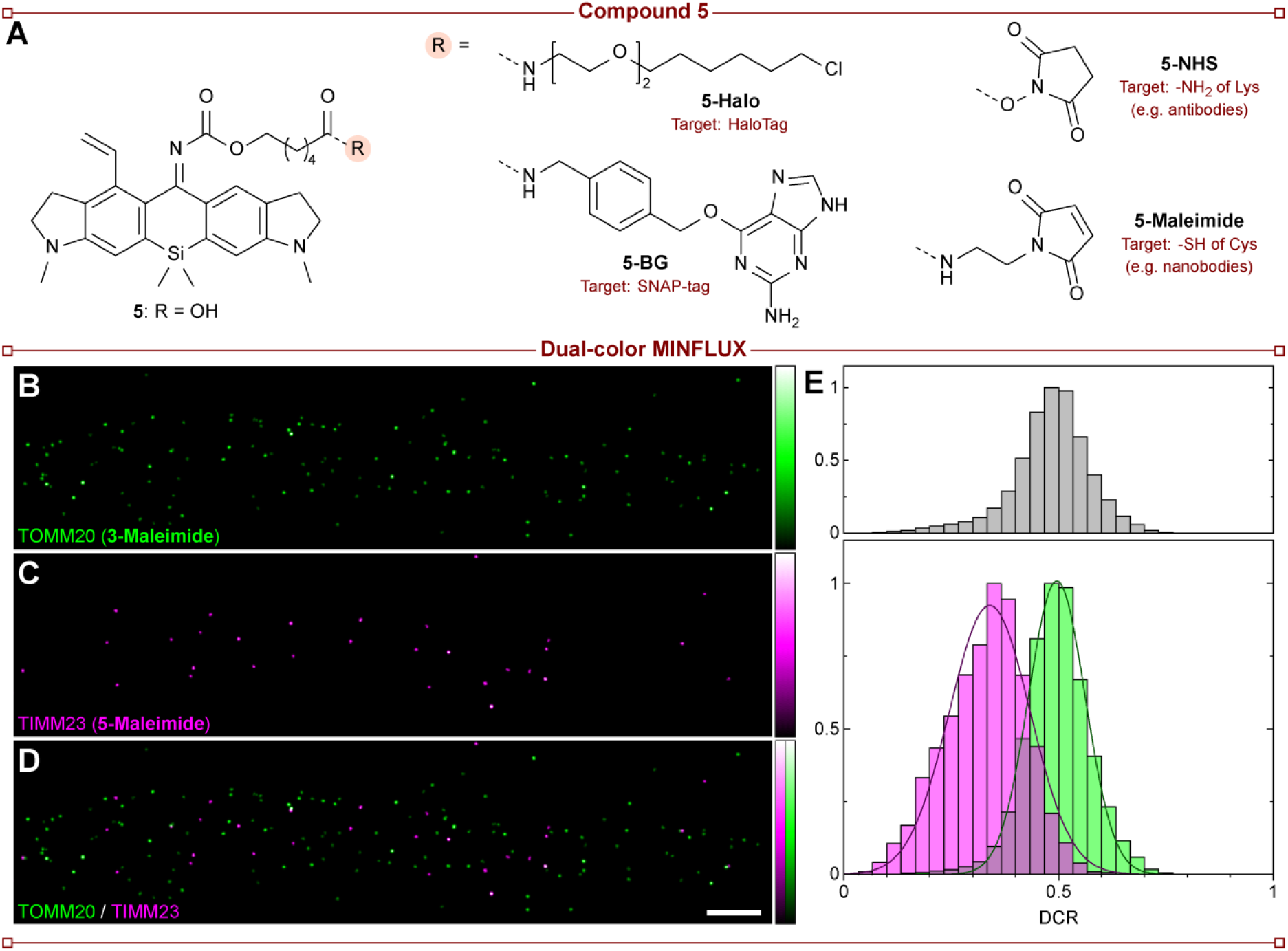
Bathochromically-shifted photoactivatable dye for dual-color MINFLUX imaging by spectral separation. (A) Structures of compound **5** and its derivatives of live cell labelling of HaloTag (**5-Halo**) or SNAP-tag (**5-BG**) self-labelling protein tags and fixed cell labelling (**5-NHS** and **5-Maleimide**). (B–D) Dual color 640 nm MINFLUX imaging of mitochondria proteins TOMM20 (B) and TIMM23 (C) labelled by secondary immunofluorescence with secondary nanobodies conjugated to compound **3-Maleimide** and **5-Maleimide**, respectively, and the combined two-color image (D). Scale bar: 200 nm. (E) Normalized histograms of the DCR value of all localizations (grey) and the ones after spectral classification (green/magenta).

Lastly, we leveraged the difference in emission spectra of compounds **3** and **5** for dual color MINFLUX imaging based on spectral classification (Figure 5B–D).^43^ Using the same wavelength (640 nm) for excitation of both fluorophores, the fluorophore type was classified by splitting the single molecule emission into two spectral ranges (650-685 nm and 685–710 nm), detected by two dedicated photon-counting detectors. The mean detector channel ratio (DCR) for all localizations belonging to a single molecule event was used to determine the emitting fluorophore (Figure 5E). Using this strategy, we visualized the mitochondrial proteins TOMM20 and TIMM23 orthogonally labelled by indirect immunofluorescence with secondary nanobodies tagged with **3-maleimide** and **5-maleimide**, respectively. We obtained comparable localization precisions (2 nm) for the two fluorophores (Figure S10), with more detected photons and localizations per single molecule event for compound **3**, which can be rationalized by its better spectral overlap with the available channels, and the diminishing detector efficiency for far-red photons.

## CONCLUSIONS

Taken together, our results highlight the utility and versatility of 9-(alkoxycarbonyl)imino silaxanthones as a platform for caging-group free fluorophores with far-red emission. The compounds reported in this work, particularly **PaX**_**650**_ and **PaX**_**695**_, expand the family of PaX dyes to cover the entire visible spectrum of light, yielding live-cell compatible labels that can be used in multicolor STED, PALM, or MINFLUX nanoscopy techniques. Rapid and byproduct-free activation of the new red-emitting PaX fluorophores **3** and **5** in biological media will make them particularly appealing in quantitative super-resolution microscopy applications, single-molecule tracking and protein-protein interaction studies with molecular precision. Beyond their use in microscopy, we also anticipate that these photoactivatable dyes will further stimulate the development of light-controlled probes and sensors for biological and material science.

## EXPERIMENTAL PROCEDURES

### Resource availability

#### Lead contact

Further information and requests for resources should be directed to and will be fulfilled by the lead contact, Alexey N. Butkevich.

#### Materials availability

Requests for materials generated during this study can be made to.

#### Data and code availability

The datasets generated during this study are available from the from the corresponding authors upon reasonable request.

## SUPPLEMENTAL INFORMATION

Supplemental experimental procedures, Figures S1–S10 and Tables S1–S5, and synthesis and characterization of compounds.

## ACKNOWLEDGMENTS

The research was funded by Bundesministerium für Bildung und Forschung (German Federal Ministry of Education and Research), Project No. 13N14122 “3D Nano Life Cell” (to S.W.H.) and the Deutsche Forschungsgemeinschaft (DFG, SFB1286/A07 to S.W.H.). R.L. is grateful to the Max Planck Society for a Nobel Laureate Fellowship. We thank the EMBL for providing the U-2 OS-CRISPR-NUP96-Halo clone #252 (330448, CLS) and Prof. S. Jakobs (Max Planck Institute for Multidisciplinary Sciences, University of Göttingen) for providing the U2OS-Vim-Halo and HeLa-COX8A-SNAP cell lines. We thank the staff of the Mass Spectrometry Core Facility and the Optical Microscopy Facility at the Max Planck Institute for Medical Research for their support.

## AUTHOR CONTRIBUTIONS

R.L. was responsible for the project’s conception and wrote the manuscript with input from A.N.B., M.L.B. and S.W.H. A.N.B. and R.L. performed the chemical synthesis. M.L.B., A.R.E., M.F. and R.L. performed dye characterization and mechanistic studies. A.F. performed cell culture and labelling. M.L.B., R.L., and C.-M.G. performed microscopy. M.L.B. and R.L. performed data analysis. S.W.H. acquired funding, directed and supervised the investigations. All the authors discussed the results and commented on the manuscript.

