## Supplementary Material for "Caging-group-free photoactivatable fluorophores with far-red emission"

***This section includes:***

SUPPLEMENTARY REFERENCES ... 41
NMR SPECTRA ... 43
Compound S2 $^{1}H$ ... 43
Compound S2 $^{13}C$ ... 44
Compound S3 $^{1}H$ ... 45
Compound S3 $^{13}C$ ... 46
Compound 1-CF* $^{1}H$ ... 47
Compound 1-CF* gHSQCad ... 48
Compound S4 $^{1}H$ ... 49
Compound S4 $^{13}C$ ... 50
Compound 1 $^{1}H$ ... 51
Compound 1 $^{13}C$ ... 52
Compound S6 $^{1}H$ ... 53
Compound S6 $^{13}C$ ... 54
Compound S7 $^{1}H$ ... 55
Compound S7 $^{13}C$ ... 56
Compound S8 $^{1}H$ ... 57
Compound S8 $^{13}C$ ... 58
Compound 2 $^{1}H$ ... 59
Compound 2 $^{13}C$ ... 60
Compound 3 $^{1}H$ ... 61
Compound 3 $^{13}C$ ... 62
Compound 4 $^{1}H$ ... 63
Compound 4 $^{13}C$ ... 64
Compound 1-Maleimide $^{1}H$ ... 65
Compound 1-Maleimide $^{13}C$ ... 66
Compound 3-Halo $^{1}H$ ... 67
Compound 3-Halo $^{13}C$ ... 68
Compound S13 $^{1}H$ ... 69
Compound S13 $^{13}C$ ... 70
Compound S14 $^{1}H$ ... 71
Compound S14 $^{13}C$ ... 72
Compound 5 $^{1}H$ ... 73
Compound 5 gHSQCad ... 74
Compound 5-Halo 1H ... 75
Compound 5-Halo gHSQCad ... 76

### SUPPLEMENTARY FIGURES

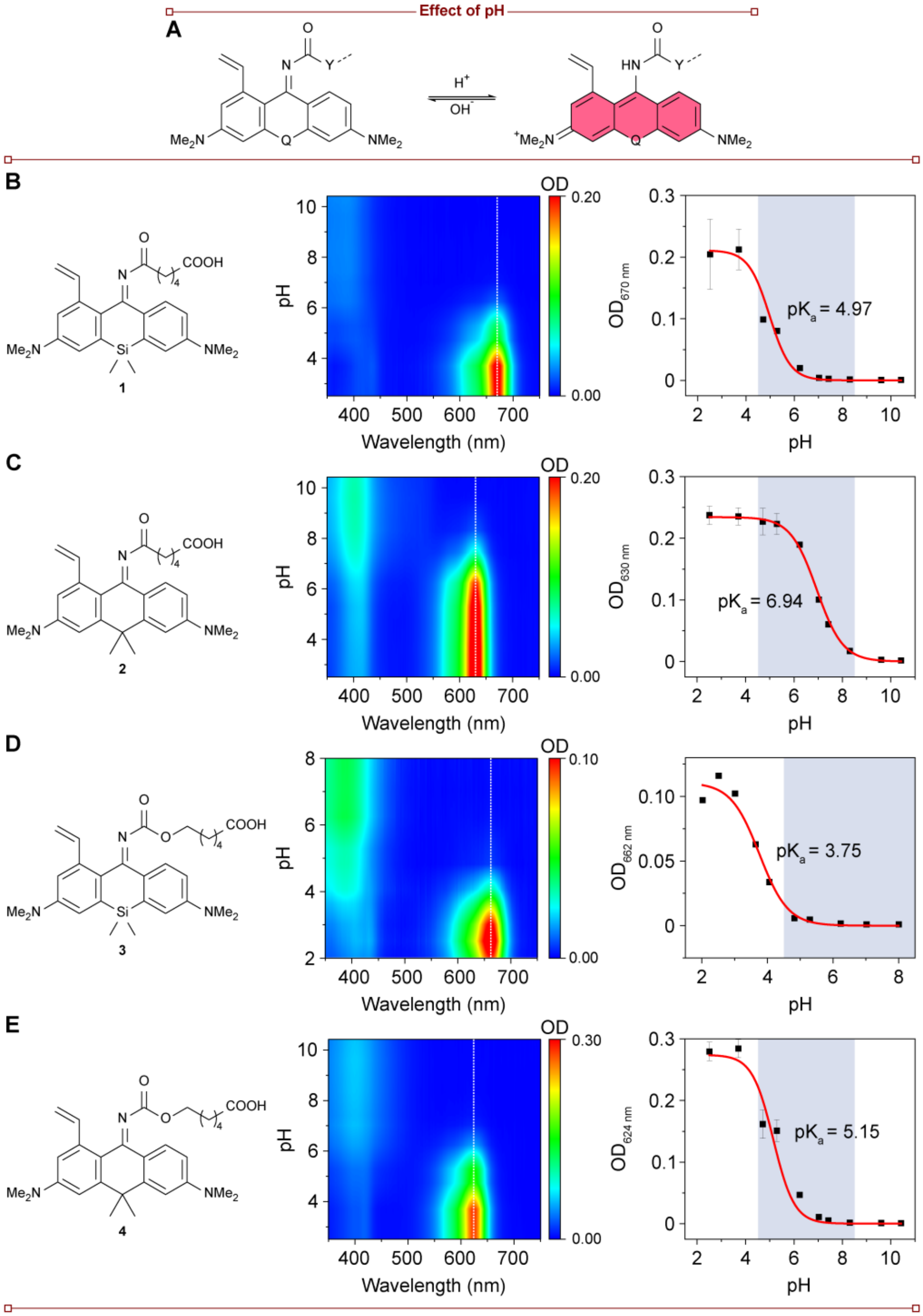

**Figure S1. Effect of pH on compounds 1–4.**

(A) Proposed change in structure of compounds **1**–**4** upon protonation.

(B–E) Chemical structure (left), absorption spectral changes of 5.0 μM solutions (center), and the determined pKa (right) for compounds **1**–**4** with the biologically relevant pH window (4.5–8.5) is show in gray.

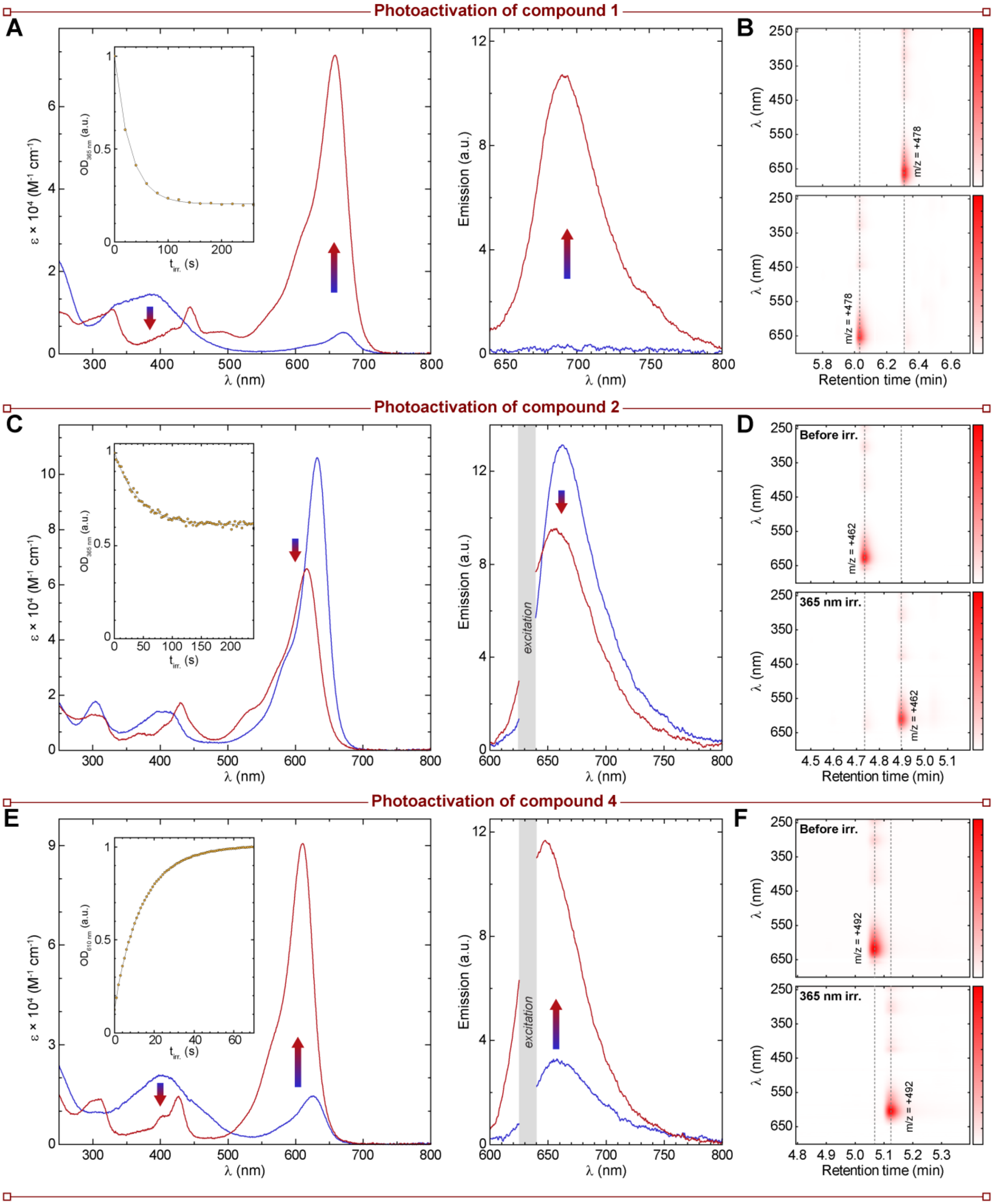
Photoactivation of compound 1
A
ε × 10⁴ (M⁻¹ cm⁻¹)
λ (nm)
OD365 nm (a.u.)
tirr. (s)
Emission (a.u.)
B
λ (nm)
m/z = +478
m/z = +478
Retention time (min)
Photoactivation of compound 2
C
ε × 10⁴ (M⁻¹ cm⁻¹)
λ (nm)
OD365 nm (a.u.)
tirr. (s)
Emission (a.u.)
excitation
D
Before irr.
365 nm irr.
λ (nm)
m/z = +462
m/z = +462
Retention time (min)
Photoactivation of compound 4
E
ε × 10⁴ (M⁻¹ cm⁻¹)
λ (nm)
OD610 nm (a.u.)
tirr. (s)
Emission (a.u.)
excitation
F
Before irr.
365 nm irr.
λ (nm)
m/z = +492
m/z = +492
Retention time (min)

**Figure S2. Photoactivation behavior.**

(A) Temporal evolution of the absorption (left) and fluorescence (right) spectra of **1** (**1** → **1-CF**; 5 μM) irradiated in phosphate buffer (100 mM, pH = 7; $\lambda_{act}$ = 365 nm).

(B) LC-MS chromatograms of the reaction mixtures of compound **1** before (top) and after (bottom) photoactivation with 365 nm irradiation.

(C) Temporal evolution of the absorption (left) and fluorescence (right) spectra of **2** (**2**→ **2-CF**; 5 μM) irradiated in phosphate buffer (100 mM, pH = 7; $\lambda_{act}$ = 365 nm).

(D) LC-MS chromatograms of the reaction mixtures of compound **2** before (top) and after (bottom) photoactivation with 365 nm irradiation.

(E) Temporal evolution of the absorption (left) and fluorescence (right) spectra of **4** (**4** → **4-CF**; 5 μM) irradiated in phosphate buffer (100 mM, pH = 7; $\lambda_{act}$ = 365 nm).

(F) LC-MS chromatograms of the reaction mixtures of compound **4** before (top) and after (bottom) photoactivation with 365 nm irradiation.

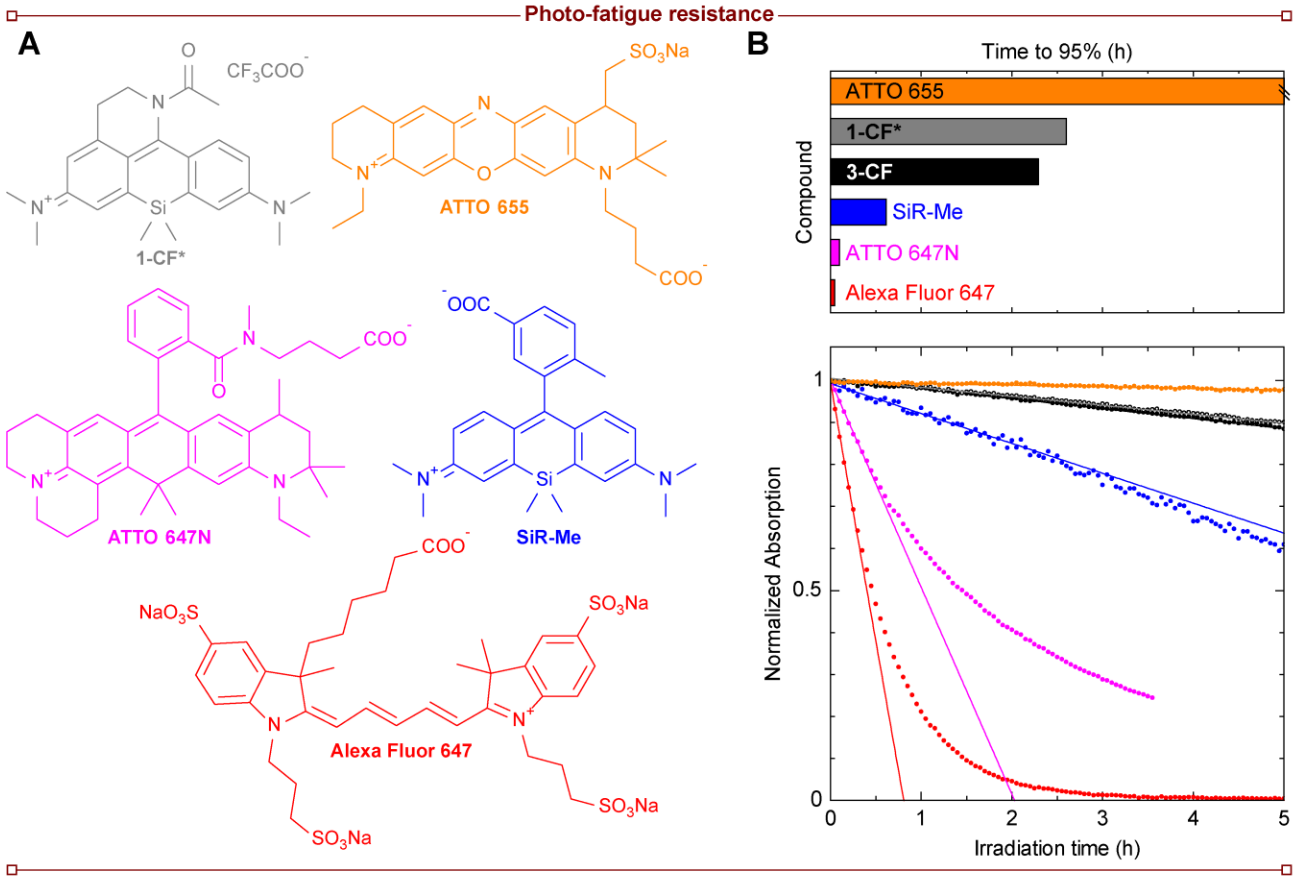

**Figure S3. Photo-fatigue resistance of closed form structures and established commercial fluorophores with similar spectral properties.**
(A) Structures of far-red fluorophores used.
(B) Photobleaching measurements performed in phosphate buffer ($\lambda_{exc}$ = 637 nm) for the tested compounds **1-CF*** (grey circles), **3-CF** (black circles), ATTO 655 (orange circles), ATTO 647N (magenta circles), SiR-Me (blue circles), and Alexa Fluor 647 (red circles). Solid lines denote the range of the linear fittings (5% decrease in signal) and the corresponding time in the top panel. For details on the determination of the quantum yield of bleaching ($\Phi_{bl}$) see reference [1].

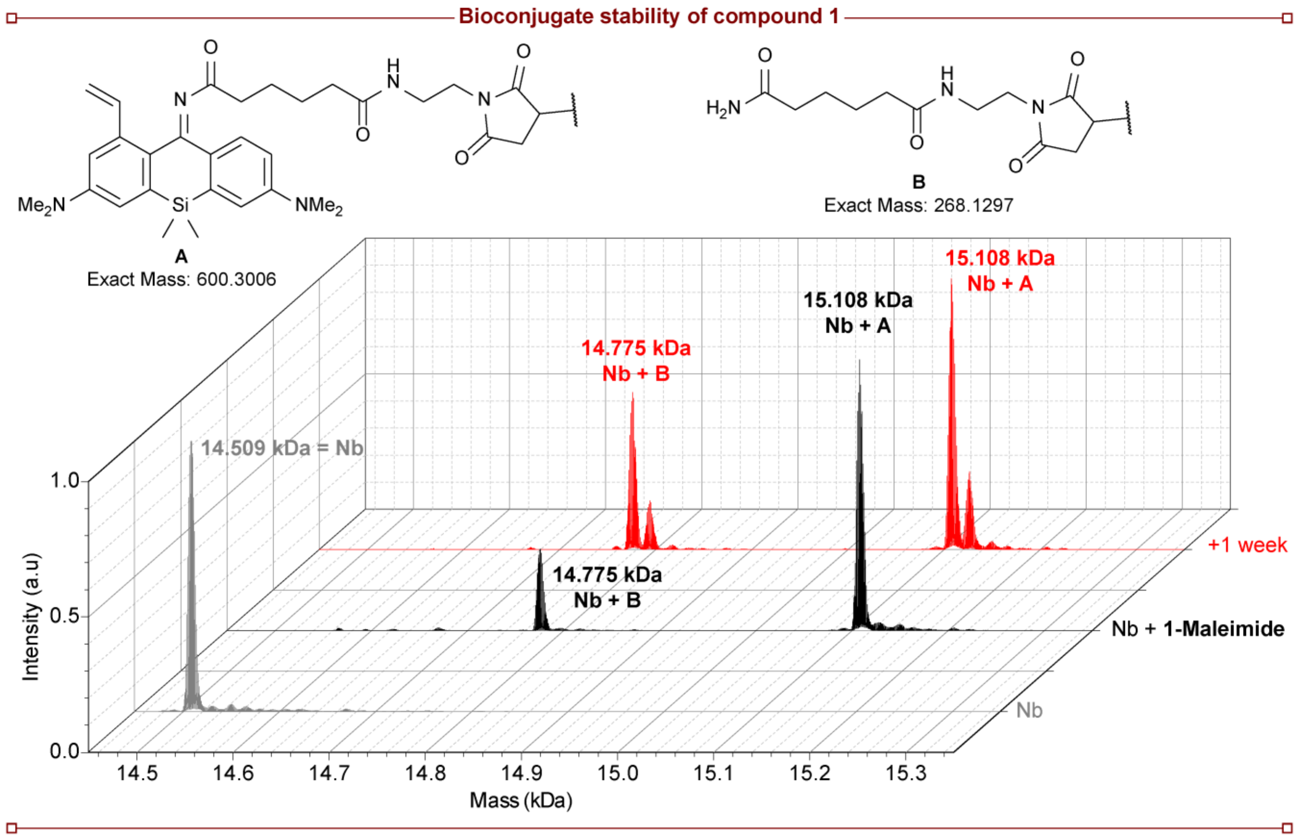

**Figure S4. Mass spec of nanobody-dye conjugate of compound 1-Maleimide.** Electrospray ionization mass spectrometry of FluoTag-Q anti-rabbit (clone 10E10), containing one cysteine residue. The spectra of the unlabeled nanobody (grey) and immediately after labelling (black) and after one week in storage at 4°C are presented, and the molecular mass of the dye-adduct (**A**) or linker fragment (**B**) are indicated.

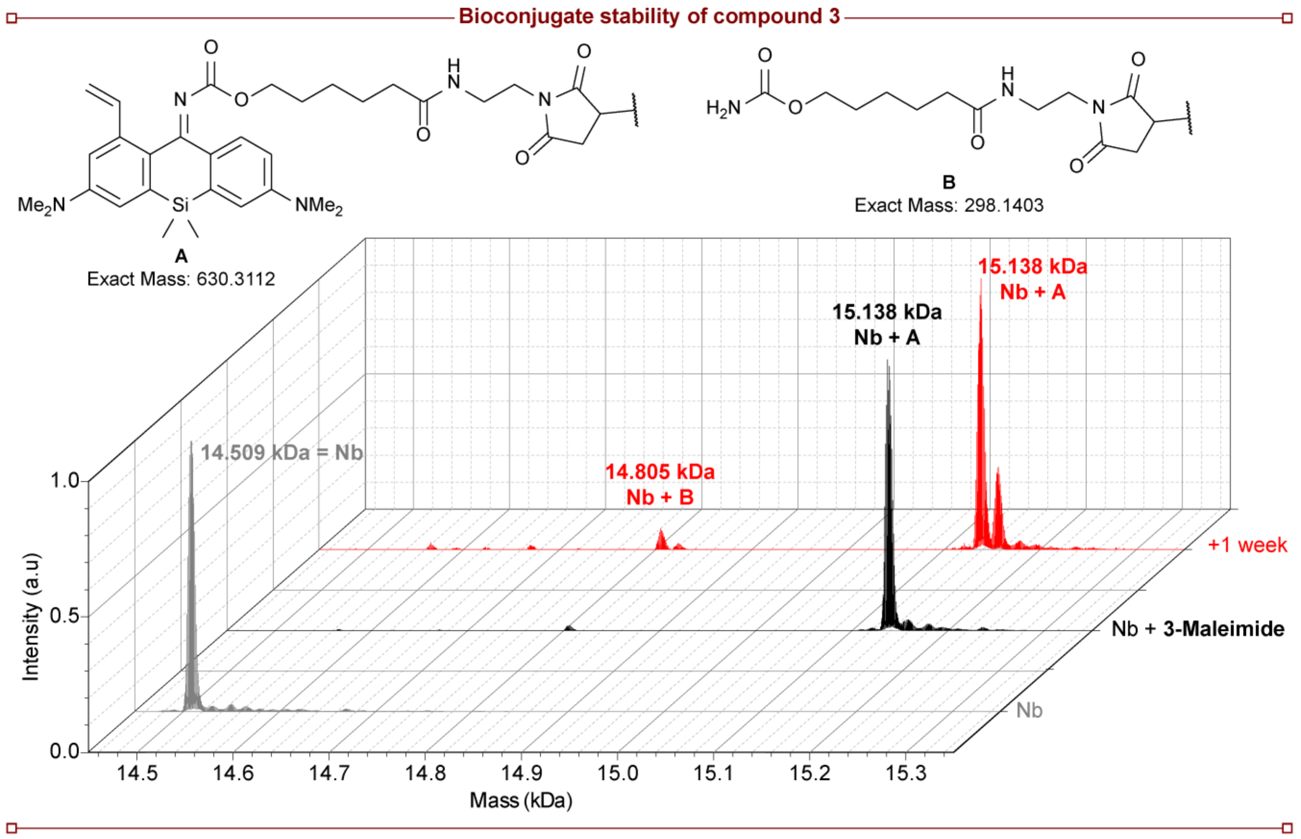

**Figure S5. Mass spec of nanobody-dye conjugate of compound 3-Maleimide.** Electrospray ionization mass spectrometry of FluoTag-Q anti-rabbit (clone 10E10), containing one cysteine residue. The spectra of the unlabeled nanobody (grey) and immediately after labelling (black) and after one week in storage at 4°C are presented, and the molecular mass of the dye-adduct (**A**) or linker fragment (**B**) are indicated.

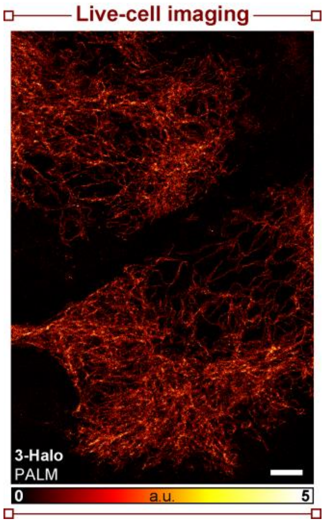

**Figure S6. Live-cell PALM imaging.** PALM image of live U-2 OS cells stably expressing a vimentin-HaloTag construct labelled with **3-Halo**. Scale bar: 5 µm.

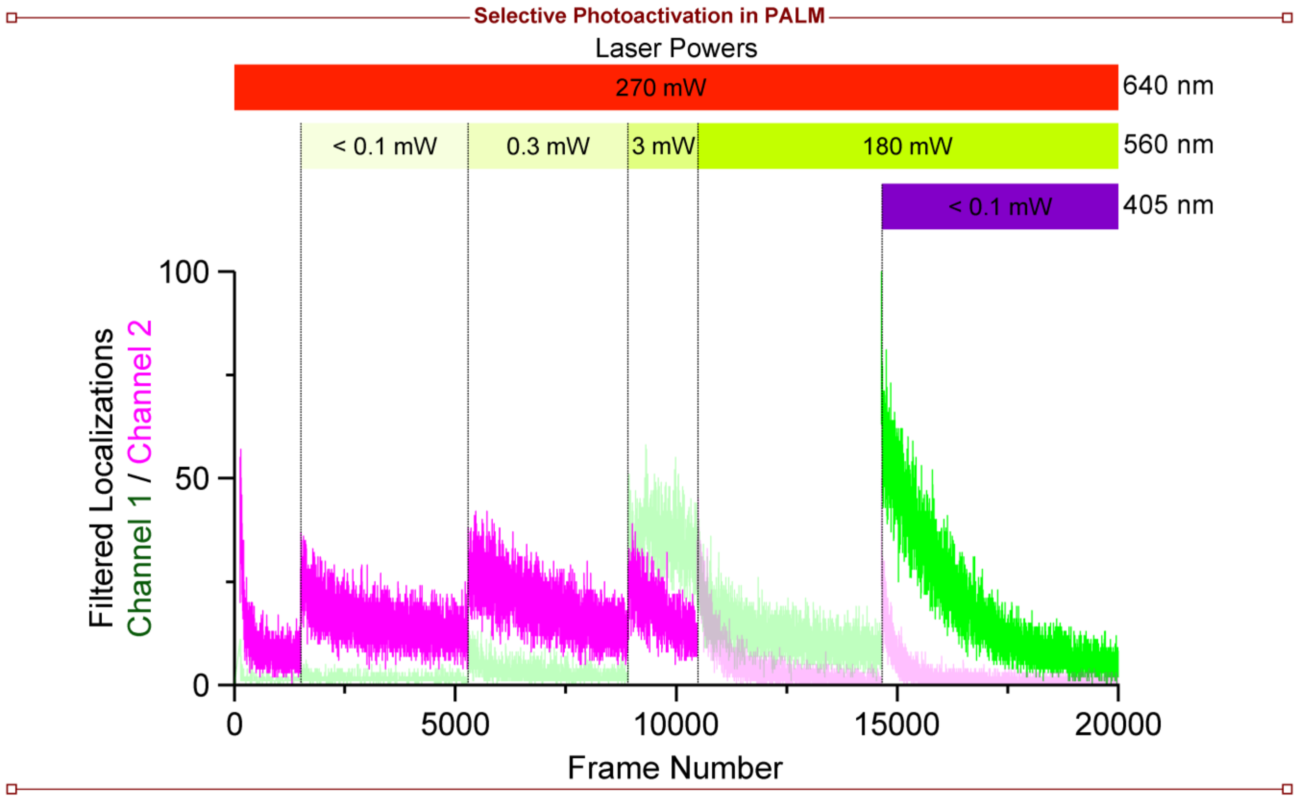

**Figure S7. Selective photoactivation and imaging of compound 3 and PaX$_{560}$.** Imaging sequence of 3D-PALM imaging shown in Figure 3K and the corresponding localizations obtained in each channel (after filtering). Localizations excluded from the rendering are shown as transparent lines.

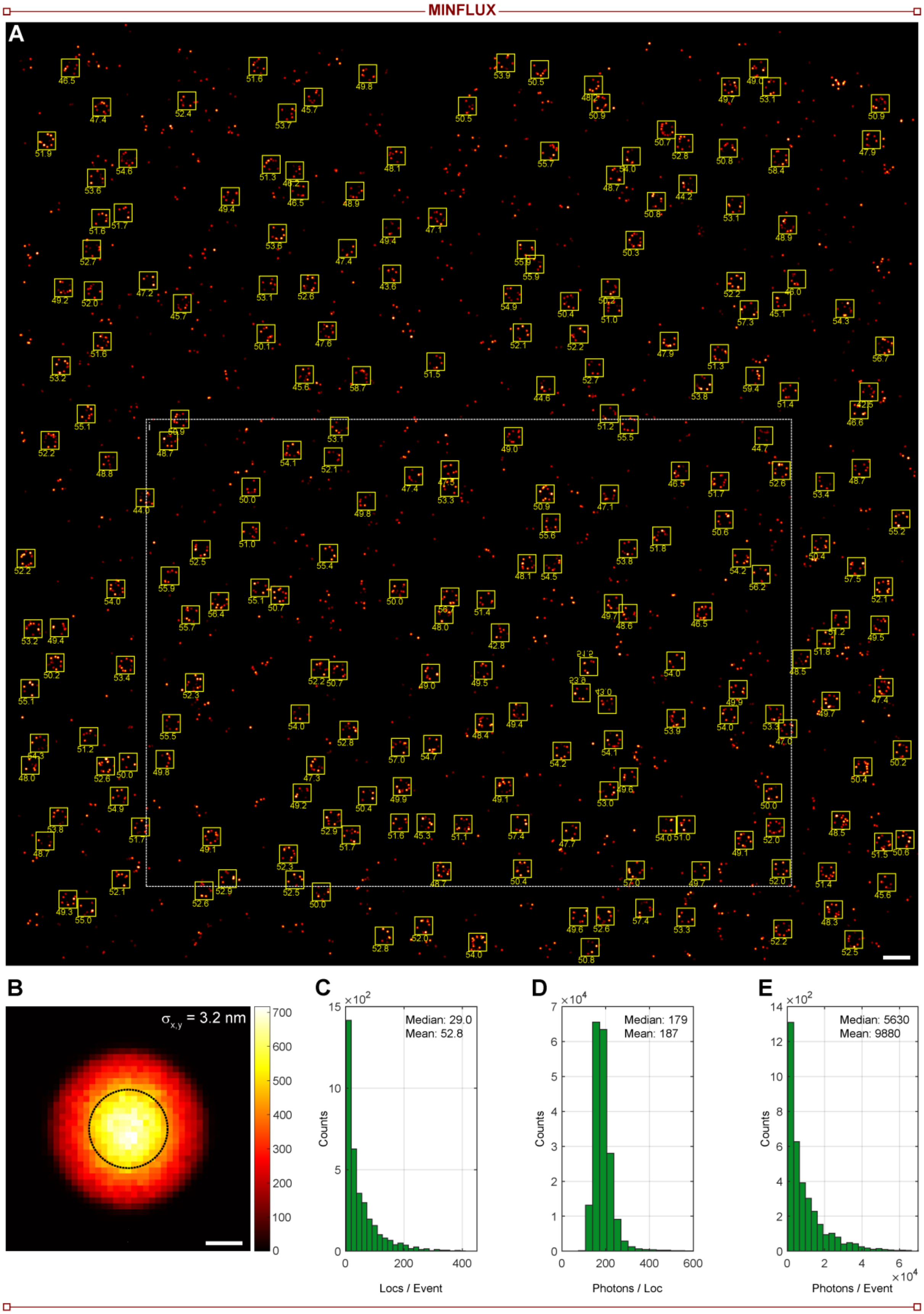
MINFLUX
A
B
$\sigma_{x,y}$ = 3.2 nm
700
600
500
400
300
200
100
0
C
×10^2
15
10
5
0
Counts
Median: 29.0
Mean: 52.8
0
200
400
Locs / Event
D
×10^4
7
6
5
4
3
2
1
0
Counts
Median: 179
Mean: 187
0
200
400
600
Photons / Loc
E
×10^2
14
12
10
8
6
4
2
0
Counts
Median: 5630
Mean: 9880
0
2
4
6
×10^4
Photons / Event

**Figure S8. 2D-MINFLUX image of NPCs labelled with compound 3-Halo.**

(A) 2D-MINFLUX image of U-2 OS cells stably expressing a NUP96-HaloTag construct labelled compound **3-Halo** (500 nM, overnight), fixed and mounted in Mowiol prior to imaging. Automatically selected structures used for the calculation of the occupancy histogram (Figure 4B) are marked as yellow boxes, and the corresponding radius (in nanometers) obtained by fitting the structure a circle function. i. Region shown in Figure 4. Scale bar: 500 nm.

(B) Histogram of the localization spread around their emitter centers Gaussian-fitted localization precision with $\sigma_{x,y}$ indicated with a circle for all localizations in A. Scale bar: 3 nm.

(C) Histogram of the number of localizations obtained from a single molecule 'on'-event for the molecules in A.

(D) Histogram of the number of photons obtained per localization for the molecules in A.

(E) Histogram of the number of photons obtained from a single molecule 'on'-event for the molecules in A.

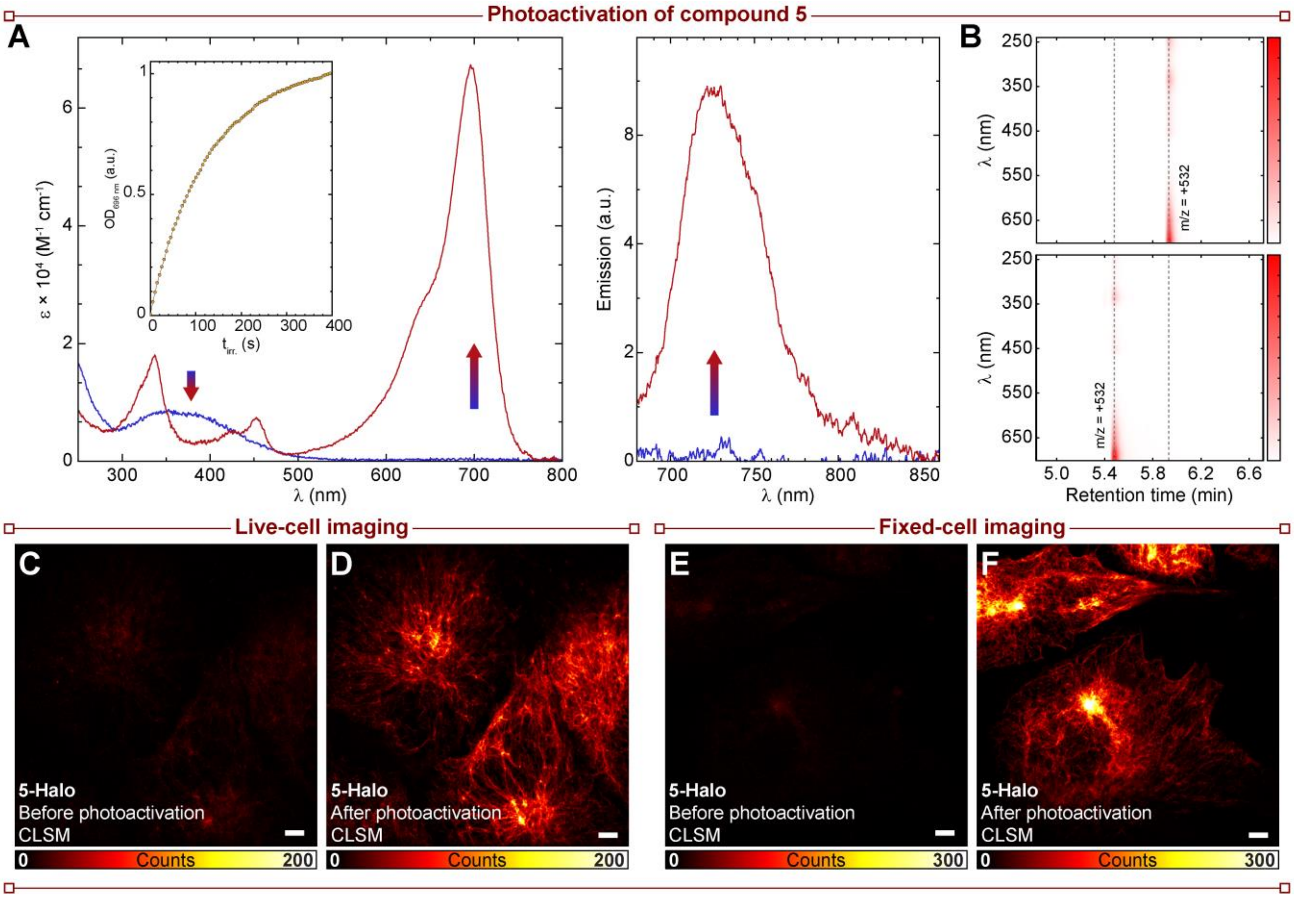

**Figure S9. Structure and characterization of compound 5.**

(A) Temporal evolution of the absorption (left) and fluorescence (right) spectra of **5** (**5** → **5-CF**; 5 μM) irradiated in phosphate buffer (100 mM, pH = 7; $\lambda_{act}$ = 365 nm).

(B) LC-MS chromatograms of the reaction mixtures of compound **5** before (top) and after (bottom) photoactivation with 365 nm irradiation.

(C) Confocal image of live U-2 OS cells stably expressing a vimentin-HaloTag construct labelled with **5-Halo** (500 nM) before photoactivation. Scale bar: 5 μm.

(D) Confocal images of the same sample as (D) following photoactivation with 405-nm laser. Scale bar: 5 μm.

(E) Confocal image of fixed U-2 OS cells stably expressing a vimentin-HaloTag construct labelled with **5-Halo** (500 nM) before photoactivation. Scale bar: 5 μm.

(F) Confocal images of the same sample as (D) following photoactivation with 405-nm laser. Scale bar: 5 μm.

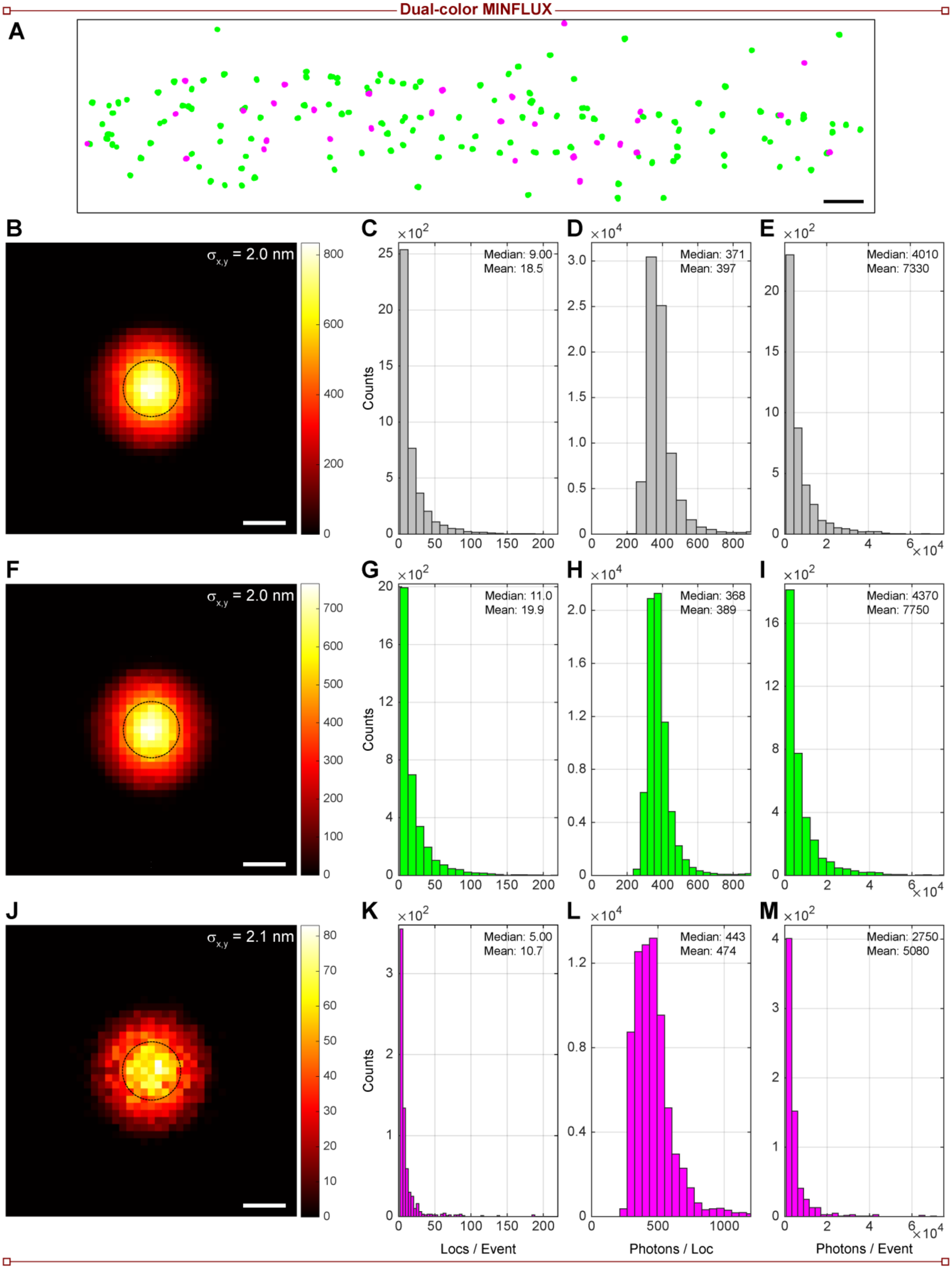
Dual-color MINFLUX
A
B
$\sigma_{x,y}$ = 2.0 nm
C
Median: 9.00
Mean: 18.5
D
Median: 371
Mean: 397
E
Median: 4010
Mean: 7330
F
$\sigma_{x,y}$ = 2.0 nm
G
Median: 11.0
Mean: 19.9
H
Median: 368
Mean: 389
I
Median: 4370
Mean: 7750
J
$\sigma_{x,y}$ = 2.1 nm
K
Median: 5.00
Mean: 10.7
L
Median: 443
Mean: 474
M
Median: 2750
Mean: 5080
Counts
Locs / Event
Photons / Loc
Photons / Event

**Figure S10. Dual-color MINFLUX imaging by spectral seperation.**

(A) Scatter plot of dual color 640 nm MINFLUX localizations of mitochondria proteins TOMM20 (green) and TIMM23 (magenta) labelled by secondary immunofluorescence with secondary nanobodies conjugated to compound **3-Maleimide** and **5-Maleimide** after color classification. Scale bar: 200 nm.

(B) Histogram of the localization spread around their emitter centers Gaussian-fitted localization precision with $\sigma_{x,y}$ indicated with a circle for all localizations before color classification. Scale bar: 3 nm.

(C) Histogram of the number of localizations obtained from a single molecule 'on'-event before color classification.

(D) Histogram of the number of photons obtained per localization before color classification.

(E) Histogram of the number of photons obtained from a single molecule 'on'-event before color classification.

(F) Histogram of the localization spread around their emitter centers Gaussian-fitted localization precision with $\sigma_{x,y}$ indicated with a circle for localizations classified as compound **3**. Scale bar: 3 nm.

(G) Histogram of the number of localizations obtained from a single molecule 'on'-event classified as compound **3**.

(H) Histogram of the number of photons obtained per localization classified as compound **3**.

(I) Histogram of the number of photons obtained from a single molecule 'on'-event classified as compound **3**.

(J) Histogram of the localization spread around their emitter centers Gaussian-fitted localization precision with $\sigma_{x,y}$ indicated with a circle for localizations classified as compound **5**. Scale bar: 3 nm.

(K) Histogram of the number of localizations obtained from a single molecule 'on'-event classified as compound **5**.

(L) Histogram of the number of photons obtained per localization classified as compound **5**.

(M) Histogram of the number of photons obtained from a single molecule 'on'-event classified as compound **5**.

### SUPPLEMENTARY METHODS

#### General Experimental Information and Synthesis

All chemical reagents (TCI, Sigma-Aldrich, Alfa Aesar) and dry solvents for synthesis (over molecular sieves, AcroSeal package, Acros Organics) were purchased from reputable suppliers and were used as received without further purification. The products were lyophilized from a suitable solvent system using Alpha 2-4 LDplus freeze-dryer (Martin Christ Gefriertrocknungsanlagen GmbH).

#### Thin Layer Chromatography

Normal phase TLC was performed on silica gel 60 $F_{254}$ (Merck Millipore, Germany). For TLC on reversed phase silica gel 60 RP-18 $F_{254}$s (Merck Millipore) was used. Compounds were detected by exposing TLC plates to UV-light (254 or 366 nm) or heating with vanillin stain (6 g vanillin and 1.5 mL conc. $H_2SO_4$ in 100 mL ethanol), unless indicated otherwise.

#### Flash Chromatography

Preparative flash chromatography was performed with an automated Isolera One system with Spektra package (Biotage AG) using commercially available cartridges of suitable size as indicated (RediSep Rf series from Teledyne ISCO, Puriflash Silica HP 30µm series from Interchim, Scorpius 30µm series from BGB Analytik).

#### Nuclear Magnetic Resonance (NMR)

NMR spectra were recorded on a Bruker DPX 400 spectrometer utilizing the Bruker Topspin 3.5 software. All spectra are referenced to tetramethylsilane as an internal standard (δ = 0.00 ppm). Multiplicities of the signals are described as follows: s = singlet, d = doublet, t = triplet, q = quartet, m = multiplet or overlap of non-equivalent resonances; br = broad signal. Coupling constants ${}^{n}J_{X\text{-}Y}$ are given in Hz, where n is the number of bonds between the coupled nuclei X and Y ($J_{H\text{-}H}$ are always listed as $J$ without indices).

#### Mass-Spectrometry (MS)

Low resolution mass spectra (100 - 1500 *m/z*) with electro-spray ionization (ESI) were obtained on a Shimadzu LC-MS system described below. High resolution mass spectra (HRMS) were obtained on a maXis II ETD (Bruker) with electrospray ionization (ESI) at the Mass Spectrometry Core facility of the Max-Planck Institute for Medical Research (Heidelberg, Germany).

#### High-Performance Liquid Chromatography (HPLC)

Analytical liquid chromatography-mass spectrometry was performed on an LC-MS system (Shimadzu, controlled with LabSolutions 5.89 software): 2x LC-20AD HPLC pumps with DGU-20A3R solvent degassing unit, SIL-20ACHT autosampler, CTO-20AC column oven, SPD-M30A diode array detector and CBM-20A communication bus module, integrated with CAMAG TLC-MS interface 2 and LCMS-2020 spectrometer with electrospray ionization (ESI, 100 – 1500 *m/z*). Analytical column: Hypersil GOLD 50×2.1 mm 1.9µm, standard conditions: sample volume 1-2 µL, solvent flow rate 0.5 mL/min, column temperature 30 °C. General method: isocratic 95:5 A:B over 2 min, then gradient 95:5 – 0:100 A:B over 5 min, then isocratic 0:100 A:B over 2 min; solvent A = water + 0.1% v/v $HCO_2H$, solvent B = acetonitrile + 0.1% v/v $HCO_2H$.

Preparative high-performance liquid chromatography was performed on a Büchi Reveleris Prep system using the suitable preparative columns and conditions as indicated for individual preparations. Method scouting was performed on a HPLC system (Shimadzu, controlled with LabSolutions 5.89 software): 2x LC-20AD HPLC pumps with DGU-20A3R solvent degassing unit, CTO-20AC column oven equipped with a manual injector with a 20 µL sample loop, SPD-M20A diode array detector, RF-20A fluorescence detector and CBM-20A communication bus module; or on a Dionex Ultimate 3000 UPLC system: LPG-3400SD pump, WPS-3000SL autosampler, TCC-3000SD column compartment with 2× 7-port 6-position valves and DAD-3000RS diode array detector. The test runs were performed on analytical columns with matching phases (HPLC: Interchim 250×4.6 mm 10 µm C18HQ, Interchim 250×4.6 mm 5 µm PhC4, solvent flow rate 1.2 mL/min; UPLC: Interchim C18HQ or PhC4 75×2.1 mm 2.2 µm, ThermoFisher Hypersil GOLD 100×2.1 mm 1.9 µm, solvent flow rate 0.5 mL/min).

**Determination of pH-dependent behavior by absorption spectroscopy**

A series of buffers ranging from pH 2.5 to 10.5 (or pH 2 to 8) was prepared from PBS adjusted with HCl or NaOH (3 M and 0.3 M) and monitoring with a PT-10 pH-meter (Sartorius AG, Göttingen, Germany). Using these buffers, 5 µM solutions of the dyes (**1**–**4**) were prepared in 96-well µ-plates with glass bottoms (ibidi GmbH, Gräfelfing, Germany) containing 10% (v/v) DMSO. Each pH condition was prepared in triplicate, and the corresponding pH buffer containing 10% (v/v) DMSO was used as the respective blank. Before measurement, the plate was equilibrated at 25 °C for 10 min. The absorption spectral measurements were performed in a plate reader (CLARIOstar® Plus, BMG Labtech, Germany) operated with CLARIOstar® software (5.70 R2). Ten repetitive measurements were performed and averaged to reduce instrument noise. Before each repetition, each well was scanned with 22 flashes. The obtained absorption spectra were further processed in MARS software (3.42 R5; BMG Labtech), analyzed and plotted with OriginPro 2019 (9.6.0.172, OriginLab Corporation, USA).

**Optical spectroscopy**

Fluorescence quantum yield measurements were measured Quantaurus-QY absolute PL quantum yield spectrometer (C11347-11, Hamamatsu) in methanol and phosphate buffer (pH = 7). Fluorescence lifetimes were measured with a FluoTime 300 fluorescence lifetime spectrometer (PicoQuant, controlled with the EasyTau1.4 software) in 3 mL quartz cells (optical path length 1 cm, model 119F-10-40 or model 111-10-40, Hellma Analytics). All measurements were performed in air-saturated solvents at ambient temperature.

**Photolysis of compounds and chemometric analysis of the photoactivation and photobleaching reaction kinetics**

Photolysis experiments were performed as previously described.[1] In brief, solutions in phosphate buffer (100 mM, pH = 7.0; 5.0 µM dye) or methanol were irradiated in a previously described[2] home-built setup with a 405 nm LED source (M405L3, Thorlabs Inc.) in combination with a bandpass (10 nm) filter (FB405-10, Thorlabs Inc.). During the irradiation, samples were maintained at 20 °C and continuously stirred with a Peltier-based temperature-controlled cuvette holder (Luma 40, Quantum Northwest, Inc.). The absorption and emission of irradiated solutions was monitored at desired irradiation intervals with a fiber-based spectrometer (Flame-S-UV-Vis-ES, Ocean Insight). For absorption measurements, a deuterium and tungsten halogen source was used for illumination (DH-2000-BAL, Ocean Insight), and for fluorescence excitation was performed in a 90° configuration with a red LED source (M625L3, Thorlabs Inc.) in combination with a bandpass filter centered at 632.8 nm, with a FWHM of 1 nm (FL632.8-1, Thorlabs Inc.). Data collection and analysis was performed with custom-made routines in Matlab. Samples for LCMS or ESI-MS analysis were taken before and after photolysis was performed. Photobleaching reactions (Figure **S3**) were studied in the same setup using a 637 nm LED source (M625L3, Thorlabs Inc.), without a bandpass filter. The Nominal Wavelength of the LED is 625 nm (informed by the vendor), but the peak wavelength measured for our particular LED is 637 nm, with a FWHM ≈ 10 nm. Photobleaching quantum yields were calculated as previously described.[1]

**Antibodies, Nanobodies and Other Fluorescent Conjugates**

**Table S1.** Antibodies and Nanobodies used.

| Reagent | Type | Target | Host | Supplier | Catalogue No. | Dilution |
|---|---|---|---|---|---|---|
| sdAb anti-Rabbit IgG, unconjugated | Nanobody | Rabbit | Camelid | NanoTag Biotechnologies | N2402 | 1:2000 |
| sdAb anti-Rabbit IgG, unconjugated | Nanobody | Rabbit | Camelid | NanoTag Biotechnologies | N2403 | 1:2000 |
| sdAb anti-Rabbit IgG, unconjugated | Nanobody | Rabbit | Camelid | NanoTag Biotechnologies | N2405 | |
| Anti-TOMM20 antibody [EPR15581-54] | Primary Antibody (monoclonal) | TOMM20 | Rabbit | Abcam | ab186735 | 1:250 |
| Anti-Tim23 antibody | Primary Antibody (monoclonal) | TIMM23 | Mouse | BD Biosciences | 611222 | 1:100 |

**Conjugation of nanobodies via thiol groups**

Unconjugated nanobodies (NanoTag Biotechnologies product nos.: N2402, N2403, N2405) with site-specific cysteine residues) were reconstituted and labelled according to manufacturer's protocols. In brief, then the pH of the solution was adjusted by the addition of 1/10 volume of 1 M Tris/HCl buffer (pH 8.0), and 1.5- to 3-fold molar excess (in respect to the number of cysteines per nanobody) of maleimide-derivative was immediately added. The reaction mixture was vortexed, overlaid with argon and incubated in the dark, on ice for 1.5 h.

The unreacted excess dye was removed, and the buffer was exchanged to PBS using a desalting column (7K MWCO Zeba Spin Desalting Column, Thermo Scientific). The DOL value of the resulting conjugate was determined by ESI-MS.

Antibodies and nanobodies were used without further validation as the obtained labelling was clearly compatible with the expected structures.

**Cell culture**

For cultivation of the genetically engineered human bone osteosarcoma epithelial cell lines U-2 OS-CRISPR-NUP96-Halo clone #252 (300448, CLS GmbH) [3] McCoy's 5a (Modified) Medium containing L-glutamine (26600023, Gibco) was supplemented with 2 mM sodium pyruvate (11360070, Gibco), 1% (v/v) penicillin/streptomycin (15140122, Gibco), 0% or 10% (v/v) minimum essential medium (11140035, Gibco) and 10% (v/v) fetal bovine serum (FBS; 10500064, Gibco). U-2 OS-Vim-Halo[4,5] and HeLa COX8A-SNAP-tag[6] cells were cultured in Dulbecco's Modified Eagle Medium (DMEM, 4.5 g/L glucose) containing GlutaMAX and sodium pyruvate (31966-021, Gibco), supplemented with 1% (v/v) penicillin/streptomycin (15140122, Gibco) and 10% (v/v) FBS (10500064, Gibco). Cells were grown at 37 °C in humidified air with 5% CO2 and were harvested using TrypLE Express Enzym (1x) (12604013, Gibco; <20 passages between thawing and experimental use). Cell line authentication and mycoplasma testing is regularly performed. For sample preparation, $<10^5$ cells/well were seeded on #1.5 glass coverslips in 12-well cell culture plates and kept for 12-72 h at 37 °C and 5% $CO_2$.

**Live-cell labelling for optical microscopy and nanoscopy of cells**

The entire sample preparation procedure was conducted under red light (generic 12 V red LED strips, IP65 waterproof, 620-640 nm). Stocks solutions of SNAP-tag ligand (**3-BG**) or HaloTag ligand derivatives of the compounds (**3-Halo**, **5-Halo**) were prepared in DMSO (500 µM – 5 mM). U-2 OS cells that stably expressed Vimentin-HaloTag,[4,5], NUP96-HaloTag,[3] or HeLa cells that stably expressed COX8A-SNAP-tag[6] were grown for 12-72 h on glass coverslips. Cells were incubated in the dark for 30 min to overnight at 37 °C and 5% $CO_2$ (depending on the dye and experiment) with the respective fluorescent ligands diluted from DMSO stock solutions with Fluorobrite (A1896701, Gibco) supplemented with 10% (v/v) FBS (10500064, ThermoFisher), 2% (v/v) GlutaMAX (35050061, Gibco) and 1% (v/v) penicillin/streptomycin (labeling medium) to a final concentration of 500 nm – 1 µM. After labeling with dyes, the samples were protected from the ambient light. Cells were washed twice with cell labeling medium for ca. 15–30 minutes; then the medium was changed for fresh media for live-cell imaging or fixed as described below.

For live-cell imaging (Figs. 3C, 3D, 3E, 3F, 3G, S6) cells were mounted in a live-cell chamber (CM-B18-1, Live Cell Instrument Co.) with labeling medium.

**Fixation and immunofluorescence labelling**

PFA fixation for preservation of vimentin filaments (Figs. 3H, 3I, 4C, 4D) was performed with a 4% formaldehyde solution in PBS (pH 7.4) at room temperature for 20 min, then washed 5 min with PBS.

PFA fixation for preservation of nuclear pore complexes (Figs. 3J, 4a) was performed according to previous procedures with minor modifications.[3] In brief, cells were prefixed with 2.4% formaldehyde solution in PBS at room temperature for 30 seconds, permeabilized with 0.4% Triton X-100 in PBS at room temperature for 3 minutes, and fixed with a 2.4% formaldehyde solution in PBS at room temperature for 30 min, then rinsed with PBS.

PFA fixation for preservation of mitochondria cristae (Fig. 3K) was performed with a warm 8% formaldehyde solution in PBS at 37 degrees for 7 minutes, and permeabilized with 0.5% Triton X-100 in PBS at room temperature for 5 min. To reduce unspecific binding blocking buffer (5% BSA in PBS) was added and incubated for 10 minutes at room temperature, then washed with PBS. The coverslips were overlaid with the primary antibody solution for TOMM20 (from rabbit) in blocking buffer and incubated in a humid chamber for 1 h at room temperature and then washed with blocking buffer (3×5 min). The coverslips were then incubated with the secondary anti-rabbit nanobody labelled with **$PaX_{560}$-Maleimide**[1] in blocking buffer, in a humid chamber for 1 h at room temperature, and then washed with blocking buffer (3×5 min), and with PBS (3×5 min).

Fixation for dual-color MINFLUX imaging of mitochondria (Fig. 5B–D) was performed with a 4% formaldehyde solution containing 0.2% gluteraldehyde in PBS at 37 degrees for 10 min. Samples were incubated with a quenching solution (0.1 % sodium borohydride in PBS) for 7 min at room temperature then washed twice with PBS. To permeabilize and reduce unspecific binding, a blocking buffer (5% BSA in PBS with 0.1% Triton X-100) was added and incubated for 30 minutes at room temperature.

The coverslips were overlaid with the primary antibodies for TOMM20 (from rabbit) and TIMM23 (from mouse) in blocking buffer (5% BSA in PBS with 0.1% Triton X-100) and incubated in a humid chamber for 1 h at room temperature and then washed with PBS (3×5 min). The coverslips were then incubated with the secondary nanobodies labelled with **3-Maleimide** (anti-rabbit) and **5-Maleimide** (anti-mouse) in blocking buffer, in a humid chamber for 1 h at room temperature, and then washed with PBS (3×5 min). Samples were post-fixed with a 4% PFA solution for 10 min at room temperature, then washed with PBS (3×5 min).

For confocal and PALM imaging, fixed cells were mounted in a live-cell chamber with PBS.

For MINFLUX imaging, gold nanoparticle fiducials were added to the sample (BBI Solutions, cat. SKU:EM.GC150-7, for 5 min), before mounting in either Mowiol or PBS and sealed with silicone (Picodent-ecosil, cat 1307100).

### Confocal and STED (stimulated emission depletion) microscopy

Confocal and STED images were acquired using two Abberior Expert Line (Abberior Instruments GmbH, Göttingen, Germany) fluorescence microscopes built on a motorized inverted microscope IX83 (Olympus, Tokyo, Japan). Microscope 1 is equipped with pulsed STED lasers at 595 nm and 775 nm shaped by Spatial Light Modulators (SLMs), and with 355 nm, 405 nm, 485 nm, 561 nm, and 640 nm excitation lasers, and a 100x/1.40 oil immersion objective lenses (Olympus). Microscope 2 is equipped with pulsed STED lasers at 655 nm and 775 nm, and with 520 nm, 561 nm, 640 nm, and multiphoton (Chameleon Vision II, Coherent, Santa Clara, USA) excitation lasers, and a 60x/1.42 oil immersion objective lens (Olympus). The multiphoton laser is tuneable in the 680 nm – 1080 nm range. Spectral detection is performed in both cases with avalanche photodiodes at spectral windows adjusted for each particular fluorophore.

Imaging and image processing was done with ImSpector software (v. 16.3.13367; Abberior Instruments GmbH, Göttingen, Germany), and all images are displayed as raw data unless otherwise noted.

### Superresolution single molecule localization microscopy (SMLM) / Photoactivated localization microscopy (PALM)

SMLM / PALM images was acquired using an ONI Nanoimager V3 (Oxford Nanoimaging, Oxford, UK). The 405 nm activation laser was applied as CW illumination.

Particular imaging conditions are given in **Table S2**.

**Table S2.** PALM imaging parameters.

| Figure | Compound | Exposure Time (ms) | Excitation | | Mean Uncertainty (nm) | Total Frames | Mean $N_{photons}$ Per Localization |
|---|---|---|---|---|---|---|---|
| | | | Laser (nm) | Power (mW) | | | |
| Fig. 3H | 3-Halo | 20 | 640 nm | 270 | 18.0 | 15k | 4900 |
| Fig. 3J | 3-Halo | 20 | 640 nm | 270 | 15.8 | 4.3k | 4400 |
| Fig. 3K | 3-BG<br>$PaX_{560}$ | 20 | 640 nm<br>560 nm | 270<br>180 | -<br>- | -<br>- | -<br>- |
| Supp. Fig. S6 | 3-Halo | 50 | 640 nm | 120 | 17.4 | 2.2k | 2100 |

2D images (Figures 3H, 3J, S6) were analyzed and processed using the ThunderSTORM plugin[2] on ImageJ (version 1.52p). In brief, images were filtered with a wavelet filer (B-spline order 3, scale 2.0), approximate localization of the molecules was performed with a local maximum method (peak intensity threshold of ca. 1.4–1.6 standard deviations; connectivity 8-neighbourhood), and sub-pixel localization of the molecules was performed with a maximum likelihood fitting method (PSF integrated method, fitting radius of 3 pixels, initial sigma 1.6). Post-processing was performed with the same plug-in. Data was drift-corrected based on the cross-correlation method, merged within the size of 50 nm (0.42 pixel) with 0 off-frames allowed. Sigma values were filtered to converge to a normal Gaussian distribution function (80 < sigma < 200). Photon numbers and uncertainty values were restricted (to 300 < intensity < 50000 and uncertainty < 30) to filter any outliers. Final images were produced using normalized Gaussian rendered with a fixed sigma corresponding to the mean localization uncertainties, and a pixel-size of 10 nm.

The 3D image (Figure 3K) was analyzed and processed using the ONI Nanoimager™ Software, Development build: Apr 9 2023 22:54:56 Version: 1.19.7.20230409223555 - 28f00b5. Data was drift-corrected, and the data filtered according to **Table S3**. Final images were produced with a fixed gaussian width of 10 nm and a pixel-size of 5 nm.

**Table S3.** Filter conditions for Figure 3K.

| | Channel 0 (580 – 620nm) | | Channel 1 (662 – 710 nm) | |
|---|---|---|---|---|
| | **Min** | **Max** | **Min** | **Max** |
| **Photon Count** | 300 | 50000 | 300 | 50000 |
| **Z Position** | -1000 | 1000 | -1000 | 1000 |
| **Localization Precision (X)** | 0 | 20 | 0 | 20 |
| **Localization Precision (Y)** | 0 | 20 | 0 | 20 |
| **Sigma (X)** | 70 | 300 | 80 | 300 |
| **Sigma (Y)** | 70 | 300 | 80 | 300 |
| **p-Value** | 0 | 1 | 0 | 1 |
| **Frame Index** | 14650 | 50000 | 10 | 10500 |

**MINFLUX**

MINFLUX Imaging

For MINFLUX imaging an Abberior 3D MINFLUX microscope was used. Details of the instrument are described in Schmidt et al., 2021. The microscope, built on an Olympus IX83 body with a 60x UPLXAPO60XO oil objective, was equipped with a 640 nm excitation laser, a 405 nm activation laser, 488 nm and 560 nm confocal lasers, a 980 nm stabilization laser and an xyz piezo stage (Piezoconcept) for active sample stabilization.

For acquisition the pinhole was set to 0.83 AU and signal was detected on APDs in the spectral window of Cy5 (650-720 nm). The excitation power was set to initially 33 µW in the sample position for 2D imaging and to 56 µW for 3D imaging. The activation 405 nm laser was attenuated with an additional ND2 filter. Activation was switched on and the power was gradually increased up to 1.1 µW, sustaining the frequency of detected events, until the events became sparse in time and the imaging was stopped.

MINFLUX Sequences

MINFLUX imaging was performed with modified imaging sequences based on the standard imaging sequences provided by the manufacturer. For both, 2D and 3D imaging, the stickiness was globally increased to 5 for more robustness. Iteration specific sequence settings are listed in **Tables S4** and **S5**.

**Table S4.** Iteration specific sequence settings for 2D MINFLUX imaging

| Iteration | Background threshold [Hz] | CFR limit | Dwell time [ms] | L [nm] | Minimum photon count | Laser power factor |
|---|---|---|---|---|---|---|
| **1** | 15000 | 2.0 | 1 | 288 | 50 | 1 |
| **2** | 10000 | -1.0 | 1 | 288 | 50 | 1 |
| **3** | 10000 | 0.8 | 1 | 151 | 50 | 2 |
| **4** | 10000 | 0.8 | 1 | 76 | 50 | 4 |
| **5** | 10000 | 2.0 | 1 | 40 | 50 | 6 |

**Table S5.** Iteration specific sequence settings for 3D MINFLUX imaging

| Iteration | Background threshold [Hz] | CFR limit | Dwell time [ms] | L [nm] | Minimum photon count | Laser power factor |
|---|---|---|---|---|---|---|
| **1 (xy)** | 15000 | - | 1 | 288 | 50 | 1 |
| **2 (z)** | 15000 | - | 1 | 288 | 400 | 1 |
| **3 (xy)** | 10000 | - | 1 | 288 | 50 | 1 |
| **4 (z)** | 10000 | - | 1 | 288 | 50 | 1 |
| **5 (xy)** | 10000 | 0.9 | 1 | 151 | 50 | 2 |
| **6 (z)** | 10000 | - | 1 | 151 | 33 | 2 |
| **7 (xy)** | 10000 | 0.8 | 1 | 76 | 50 | 4 |
| **8 (z)** | 10000 | - | 1 | 76 | 33 | 4 |
| **9 (xy)** | 10000 | - | 1 | 40 | 50 | 6 |
| **10 (z)** | 10000 | - | 1 | 40 | 50 | 6 |

MINFLUX Data Analysis

Data analysis and image rendering was performed using dedicated MATLAB routines as previously described.[1,7] A first filter was applied to the list of localizations using the MATLAB implementation of the density-based clustering algorithm dbscan (epsilon = 9 nm, minPts = 3). A second filter was applied to include only molecules which provided at least 2 localizations, and to discard localizations outside a radius of 6.6 nm around their mean position. The final image was produced using an amplitude-normalized Gaussian rendering method from the list of filtered localizations, with a fixed sigma value of 3 nm (corresponding approximately to their average localization uncertainty) and a pixel-size of 1 nm for the rendered image. For ease of visualization, the images (Figure 4A and S8A)are displayed with a colormap with a nonlinear (gamma correction with A = 1 and $\gamma$ = 0.5) brightness progression.

For the 3D rendering of vimentin filaments (**Figure 4C**–**D**), the 3D stack was generated from the localization positions with Gaussian rendering, using a voxel size of 2×2×2 nm and a Gaussian blurring with a full width at half maximum of 4 nm. A gamma correction ($\gamma$ = 0.33) was applied in FIJI[8] before visualizing with the 3DScript plugin.[9]

NPC Occupancy Analysis

The occupancy (Occ) value for nuclear pore complexes was obtained using dedicated routines previously described.[3,7,10,11] The MINLUX image was automatically segmented for NPCs structures by a cross correlation of the image (binarized) with a pattern structure generated from ring-like structure (radius = 50 nm) convoluted with a Gaussian (FWHM = 10 nm) to select NUP structures. The center position of each ROI (150×150 px$^2$) was selected from hot spots, using a threshold of 0.55 (Figure S8A). ROIs closer than 140 nm were eliminated (the one with lowest correlation value) to avoid counting the same structure twice. Within a single ROI (a single NPC), all MINFLUX localizations from the same single molecule 'on'-event were combined fitted to a circle function of variable center and radius (mean fitted radius obtained = 51.16 nm). Then, the center coordinates of the fitted circle were subtracted from localization coordinates, and outliers were eliminated (localizations with a radius, >70 nm away from the origin, were eliminated). Finally, we constructed an 8-bin histogram of the angular positions, rotated such that a maximum number of positions fell into a single bin, to avoid that a bin edge divided an NPC corner in half. Bins with at least one 'on'-event were considered as occupied, and occupancy (*Occ*) histograms were computed based on this information (**Figure 4B**).

To infer the effective labeling efficiency (*ELE*), the obtained occupancy histograms were fitted using a binomial probabilistic model, assuming an eightfold symmetry of the NPC with four labeling sites on each corner (two copies per corner and per cytoplasmic/nucleoplasmic plane). Under this assumption, the probability of observing at least one localization in *Occ* corners, with $Occ = 0 - 8$ as the measured occupancy, is described by:

$$p(Occ) = \binom{8}{Occ}(1-(1-ELE)^4)^{\mathrm{Occ}}\left(1-(1-(1-ELE)^4)\right)^{(8-Occ)} \qquad \textbf{Equation S1}$$

with the first term being the binomial coefficient:

$$\binom{8}{Occ} = \frac{8!}{(8-Occ)!\ Occ!} \qquad \textbf{Equation S2}$$

We note that our segmenting process may discard individual NPCs with a low number of imaged corners (*Occ* < 3), as they could not be clearly delineated against unspecific background localizations.[7,12] The occupancy histograms were normalized and fitted with the model in **Equation S1**, with *ELE* as the only variable parameter, using a dedicated MATLAB routine based on the fitting function *lsqcurvefit* with 0.5 as the initial value. The statistical error of *ELE* was estimated by bootstrapping with 10000 randomly drawn pseudo data sets.

Dual Color MINFLUX Data Analysis

In the first step, dual color MINFLUX data were treated in a similar way to the single-color data, with adapted parameters for the applied filters.

A first filter was applied to the list of localizations using the MATLAB implementation of the density-based clustering algorithm dbscan (epsilon = 6.6 nm, minPts = 3). A second filter was applied to include only molecules which provided at least 2 localizations, and to discard localizations outside a radius of 4.4 nm around their mean position. Subsequently, spectral classification was applied. All filtered localizations of the same DBSCAN cluster were considered to belong to the same single molecule and assigned a color based on their mean value ($DCR_{cluster}$ = mean($DCR_{loc}$)) of the detector channel ratio (*DCR*, fraction of photons on the green detector). Clusters with $DCR_{cluster}$ in the 0.00-0.4 range were assigned to the red channel ($PaX_{695}$, **Figure 5C**), and clusters in the 0.4-1.00 range were assigned to the green one ($PaX_{650}$, **Figure 5B**).. It must be noted that individual values of $DCR_{loc}$ fall outside of the corresponding selected range for $DCR_{cluster}$, and thus the $DCR_{loc}$ distributions after spectral classification (**Figure 5E**) are not truncated at *DCR* = 0.4. Images (**Figure 5B–D**) were generated using an amplitude-normalized Gaussian rendering method with a fixed sigma value of 2 nm and a pixel size of 1 nm, and displayed using a color map with a nonlinear (gamma correction with *A* = 1 and $\gamma$ = 0.5) brightness progression. Relevant data properties for the three images (indiscriminate, green channel, red channel) are presented in **Figure S10**.

### SYNTHESIS AND CHARACTERIZATION

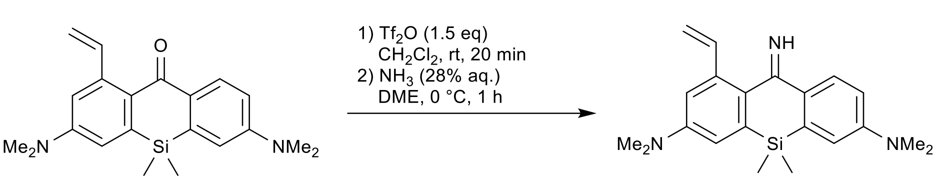

**Compound S2.** A solution of trifluoromethanesulfonic anhydride ($Tf_2O$, 1 M in $CH_2Cl_2$; 0.68 mL, ~0.68 mmol, 1.5 equiv.) was added to the stirred solution of **S1** (160 mg, 0.46 mmol) in dry $CH_2Cl_2$ (7 mL) under argon, and the resulting dark blue solution was stirred at rt for 20 min. It was then transferred dropwise into the stirred mixture of aqueous ammonia (28% aq., 3.5 mL) and 1,2-dimethoxyethane (DME, 6 mL), cooled in ice-water bath. The reaction mixture was stirred at 0 °C for 1 h, diluted with brine (30 mL), the product was then extracted with $CH_2Cl_2$ (3×20 mL) and the combined extracts were dried over $Na_2SO_4$. The product was isolated by flash column chromatography (25 g Interchim SiHP 30 µm cartridge, gradient 0% to 100% A/B, A = $CH_2Cl_2$ – ethanol – 25% aq. $NH_3$ 80:20:2, B = $CH_2Cl_2$) and freeze-dried from 1,4-dioxane to yield 117 mg (73%) of **S2** as brown-orange solid.

$^1$H NMR (400 MHz, $CDCl_3$): δ 7.94 (dd, *J* = 8.2, 1.0 Hz, 1H), 7.05 (dd, *J* = 17.3, 10.8 Hz, 1H), 6.87 – 6.79 (m, 4H), 5.70 (dd, *J* = 17.3, 1.5 Hz, 1H), 5.32 (dd, *J* = 10.8, 1.5 Hz, 1H), 3.05 (s, 6H), 3.02 (s, 6H), 0.45 (s, 6H).

$^{13}$C NMR (101 MHz, $CDCl_3$): δ 173.1, 150.5, 149.9, 138.5, 138.2, 137.8, 136.5, 134.3, 131.7, 128.1, 115.5, 115.4, 114.7, 113.9, 112.9, 40.5, 40.4, -1.8.

HRMS ($C_{21}H_{27}N_3Si$): *m/z* (positive mode) = 350.2042 (found $[M+H]^+$), 350.2047 (calc.).

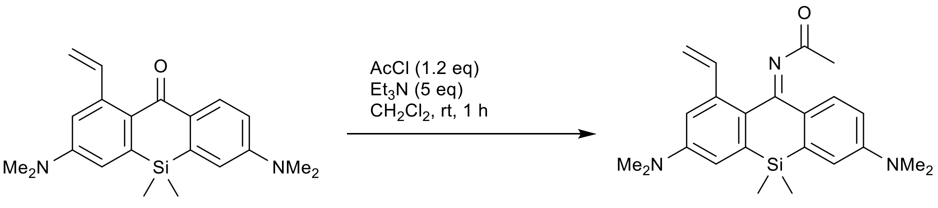

**Compound S3.** Acetyl chloride (4 µL dissolved in 0.5 mL anhydrous $CH_2Cl_2$) was added to the stirred solution of **S1** (16 mg, 45.7 µmol) in dry $CH_2Cl_2$ (1 mL) under argon, and the resulting green solution was stirred at rt for 1 h. The organic solvents were evaporated, and the product was isolated by preparative HPLC (ThermoFisher Hypersil Gold C18 250×21.2 mm 5 µm, solvent flow rate 18 mL/min, gradient 20% to 70% A:B, A – acetonitrile + 0.1% (v/v) $HCO_2H$, B – water + 0.1% (v/v) $HCO_2H$) and freeze-dried from dioxane to give 14 mg (78%) of **S3** as green solid.

$^1$H NMR (400 MHz, $CD_3CN$): δ 7.48 (d, *J* = 8.8 Hz, 1H), 7.33 (dd, *J* = 17.4, 10.9 Hz, 1H), 6.97 – 6.91 (m, 3H), 6.74 (dd, *J* = 8.8, 2.8 Hz, 1H), 5.65 (dd, *J* = 17.4, 1.5 Hz, 1H), 5.20 (dd, *J* = 10.9, 1.5 Hz, 1H), 3.03 (s, 6H), 3.00 (s, 6H), 1.90 (s, 3H), 0.47 (s, 6H).

$^{13}$C NMR (101 MHz, $CD_3CN$): δ 184.1, 162.2, 151.6, 151.3, 139.03, 138.95, 138.85, 138.6, 132.0, 130.7, 129.7, 116.6, 116.1, 114.3, 113.5, 112.2, 40.4, 40.3, 25.6, -2.2.

HRMS ($C_{23}H_{29}N_3OSi$): *m/z* (positive mode) = 392.2146 (found $[M+H]^+$), 392.2153 (calc.).

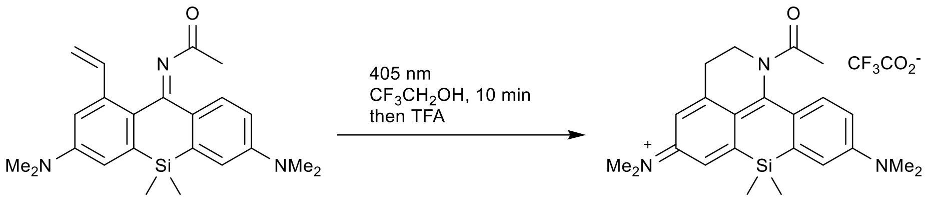

**S3** **1-CF***

**Compound 1-CF*.** Compound **S3** (9 mg, 23.0 µmol) was dissolved in degassed 2,2,2-trifluoroethanol (4.5 mL) in a 10 mL pear-shaped flask equipped with a stirring bar. The solution was irradiated in a Penn OC Photoreactor m1 using a custom-built[1] 405 nm LED light source (50% power, 10 min). Trifluoroacetic acid (10 µL) was then added, the solution was evaporated, and the major product was isolated by preparative HPLC (Interchim Uptisphere Strategy PhC4 250×21.2 mm 5 µm, solvent flow rate 18 mL/min, gradient 30% to 70% A:B, A – acetonitrile + 0.1% (v/v) $CF_3CO_2H$, B – water + 0.1% (v/v) $CF_3CO_2H$) and freeze-dried from dioxane to give 6.6 mg (57%) of **1-CF*** as black solid.

$^1$H NMR (400 MHz, $CDCl_3$): δ 7.65 (d, *J* = 9.4 Hz, 1H), 7.13 (d, *J* = 2.5 Hz, 1H), 7.01 (d, *J* = 2.4 Hz, 1H), 6.80 (dd, *J* = 9.4, 2.5 Hz, 1H), 6.68 – 6.61 (m, 1H), 4.31 (br.s, 2H), 3.37 (s, 6H), 3.36 (s, 6H), 3.09 (br.t, *J* = 7.2 Hz, 2H), 1.97 (s, 3H), 0.55 (s, 6H).

$^{19}$F NMR (376 MHz, $CDCl_3$): δ -75.73 ($CF_3CO_2^-$).

$^{13}$C NMR (101 MHz, $CDCl_3$): δ 137.7, 120.5, 119.3, 114.2, 114.0, 41.0, 30.8, 25.0, -1.3 (indirect detection from a gHSQC experiment, only H-coupled carbons are resolved).

HRMS ($C_{23}H_{30}N_3OSi$): *m/z* (positive mode) = 392.2152 (found $[M]^+$), 392.2153 (calc.).

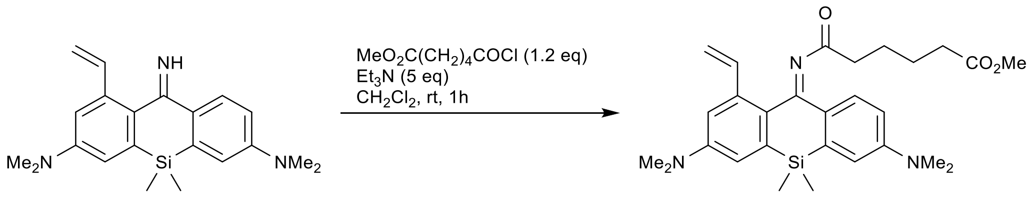

**S2** **S4**

**Compound S4.** Methyl adipoyl chloride solution (36 µL in 0.5 mL of dry $CH_2Cl_2$, 0.23 mmol, 1.2 equiv) was added to the stirred mixture of **S2** (66 mg, 0.19 mmol) and triethylamine (132 µL, 0.95 mmol, 5 equiv) in dry $CH_2Cl_2$ (2 mL), and the resulting green solution was stirred at rt for 1 h. The mixture was diluted with $CH_2Cl_2$ (20 mL), poured into brine (50 mL), and the product was extracted with $CH_2Cl_2$ (2×20 mL). The combined extracts were dried over $Na_2SO_4$, the product was isolated by flash column chromatography (12 g Interchim SiHP 30 µm cartridge, gradient 20% to 70% EtOAc/hexane) and freeze-dried from 1,4-dioxane to yield 77 mg (83%) of **S4** as yellow-green solid.

$^1$H NMR (400 MHz, $CDCl_3$): δ 7.65 (d, *J* = 8.8 Hz, 1H), 7.45 (dd, *J* = 17.3, 10.8 Hz, 1H), 6.90 (d, *J* = 2.8 Hz, 1H), 6.833 (d, *J* = 2.8 Hz, 1H), 6.829 (d, *J* = 2.8 Hz, 1H), 6.66 (dd, *J* = 8.8, 2.8 Hz, 1H), 5.62 (dd, *J* = 17.2, 1.5 Hz, 1H), 5.26 (dd, *J* = 10.8, 1.5 Hz, 1H), 3.61 (s, 3H), 3.04 (s, 6H), 3.02 (s, 6H), 2.20 – 2.12 (m, 4H), 1.56 – 1.46 (m, 4H), 0.49 (s, 6H).

$^{13}$C NMR (101 MHz, $CDCl_3$): δ 185.8, 174.0, 160.7, 150.3, 150.1, 138.5, 138.0, 137.9, 137.6, 131.8, 130.1, 129.7, 115.3, 114.8, 114.1, 112.8, 112.1, 51.5, 40.3, 40.2, 37.5, 33.9, 24.6, 24.2, -2.0.

HRMS ($C_{28}H_{37}N_3O_3Si$): *m/z* (positive mode) = 492.2672 (found $[M+H]^+$), 492.2677 (calc.).

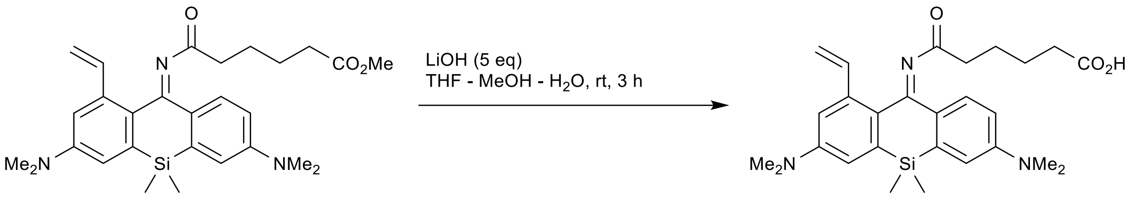

**S3** **1**

**Compound 1.** Lithium hydroxide solution (33 mg of LiOH·$H_2O$ in 1 mL water, 0.78 mmol, 5 equiv) was added to the stirred solution of **S4** (77 mg, 0.156 mmol) in THF (4 mL) and methanol (1 mL), and the reaction mixture was stirred vigorously for 3 h. The organic solvents were evaporated, and the product was isolated by preparative HPLC (Interchim Uptisphere Strategy PhC4 250×21.2 mm 5 µm, solvent flow rate 18 mL/min, gradient 20% to 70% A:B, A – acetonitrile + 0.1% (v/v) $HCO_2H$, B – water + 0.1% (v/v) $HCO_2H$) and freeze-dried from dioxane to give 65 mg (87%) of **1** as green-yellow solid.

$^1$H NMR (400 MHz, $CDCl_3$): δ 7.65 (d, *J* = 8.8 Hz, 1H), 7.44 (dd, *J* = 17.3, 10.8 Hz, 1H), 6.90 (d, *J* = 2.8 Hz, 1H), 6.84 – 6.81 (m, 2H), 6.66 (dd, *J* = 8.9, 2.8 Hz, 1H), 5.62 (dd, *J* = 17.3, 1.5 Hz, 1H), 5.26 (dd, *J* = 10.8, 1.5 Hz, 1H), 3.04 (s, 6H), 3.01 (s, 6H), 2.17 (app.q, *J* = 7.2 Hz, 4H), 1.59 – 1.44 (m, 4H), 0.48 (s, 6H).

$^{13}$C NMR (101 MHz, $CDCl_3$): δ 185.8, 178.7, 160.9, 150.4, 150.2, 138.6, 138.0, 137.6, 131.8, 130.2, 129.8, 115.4, 114.9, 114.2, 112.9, 112.1, 40.4, 40.2, 37.4, 33.7, 24.3, 24.0, -2.0.

HRMS ($C_{27}H_{35}N_3O_3Si$): *m/z* (positive mode) = 478.2516 (found $[M+H]^+$), 478.2520 (calc.).

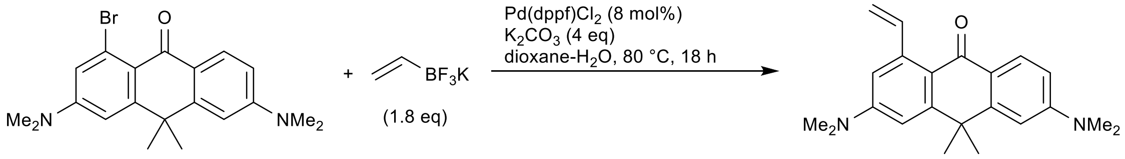

**S5** **S6**

**Compound S6.** A mixture of **S5** (220 mg, 0.568 mmol),[1] potassium vinyltrifluoroborate (229 mg, 1.70 mmol, 3 equiv), [1,1′-bis(diphenylphosphino)ferrocene]dichloropalladium(II) (33 mg, 45.4 µmol, 8 mol%) and potassium carbonate (312 mg, 2.27 mmol, 4 equiv) in 1,4-dioxane (5 mL) and water (1 mL) was stirred at 80 °C for 18 h. Upon cooling, the reaction mixture was diluted with brine (50 mL) and extracted with $CH_2Cl_2$ (3×20 mL). The combined extracts were dried over $Na_2SO_4$, and the product was isolated by flash column chromatography (12 g Interchim SiHP 30 µm cartridge, gradient 5% to 30% EtOAc/hexane + 20% $CH_2Cl_2$ constant additive) and freeze-dried from 1,4-dioxane to yield 168 mg (89%) of **S6** as yellow solid.

$^1$H NMR (400 MHz, $CDCl_3$): δ 8.20 (d, *J* = 8.4, 1H), 7.90 (dd, *J* = 17.2, 10.8, 1H), 6.79 (d, *J* = 2.7 Hz, 1H), 6.77 – 6.72 (m, 2H), 6.70 (d, *J* = 2.7, 1H), 5.44 (dd, *J* = 17.2, 2.0 Hz, 1H), 5.27 (dd, *J* = 10.8, 2.0 Hz, 1H), 3.11 (s, 6H), 3.09 (s, 6H), 1.72 (s, 6H).

$^{13}$C NMR (101 MHz, $CDCl_3$): δ 182.9, 153.6, 153.0, 152.2, 151.3, 143.6, 141.8, 129.3, 121.5, 118.1, 113.2, 111.5, 111.1, 108.3, 107.3, 40.4, 40.3, 38.9, 34.3.

HRMS ($C_{22}H_{26}N_2O$): *m/z* (positive mode) = 335.2118 (found $[M+H]^+$), 335.2118 (calc.).

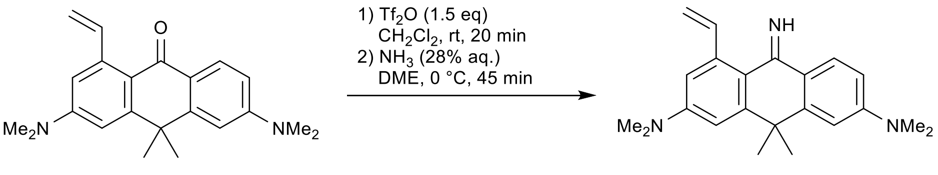

**Compound S7.** A solution of trifluoromethanesulfonic anhydride ($Tf_2O$, 1 M in $CH_2Cl_2$; 0.72 mL, ~0.72 mmol, 1.5 equiv.) was added to the stirred solution of **S6** (160 mg, 0.479 mmol) in dry $CH_2Cl_2$ (7 mL) under argon, and the resulting blue-violet solution was stirred at rt for 20 min. It was then transferred dropwise into the stirred mixture of aqueous ammonia (28% aq., 3.5 mL) and 1,2-dimethoxyethane (DME, 6 mL), cooled in ice-water bath. The reaction mixture was stirred at 0 °C for 45 min, diluted with brine (30 mL), the product was then extracted with $CH_2Cl_2$ (4×20 mL) and the combined extracts were dried over $Na_2SO_4$. The product was isolated by flash column chromatography (12 g Interchim SiHP 30 µm cartridge, gradient 0% to 100% A/B, A = $CH_2Cl_2$ – ethanol – 25% aq. $NH_3$ 80:20:2, B = $CH_2Cl_2$) and freeze-dried from 1,4-dioxane to yield 83 mg (52%) of **S7** as brown-orange solid.

$^1$H NMR (400 MHz, $CDCl_3$): δ 8.9 (br.s, 1 H), 7.96 (br.d, *J* = 7.5 Hz, 1H), 7.26 – 7.14 (m, 1H), 6.86 (d, *J* = 2.6 Hz, 1H), 6.82 (d, *J* = 2.5 Hz, 1H), 6.75 – 6.68 (m, 2H), 5.70 (dd, *J* = 17.3, 1.6 Hz, 1H), 5.35 (dd, *J* = 10.8, 1.6 Hz, 1H), 3.06 (s, 6H), 3.04 (s, 6H), 1.65 (s, 6H).

$^{13}$C NMR (101 MHz, $CDCl_3$): δ 167.7, 151.8, 150.9, 149.1, 147.7, 139.0, 138.8, 126.6, 124.4, 121.2, 115.6, 110.9, 110.8, 107.8, 107.1, 40.6, 40.5, 39.8, 32.0.

HRMS ($C_{22}H_{27}N_3$): *m/z* (positive mode) = 334.2277 (found $[M+H]^+$), 334.2278 (calc.).

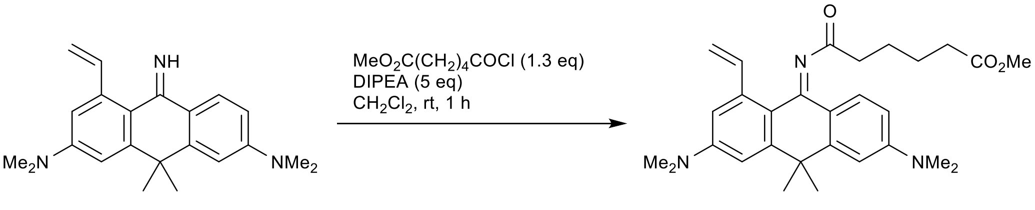

**Compound S8.** Methyl adipoyl chloride solution (24 µL in 0.5 mL of dry $CH_2Cl_2$, 0.156 mmol, 1.3 equiv) was added to the stirred mixture of **S7** (40 mg, 0.12 mmol) and DIPEA (104 µL, 0.6 mmol, 5 equiv) in dry $CH_2Cl_2$ (1.5 mL), and the resulting solution was stirred at rt for 1 h. The mixture was evaporated to dryness, and the product was isolated by flash column chromatography (12 g Interchim SiHP 30 µm cartridge, gradient 20% to 100% EtOAc/hexane) and freeze-dried from 1,4-dioxane to yield 48.5 mg (85%) of **S8** as yellow solid.

$^1$H NMR (400 MHz, $CDCl_3$): δ 7.64 (d, *J* = 8.8 Hz, 1H), 7.60 (dd, *J* = 17.2, 10.8 Hz, 1H), 6.84 (d, *J* = 2.6 Hz, 1H), 6.81 (d, *J* = 2.6 Hz, 1H), 6.76 (d, *J* = 2.6 Hz, 1H), 6.57 (dd, *J* = 8.8, 2.6 Hz, 1H), 5.56 (dd, *J* = 17.2, 1.7 Hz, 1H), 5.22 (dd, *J* = 10.8, 1.7 Hz, 1H), 3.62 (s, 3H), 3.07 (s, 6H), 3.05 (s, 6H), 2.32 (t, *J* = 7.0 Hz, 2H), 2.24 (t, *J* = 7.2 Hz, 2H), 1.67 (s, 6H), 1.65 – 1.52 (m, 4H).

$^{13}$C NMR (101 MHz, $CDCl_3$): δ 185.2, 174.1, 154.6, 151.8, 151.2, 149.9, 149.4, 139.8, 139.1, 129.1, 121.3, 119.8, 113.6, 110.14, 110.12, 107.4, 107.0, 51.5, 40.4, 40.3, 40.2, 37.7, 34.0, 32.1, 24.73, 24.70.

HRMS ($C_{29}H_{37}N_3O_3$): *m/z* (positive mode) = 476.2906 (found $[M+H]^+$), 476.2908 (calc.).

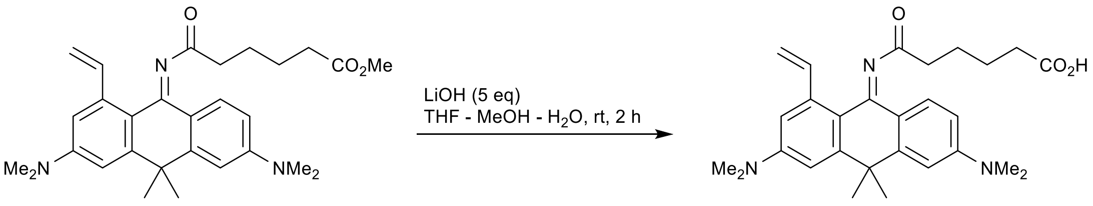

**Compound 2.** Lithium hydroxide solution (21 mg of LiOH·$H_2O$ in 0.5 mL water, 0.50 mmol, 5 equiv) was added to the stirred solution of **S8** (48 mg, 0.1 mmol) in THF (2 mL) and methanol (0.5 mL), and the reaction mixture was stirred at rt for 2 h. Acetic acid (150 µL) was then added, and the reaction mixture was evaporated to dryness. The product was isolated by preparative HPLC (ThermoFisher Hypersil Gold C18 250×21.2 mm 5 µm, solvent flow rate 18 mL/min, gradient 20% to 80% A:B, A – acetonitrile + 0.1% (v/v) $HCO_2H$, B – water + 0.1% (v/v) $HCO_2H$) and freeze-dried from dioxane to give 42 mg (91%) of **2** as blue-green solid.

$^{1}H$ NMR (400 MHz, $CD_3CN$): δ 7.53 (dd, *J* = 17.4, 10.8 Hz, 1H), 7.46 (d, *J* = 8.8 Hz, 1H), 6.96 (d, *J* = 2.6 Hz, 1H), 6.94 (d, *J* = 2.6 Hz, 1H), 6.81 (d, *J* = 2.6 Hz, 1H), 6.66 (dd, *J* = 8.8, 2.6 Hz, 1H), 5.61 (dd, *J* = 17.4, 1.7 Hz, 1H), 5.19 (dd, *J* = 10.8, 1.7 Hz, 1H), 3.09 (s, 6H), 3.05 (s, 6H), 2.36 (t, *J* = 7.1 Hz, 2H), 2.23 (t, *J* = 7.1 Hz, 2H), 1.68 (s, 6H), 1.66 – 1.52 (m, 4H).

$^{13}C$ NMR (101 MHz, $CD_3CN$): δ 185.8, 175.0, 156.2, 153.1, 152.4, 151.1, 150.4, 140.2, 139.9, 129.3, 121.3, 120.2, 113.8, 110.8, 110.5, 108.6, 108.2, 41.0, 40.5, 40.4, 38.4, 33.9, 32.1, 25.3, 25.2.

HRMS ($C_{28}H_{35}N_3O_3$): *m/z* (positive mode) = 462.2749 (found $[M+H]^+$), 462.2751 (calc.).

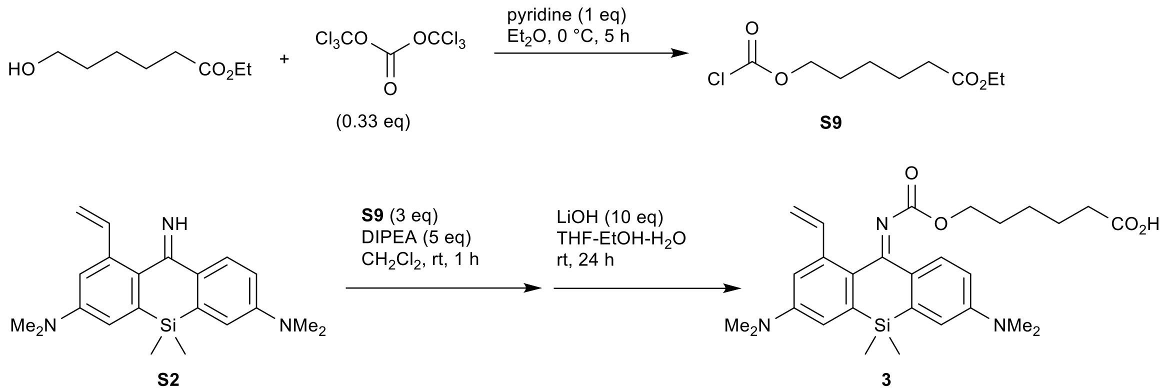

**Compound 3.** Chloroformate ester **S9** was prepared according to the procedure from.[13] Ethyl 6-hydroxyhexanoate (160 mg, 1 mmol) and pyridine (81 µL, 1 mmol, 1 equiv) were added to the stirred solution of triphosgene (99 mg, 0.33 mmol, 0.33 equiv) in dry diethyl ether (2 mL), cooled to 0 °C. The reaction mixture was stirred at 0 °C for 5 h, the precipitate of pyridine hydrochloride was filtered off on a short plug of Celite, washed with dry diethyl ether (3 mL), the filtrate was evaporated and the residue was redissolved in dry $CH_2Cl_2$ (1 mL) and used directly in the next step.

The prepared solution of **S9** in $CH_2Cl_2$ (0.3 mL, ~0.3 mmol, 3 equiv) was added to the stirred solution of **S2** (35 mg, 0.1 mmol) and DIPEA (87 µL, 0.5 mmol, 5 equiv) in dry $CH_2Cl_2$ (0.5 mL), and the resulting solution was stirred at rt for 1 h. The crude reaction mixture was evaporated, the residue was dissolved in the mixture of THF (2 mL) and ethanol (0.5 mL), and lithium hydroxide solution (21 mg of $LiOH{\cdot}H_2O$ in 0.5 mL water, 0.5 mmol, 5 equiv) was added to the mixture, which was left stirring at rt for 24 h (the second portion of $LiOH{\cdot}H_2O$ (21 mg, 0.5 mmol, 5 equiv) was added after 8 h). Acetic acid (100 µL) was then added, and the reaction mixture was evaporated to dryness. The product was isolated by preparative HPLC (ThermoFisher Hypersil Gold C18 250×21.2 mm 5 µm, solvent flow rate 18 mL/min, gradient 40% to 100% A:B, A – acetonitrile + 0.1% (v/v) $HCO_2H$, B – water + 0.1% (v/v) $HCO_2H$) and freeze-dried from dioxane to give 47 mg (93% over 2 steps) of **3** as yellow solid.

$^{1}$H NMR (400 MHz, $CDCl_3$): δ 7.69 (d, *J* = 8.7 Hz, 1H), 7.37 (dd, *J* = 17.3, 10.8 Hz, 1H), 6.92 (br.s, 1H), 6.88 (br.s, 2H), 6.72 (br.s, 1H), 5.66 (dd, *J* = 17.3, 1.5 Hz, 1H), 5.29 (dd, *J* = 10.8, 1.5 Hz, 1H), 4.02 (t, *J* = 6.2 Hz, 2H), 3.05 (s, 6H), 3.02 (s, 6H), 2.22 (t, *J* = 7.5 Hz, 2H), 1.57 – 1.44 (m, 4H), 1.16 – 1.03 (m, 2H), 0.49 (s, 6H).

$^{13}$C NMR (101 MHz, $CDCl_3$): δ 178.5, 168.8, 162.8, 150.0, 138.2, 137.9, 137.3, 128.1, 115.2, 112.8, 111.6, 65.4, 40.4, 33.7, 28.3, 25.2, 24.2, -2.5.

HRMS ($C_{28}H_{37}N_3O_4Si$): *m/z* (positive mode) = 508.2640 (found $[M+H]^+$), 508.2626 (calc.).

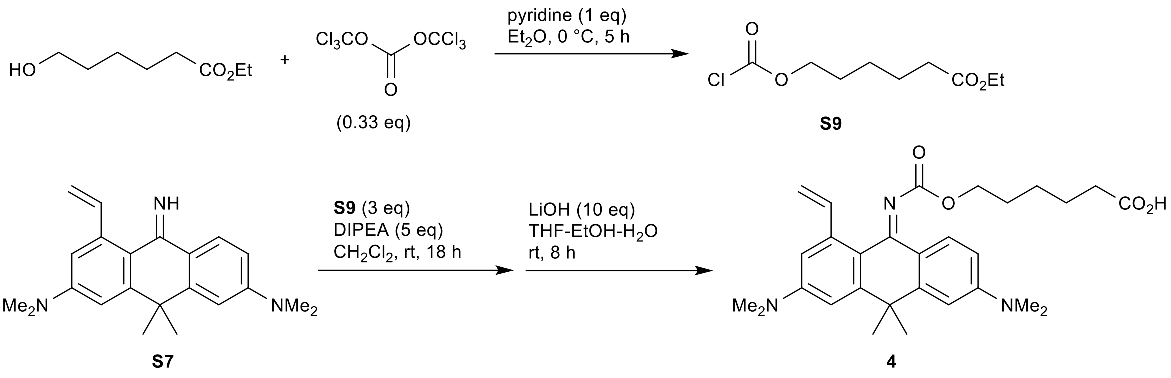

**Compound 4.** Chloroformate ester **S9** was prepared according to the procedure from [13]. Ethyl 6-hydroxyhexanoate (160 mg, 1 mmol) and pyridine (81 µL, 1 mmol, 1 equiv) were added to the stirred solution of triphosgene (99 mg, 0.33 mmol, 0.33 equiv) in dry diethyl ether (2 mL), cooled to 0 °C. The reaction mixture was stirred at 0 °C for 5 h, the precipitate of pyridine hydrochloride was filtered off on a short plug of Celite, washed with dry diethyl ether (3 mL), the filtrate was evaporated and the residue was redissolved in dry $CH_2Cl_2$ (1 mL) and used directly in the next step.

The prepared solution of **S9** in $CH_2Cl_2$ (0.36 mL, ~0.36 mmol, 3 equiv) was added to the stirred solution of **S7** (40 mg, 0.12 mmol) and DIPEA (104 µL, 0.6 mmol, 5 equiv) in dry $CH_2Cl_2$ (1.2 mL), and the resulting solution was stirred at rt for 18 h. The crude reaction mixture was evaporated, the residue was dissolved in the mixture of THF (2 mL) and ethanol (0.5 mL), and lithium hydroxide solution (50 mg of $LiOH{\cdot}H_2O$ in 0.5 mL water, 1.2 mmol, 10 equiv) was added to the mixture, which was left stirring at rt for 8 h. Acetic acid (150 µL) was then added, and the reaction mixture was evaporated to dryness. The product was isolated by preparative HPLC (ThermoFisher Hypersil Gold C18 250×21.2 mm 5 µm, solvent flow rate 18 mL/min, gradient 20% to 80% A:B, A – acetonitrile + 0.1% (v/v) $HCO_2H$, B – water + 0.1% (v/v) $HCO_2H$) and freeze-dried from dioxane to give 66 mg (95% over 2 steps) of **4** as green hygroscopic solid, containing ~ 1 eq./eq. 1,4-dioxane.

$^1$H NMR (400 MHz, $CDCl_3$): δ 7.67 (d, $J$ = 8.8 Hz, 1H), 7.55 (dd, $J$ = 17.3, 10.8 Hz, 1H), 6.83 (d, $J$ = 2.5 Hz, 1H), 6.81 (d, $J$ = 2.5 Hz, 1H), 6.75 (d, $J$ = 2.5 Hz, 1H), 6.57 (dd, $J$ = 8.8, 2.5 Hz, 1H), 5.63 (dd, $J$ = 17.3, 1.6 Hz, 1H), 5.25 (dd, $J$ = 10.8, 1.6 Hz, 1H), 4.10 (t, $J$ = 6.6 Hz, 2H), 3.06 (s, 6H), 3.03 (s, 6H), 2.22 (t, $J$ = 7.5 Hz, 2H), 1.66 (s, 6H), 1.62 – 1.36 (m, 4H), 1.27 – 1.09 (m, 2H).

$^{13}$C NMR (101 MHz, $CDCl_3$): δ 179.0, 163.7, 163.5, 151.7, 151.2, 150.0, 149.3, 139.1, 138.4, 127.7, 122.7, 120.8, 114.1, 109.9, 109.4, 107.1, 106.7, 65.5, 40.43, 40.37, 34.0, 31.5, 28.5, 25.5, 24.4.

HRMS ($C_{29}H_{37}N_3O_4$): $m/z$ (positive mode) = 492.2859 (found $[M+H]^+$), 492.2857 (calc.).

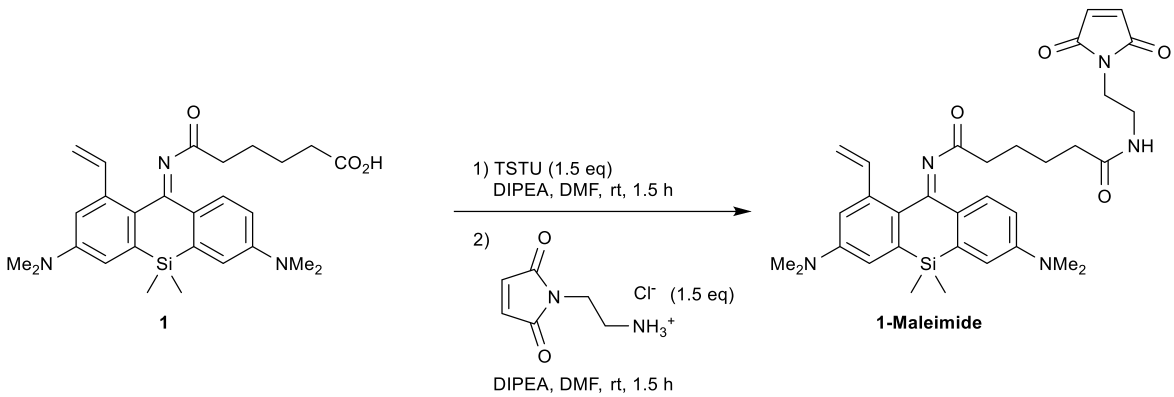

**Compound 1-Maleimide.** TSTU solution (*N,N,N',N'*-tetramethyl-*O*-(*N*-succinimidyl)uronium tetrafluoroborate; 9.4 mg in 50 µL DMF, 31.4 µmol, 1.5 equiv) was added to the stirred solution of **1** (10 mg, 20.9 µmol) in DMF (100 µL) and DIPEA (30 µL), and the reaction mixture was stirred at rt for 1.5 h. A solution of 1-(2-aminoethyl)maleimide hydrochloride (5.6 mg, 31.4 µmol, 1.5 equiv) was then added, followed by DIPEA (50 µL) and the reaction mixture was stirred for further 1.5 h. The organic solvents were evaporated *in vacuo*, and the product was isolated by preparative HPLC (Interchim Uptisphere Strategy PhC4 250×21.2 mm 5 µm, solvent flow rate 18 mL/min, gradient 20% to 70% A:B, A – acetonitrile + 0.1% (v/v) $HCO_2H$, B – water + 0.1% (v/v) $HCO_2H$) to give 12.5 mg (99%) of **1-Maleimide** as brown yellow-solid.

$^1$H NMR (400 MHz, $CDCl_3$): δ 7.62 (d, *J* = 8.8 Hz, 1H), 7.43 (dd, *J* = 17.3, 10.9 Hz, 1H), 6.90 (d, *J* = 2.7 Hz, 1H), 6.83 (dd, *J* = 2.7, 1.4 Hz, 2H), 6.72 – 6.65 (m, 3H), 6.18 (br.t, *J* = 5.7 Hz, 1H), 5.62 (dd, *J* = 17.3, 1.5 Hz, 1H), 5.26 (dd, *J* = 10.9, 1.5 Hz, 1H), 3.68 – 3.62 (m, 2H), 3.48 – 3.39 (m, 2H), 3.05 (s, 6H), 3.03 (s, 6H), 2.17 – 2.10 (m, 3H), 2.08 – 2.01 (m, 2H), 1.54 – 1.43 (m, 4H), 0.48 (s, 6H).

$^{13}$C NMR (101 MHz, $CDCl_3$): δ 186.2, 173.4, 171.1, 160.9, 150.4, 150.2, 138.5, 138.0, 137.7, 134.3, 131.7, 130.1, 129.7, 115.4, 114.9, 114.1, 112.8, 112.1, 40.4, 40.2, 38.7, 37.8, 37.3, 36.2, 24.9, 24.0, -2.0.

HRMS ($C_{33}H_{41}N_5O_4Si$): *m/z* (positive mode) = 600.2996 (found $[M+H]^+$), 600.3001 (calc.).

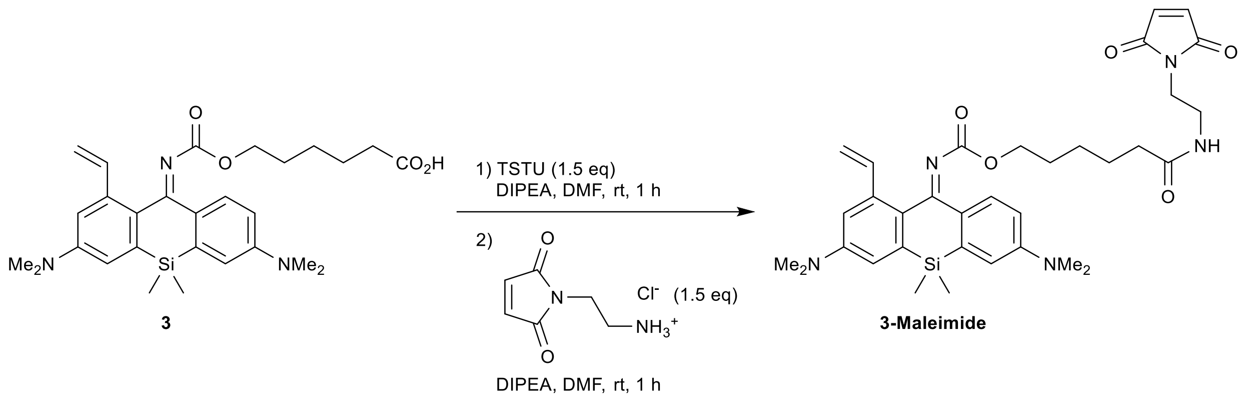

**Compound 3-Maleimide.** TSTU solution (*N,N,N',N'*-tetramethyl-*O*-(*N*-succinimidyl)uronium tetrafluoroborate; 9.0 mg in 50 µL DMF, 30 µmol, 1.5 equiv) was added to the stirred solution of **3** (10 mg, 19.7 µmol) in DMF (100 µL) and DIPEA (30 µL), and the reaction mixture was stirred at rt for 1 h. A solution of 1-(2-aminoethyl)maleimide hydrochloride (5.3 mg, 30 µmol, 1.5 equiv) was then added, followed by DIPEA (50 µL) and the reaction mixture was stirred for further 1 h. The organic solvents were evaporated *in vacuo*, and the product was isolated by preparative HPLC (ThermoFisher Hypersil Gold C18 250×21.2 mm 5 µm, solvent flow rate 18 mL/min, gradient 30% to 100% A:B, A – acetonitrile + 0.1% (v/v) $HCO_2H$, B – water + 0.1% (v/v) $HCO_2H$) to give 8.5 mg (68%) of **3-Maleimide** as yellow solid.

HRMS ($C_{33}H_{43}N_5O_5Si$): *m/z* (positive mode) = 630.3104 (found $[M+H]^+$), 630.3106 (calc.).

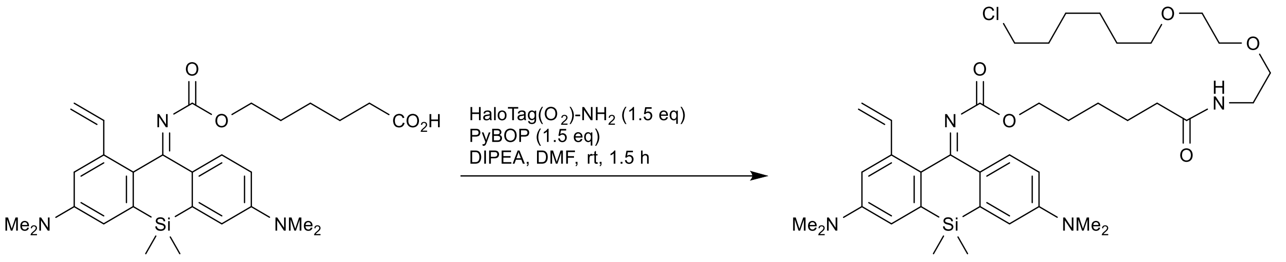

**3** **3-Halo**

**Compound 3-Halo.** PyBOP solution (benzotriazol-1-yloxy-tris(pyrrolidino)phosphonium hexafluorophosphate; 15.6 mg in 100 µL DMF, 30.0 µmol, 1.5 equiv) was added to the stirred solution of **3** (10 mg, 19.7 µmol) and HaloTag(O2) amine (7.7 mg, 30.0 µmol, 1.5 equiv) in DMF (200 µL) and DIPEA (70 µL), and the reaction mixture was stirred at rt for 1.5 h. The organic solvents were evaporated *in vacuo*, and the product was isolated by preparative HPLC (ThermoFisher Hypersil Gold C18 250×21.2 mm 5 µm, solvent flow rate 18 mL/min, gradient 50% to 100% A:B, A – acetonitrile + 0.1% (v/v) $HCO_2H$, B – water + 0.1% (v/v) $HCO_2H$) to give 11 mg (78%) of **3-Halo** as viscous yellow oil.

$^1$H NMR (400 MHz, $CDCl_3$): δ 7.66 (d, *J* = 8.8 Hz, 1H), 7.36 (dd, *J* = 17.3, 10.8 Hz, 1H), 6.89 (d, *J* = 2.7 Hz, 1H), 6.84 (d, *J* = 2.7 Hz, 1H), 6.83 (d, *J* = 2.7 Hz, 1H), 6.68 (dd, *J* = 8.8, 2.7 Hz, 1H), 5.97 (t, *J* = 5.6 Hz, 1H), 5.65 (dd, *J* = 17.3, 1.5 Hz, 1H), 5.26 (dd, *J* = 10.8, 1.5 Hz, 1H), 4.02 (t, *J* = 6.4 Hz, 2H), 3.63 – 3.50 (m, 8H), 3.48 – 3.41 (m, 4H), 3.04 (s, 6H), 3.01 (s, 6H), 2.11 – 2.04 (m, 2H), 1.77 (dq, *J* = 8.0, 6.7 Hz, 2H), 1.60 (dq, *J* = 8.0, 6.8 Hz, 2H), 1.56 – 1.32 (m, 8H), 1.21 – 1.10 (m, 2H), 0.48 (s, 6H).

$^{13}$C NMR (101 MHz, $CDCl_3$): δ 173.0, 169.1, 163.0, 150.4, 150.2, 138.2, 137.9, 137.8, 137.6, 132.5, 130.7, 128.1, 115.4, 115.0, 114.3, 112.8, 111.5, 71.4, 70.4, 70.2, 70.1, 65.6, 45.2, 40.4, 40.3, 39.3, 36.7, 32.6, 29.6, 28.6, 26.8, 25.6, 25.54, 25.45, -2.3.

HRMS ($C_{38}H_{57}ClN_4O_5Si$): *m/z* (positive mode) = 713.3865 (found $[M+H]^+$), 713.3860 (calc.).

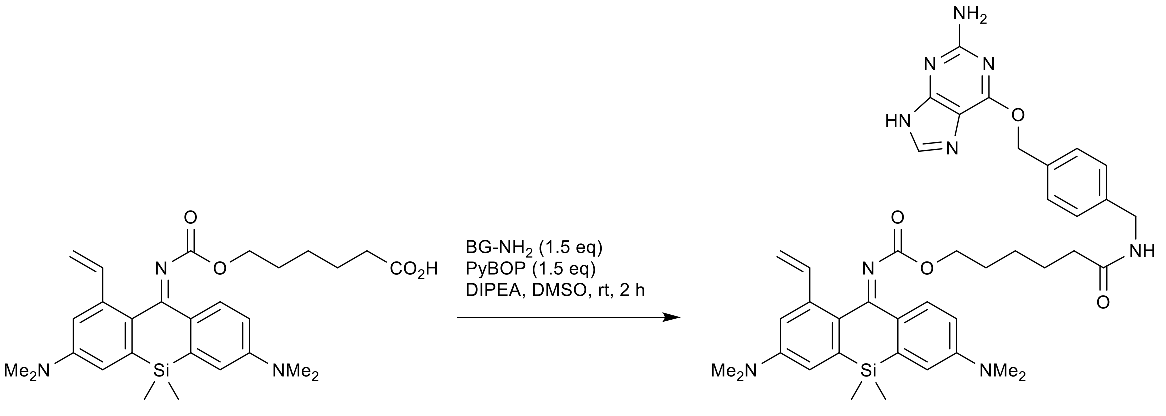

**3** **3-BG**

**Compound 3-BG.** PyBOP solution (benzotriazol-1-yloxy-tris(pyrrolidino)phosphonium hexafluorophosphate; 15.3 mg in 50 µL DMSO, 29.5 µmol, 1.5 equiv) was added to the stirred solution of **3** (10 mg, 19.7 µmol) and BG-$NH_2$ (8 mg, 29.5 µmol, 1.5 equiv) in DMSO (100 µL) and DIPEA (50 µL), and the reaction mixture was stirred at rt for 2 h. The volatiles were evaporated *in vacuo*, and the product was isolated by preparative HPLC (ThermoFisher Hypersil Gold C18 250×21.2 mm 5 µm, solvent flow rate 18 mL/min, gradient 30% to 90% A:B, A – acetonitrile + 0.1% (v/v) $HCO_2H$, B – water + 0.1% (v/v) $HCO_2H$) to give 14 mg (94%) of **3-BG** as greenish-yellow solid.

HRMS ($C_{41}H_{49}N_9O_4Si$): *m/z* (positive mode) = 380.6905 (found $[M+2H]^{2+}$), 380.6911 (calc.).

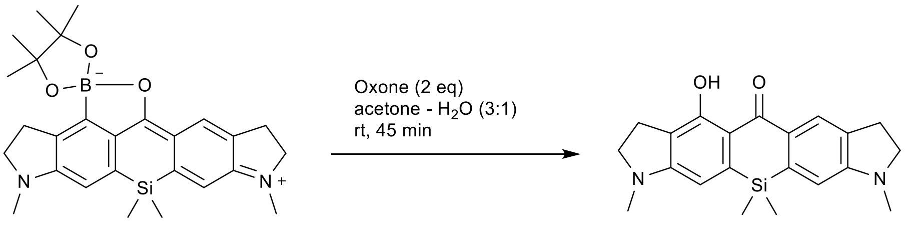

**S10** **S11**

**Compound S11.** Oxone (potassium peroxymonosulfate triple salt $KHSO_5 \cdot 0.5KHSO_4 \cdot 0.5K2SO_4$; 461 mg, 1.5 mmol, 2 equiv) was added to the stirred suspension of **S10**[7] (356 mg, 0.75 mmol) in acetone (10 mL) and water (3.3 mL) at rt. The orange color of the reaction mixture quickly changed to brown. After 45 min, the reaction mixture was diluted with water (100 mL), the product was extracted with ethyl acetate (3×50 mL) and $CH_2Cl_2$ (3×25 mL), the combined extracts were washed with brine (50 mL) and dried over $Na_2SO_4$. The solution was filtered through a plug of silica (3 cm), washing with EtOAc – $CH_2Cl_2$ (1:1, 150 mL). The filtrate was evaporated to orange crystals, yield 234 mg (85%). Known compound, the analytical data match those in [7].

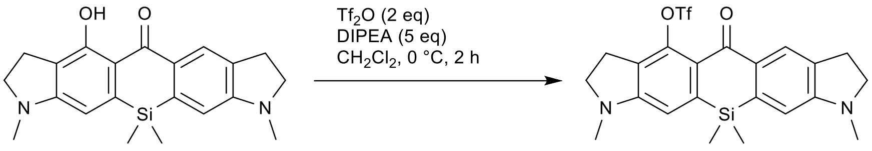

**S11** **S12**

**Compound S12.** Trifluoromethanesulfonic anhydride (0.7 mL of 1 M in $CH_2Cl_2$, ~0.7 mmol, 1 equiv) was added to the stirred solution of **S11** (242 mg, 0.66 mmol) and DIPEA (0.57 mL, 3.28 mmol, 5 equiv) in dry $CH_2Cl_2$, cooled in ice-water bath. The reaction mixture was stirred at 0 °C for 1 h, and another portion of trifluoromethanesulfonic anhydride (0.7 mL of 1 M in $CH_2Cl_2$, ~0.7 mmol, 1 equiv) was added. After stirring for further 1 h at 0 °C, the reaction mixture was quenched by addition of sat. aq. $NH_4Cl$ (50 mL), the product was extracted with $CH_2Cl_2$ (3×40 mL) and the combined extracts were dried over $Na_2SO_4$. The filtrate was evaporated on silica and the product was isolated by flash column chromatography (25 g Interchim SiHP 30 µm cartridge, gradient 5% to 50% EtOAc/hexane) and freeze-dried from 1,4-dioxane to yield 181 mg (55%) of **S12** as yellow solid. Known compound, the analytical data match those in [7].

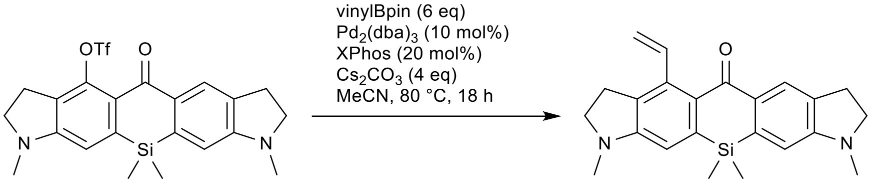

**S12** **S13**

**Compound S13.** In a 25 mL round-bottom flask, a mixture of **S12** (181 mg, 0.36 mmol), vinylboronic acid pinacol ester (336 mg, 2.18 mmol, 6 equiv), tris(dibenzylideneacetone)dipalladium(0) (33 mg, 36 µmol, 10 mol%), XPhos (35 mg, 72 µmol, 20 mol%) and cesium carbonate (474 mg, 1.44 mmol, 4 equiv) in degassed dry acetonitrile (6 mL) was stirred at 80 °C overnight (18 h). On cooling, the reaction mixture was diluted with $CH_2Cl_2$ and filtered through a plug of Celite, washing with $CH_2Cl_2$ (100 mL). The filtrate was washed with brine, dried over $Na_2SO_4$ and the product was isolated by flash column chromatography (25 g Interchim SiHP 30 µm cartridge, gradient 5% to 60% EtOAc/hexane) and freeze-dried from 1,4-dioxane to yield 92 mg (67%) of **S13** as yellow solid.

$^1$H NMR (400 MHz, $CDCl_3$): δ 8.04 (t, *J* = 1.2 Hz, 2H), 7.21 (dd, *J* = 17.8, 11.3 Hz, 1H), 6.48 (d, *J* = 2.1 Hz, 2H), 5.38 (dd, *J* = 11.3, 1.9 Hz, 1H), 5.15 (dd, *J* = 17.8, 1.9 Hz, 1H), 3.46 – 3.38 (m, 4H), 3.14 – 3.06 (m, 2H), 3.01 (td, *J* = 8.4, 1.2 Hz, 2H), 2.878 (s, 3H), 2.876 (s, 3H), 0.44 (s, 6H).

$^{13}$C NMR (101 MHz, $CDCl_3$): δ 188.0, 154.6, 154.5, 141.1, 138.9, 138.8, 138.4, 134.1, 132.3, 131.3, 131.0, 126.0, 113.9, 107.8, 107.6, 55.3, 55.1, 35.0, 34.9, 29.2, 28.3, -1.0.

HRMS ($C_{23}H_{26}N_2OSi$): *m/z* (positive mode) = 375.1882 (found $[M+H]^+$), 375.1887 (calc.).

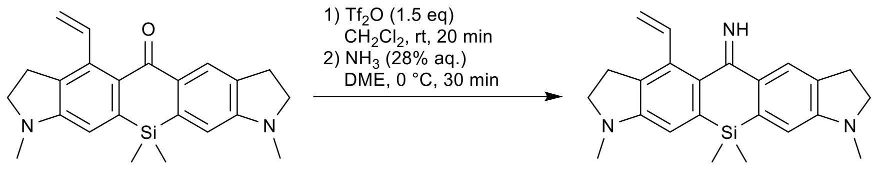

**S13** **S14**

**Compound S14.** A solution of trifluoromethanesulfonic anhydride ($Tf_2O$, 1 M in $CH_2Cl_2$; 0.37 mL, ~0.37 mmol, 1.5 equiv.) was added under argon atmosphere to the stirred solution of **S13** (92 mg, 0.25 mmol) in dry $CH_2Cl_2$ (3 mL), cooled in ice-water bath, and the resulting dark blue solution was stirred at 0 °C for 20 min. It was then transferred dropwise into the stirred mixture of aqueous ammonia (28% aq., 2.5 mL) and 1,2-dimethoxyethane (DME, 5 mL), cooled in ice-water bath. The reaction mixture was stirred at 0 °C for 30 min, diluted with brine 230 mL), the product was then extracted with $CH_2Cl_2$ (3×20 mL) and the combined extracts were dried over $Na_2SO_4$. The product was isolated by flash column chromatography (12 g Interchim SiHP 30 µm cartridge, gradient 0% to 100% A/B, A = $CH_2Cl_2$ – ethanol – 25% aq. $NH_3$ 80:20:2, B = $CH_2Cl_2$) and freeze-dried from 1,4-dioxane to yield 19 mg (21%) of **S14** as brown-orange solid.

$^1$H NMR (400 MHz, $CDCl_3$): δ 7.78 (s, 1H), 6.80 (dd, *J* = 17.9, 11.5 Hz, 1H), 6.54 (d, *J* = 6.5 Hz, 2H), 5.52 (dd, *J* = 11.5, 1.6 Hz, 1H), 5.45 (dd, *J* = 17.9, 1.6 Hz, 1H), 3.40 – 3.32 (m, 4H), 3.07 (t, *J* = 8.2 Hz, 2H), 3.00 (t, *J* = 8.5 Hz, 1H), 2.83 (s, 6H), 0.43 (s, 6H).

$^{13}$C NMR (101 MHz, $CDCl_3$): δ 173.8, 153.8, 153.4, 137.1, 136.0, 135.6, 133.4, 132.5, 131.2, 123.6, 118.9, 108.73, 108.67, 55.8, 55.7, 35.7, 29.8, 29.3, 28.6, -1.8.

HRMS ($C_{23}H_{27}N_3Si$): *m/z* (positive mode) = 374.2062 (found $[M+H]^+$), 374.2047 (calc.).

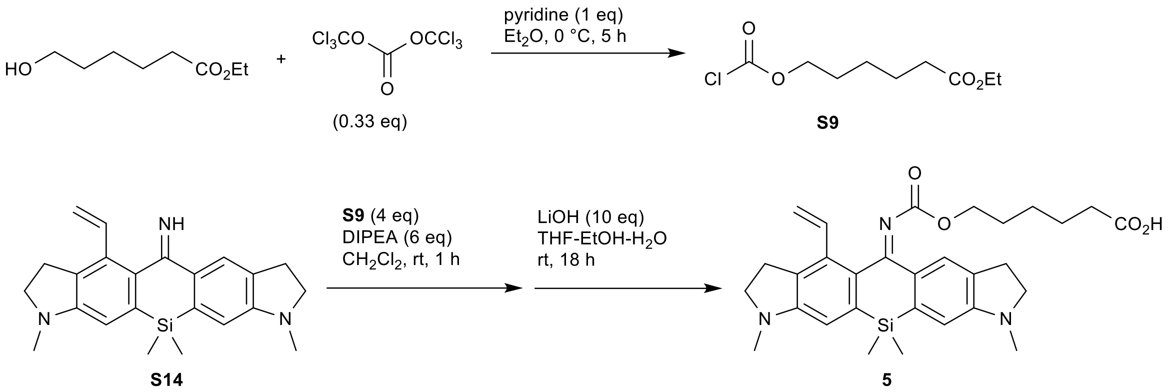

**Compound 5.** Chloroformate ester **9** was prepared according to the procedure from [13]. Ethyl 6-hydroxyhexanoate (160 mg, 1 mmol) and pyridine (81 µL, 1 mmol, 1 equiv) were added to the stirred solution of triphosgene (99 mg, 0.33 mmol, 0.33 equiv) in dry diethyl ether (2 mL), cooled to 0 °C. The reaction mixture was stirred at 0 °C for 5 h, the precipitate of pyridine hydrochloride was filtered off on a short plug of Celite, washed with dry diethyl ether (3 mL), the filtrate was evaporated and the residue was redissolved in dry $CH_2Cl_2$ (1 mL) and used directly in the next step.

The prepared solution of **S8** in $CH_2Cl_2$ (0.2 mL, ~0.2 mmol, ~4 equiv) was added to the stirred solution of **S14** (19 mg, 50.8 µmol) and DIPEA (50 µL, 0.29 mmol, ~6 equiv) in dry $CH_2Cl_2$ (0.5 mL), and the resulting solution was stirred at rt for 1 h. The crude reaction mixture was evaporated, the residue was dissolved in the mixture of THF (1 mL) and ethanol (0.25 mL), and lithium hydroxide solution (21 mg of $LiOH{\cdot}H_2O$ in 0.25 mL water, 0.5 mmol, 10 equiv) was added to the mixture, which was left stirring at rt for 18 h. Acetic acid (100 µL) was then added, and the reaction mixture was evaporated to dryness. The product was isolated by preparative HPLC (ThermoFisher Hypersil Gold C18 250×21.2 mm 5 µm, solvent flow rate 18 mL/min, gradient 30% to 80% A:B, A – acetonitrile + 0.1% (v/v) $HCO_2H$, B – water + 0.1% (v/v) $HCO_2H$) and freeze-dried from dioxane to give 14 mg (52% over 2 steps) of **5** as brown solid.

$^1$H NMR (400 MHz, $CDCl_3$): δ 7.55 (s, 1H), 7.05 (dd, *J* = 17.8, 11.5 Hz, 1H), 6.62 (s, 1H), 6.57 (s, 1H), 5.47 (dd, *J* = 11.5, 1.6 Hz, 1H), 5.34 (dd, *J* = 17.8, 1.6 Hz, 1H), 4.00 (t, *J* = 6.1 Hz, 2H), 3.47 – 3.34 (m, 4H), 3.06 (t, *J* = 8.1 Hz, 2H), 2.99 (t, *J* = 8.2 Hz, 2H), 2.85 (s, 6H), 2.26 (t, *J* = 7.4 Hz, 2H), 1.56 – 1.44 (m, 4H), 1.16 – 1.05 (m, 2H).

$^{13}$C NMR (101 MHz, $CDCl_3$): 135.3, 123.7, 118.0, 110.0, 109.1, 65.5, 55.5, 35.5, 33.9, 29.0, 28.2, 28.1, 25.2, 24.5, -2.4 (indirect detection from a gHSQC experiment, only H-coupled carbons are resolved).

HRMS ($C_{30}H_{37}N_3O_4Si$): *m/z* (positive mode) = 532.2647 (found $[M+H]^+$), 532.2626 (calc.).

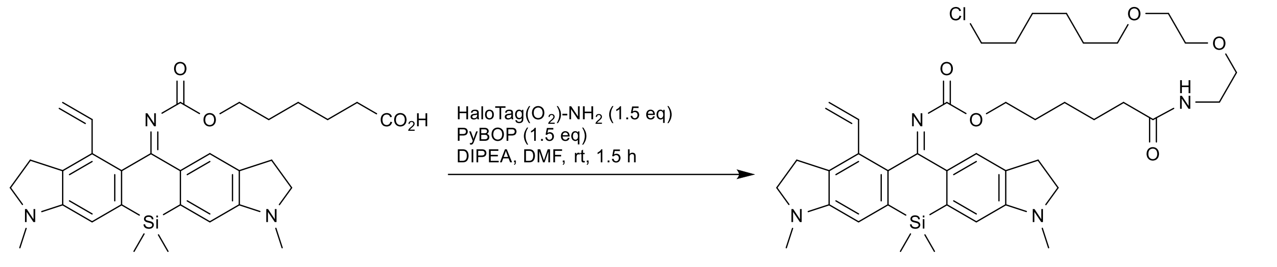

**Compound 5-Halo.** PyBOP solution (benzotriazol-1-yloxy-tris(pyrrolidino)phosphonium hexafluorophosphate; 10.4 mg in 100 µL DMF, 20.0 µmol, 1.5 equiv) was added to the stirred solution of **5** (7 mg, 13.2 µmol) and HaloTag(O2) amine hydrochloride (5.2 mg, 20.0 µmol, 1.5 equiv) in DMF (150 µL) and DIPEA (50 µL), and the reaction mixture was stirred at rt for 1.5 h. The organic solvents were evaporated *in vacuo*, and the product was isolated by preparative HPLC (ThermoFisher Hypersil Gold C18 250×21.2 mm 5 µm, solvent flow rate 18 mL/min, gradient 30% to 90% A:B, A – acetonitrile + 0.1% (v/v) $HCO_2H$, B – water + 0.1% (v/v) $HCO_2H$) to give 8.3 mg (86%) of **5-Halo** as viscous brown-yellow oil.

$^1$H NMR (400 MHz, $CDCl_3$): δ 7.61 (br.s, 1H), 7.05 (dd, *J* = 17.8, 11.5 Hz, 1H), 6.62 (s, 1H), 6.59 (s, 1H), 6.26 (br.s, 1H), 5.47 (dd, *J* = 11.5, 1.5 Hz, 1H), 5.34 (dd, *J* = 17.8, 1.5 Hz, 1H), 4.01 (t, *J* = 6.3 Hz, 2H), 3.65 – 3.50 (m, 8H), 3.50 – 3.34 (m, 8H), 3.07 (t, *J* = 8.1 Hz, 2H), 3.00 (t, *J* = 8.2 Hz, 2H), 2.87 (s, 6H), 2.10 (t, *J* = 7.7 Hz, 2H), 1.82 – 1.71 (m, 2H), 1.66 – 1.32 (m, 10H), 1.22 – 1.08 (m, 2H), 0.47 (s, 6H).

$^{13}$C NMR (101 MHz, $CDCl_3$): δ 135.2, 118.1, 109.9, 109.4, 71.4, 70.3, 70.1, 65.7, 55.6, 45.1, 39.3, 36.6, 35.6, 32.7, 29.6, 29.1, 28.5, 28.3, 26.7, 25.6, 25.4, -2.3 (indirect detection from a gHSQC experiment, only H-coupled carbons are resolved).

HRMS ($C_{40}H_{57}ClN_4O_5Si$): *m/z* (positive mode) = 737.3844 (found $[M+H]^+$), 737.3860 (calc.).

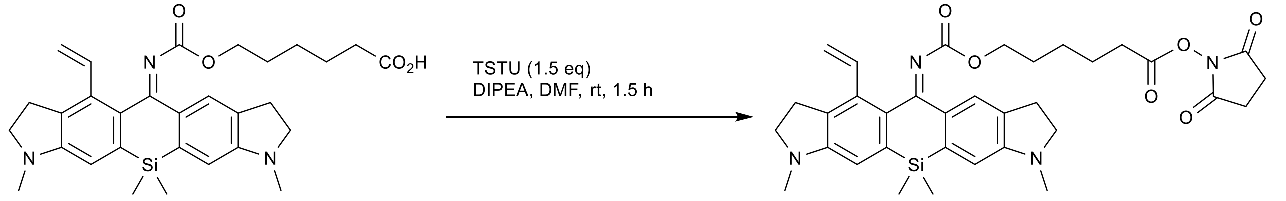

**Compound 5-NHS.** TSTU solution (*N*,*N*,*N'*,*N'*-tetramethyl-*O*-(*N*-succinimidyl)uronium tetrafluoroborate; 5.2 mg in 50 µL DMF, 17.2 µmol, 1.5 equiv) was added to the stirred solution of **5** (6.1 mg, 11.5 µmol) in DMF (150 µL) and DIPEA (50 µL), and the reaction mixture was stirred at rt for 1.5 h. The organic solvents were evaporated *in vacuo*, and the product was isolated by flash column chromatography (12 g Interchim SiHP 30 µm cartridge, gradient 30% to 100% EtOAc/hexane) and freeze-dried from 1,4-dioxane to yield 6.4 mg (89%) of **5-NHS** as yellow solid.

HRMS ($C_{34}H_{40}N_4O_6Si$): *m/z* (positive mode) = 629.2782 (found $[M+H]^+$), 629.2790 (calc.).

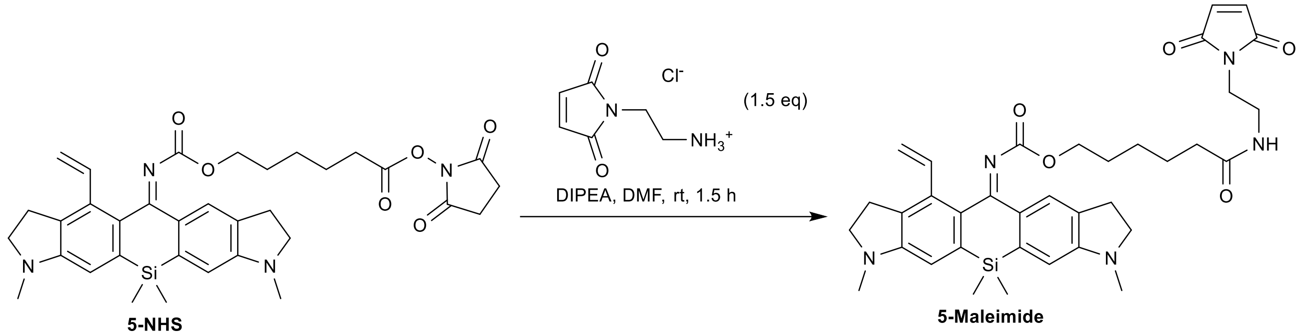

**Compound 5-Maleimide.** A solution of **5-NHS** (5.7 mg, 9.1 µmol), 1-(2-aminoethyl)maleimide hydrochloride (2.4 mg, 13.6 µmol, 1.5 equiv) in DMF (200 µL) and DIPEA (50 µL) was stirred at rt for 1.5 h. The organic solvents were evaporated *in vacuo*, and the product was isolated by preparative HPLC (ThermoFisher Hypersil Gold C18 250×21.2 mm 5 µm, solvent flow rate 18 mL/min, gradient 30% to 80% A:B, A – acetonitrile + 0.1% (v/v) $HCO_2H$, B – water + 0.1% (v/v) $HCO_2H$) to give 2.1 mg (35%) of **5-Mal** as yellow solid.

HRMS ($C_{36}H_{43}N_5O_5Si$): *m/z* (positive mode) = 654.3101 (found $[M+H]^+$), 654.3106 (calc.).

### NMR SPECTRA

### Compound S2 $^1$H

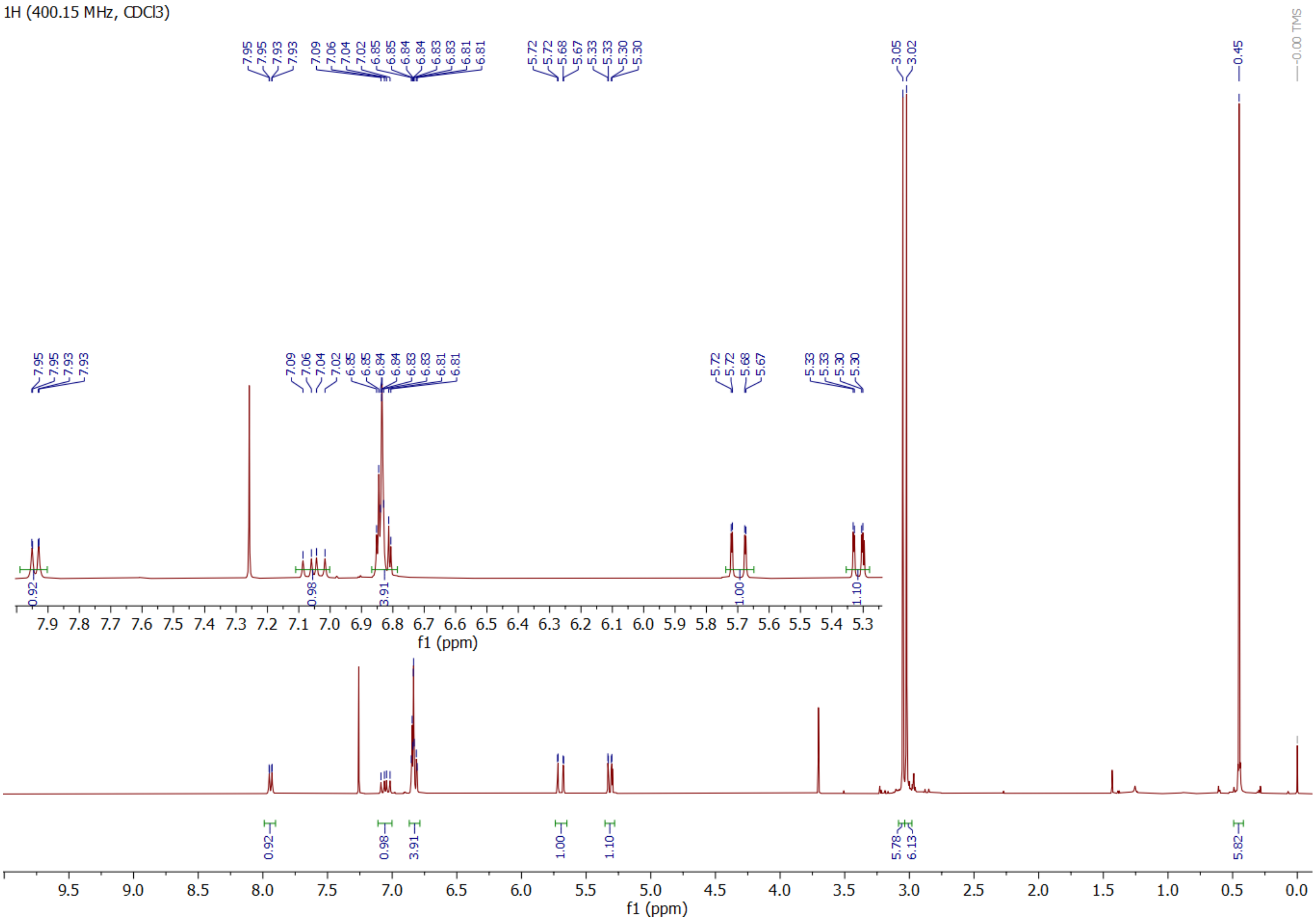

**Compound S2 $^{13}$C**

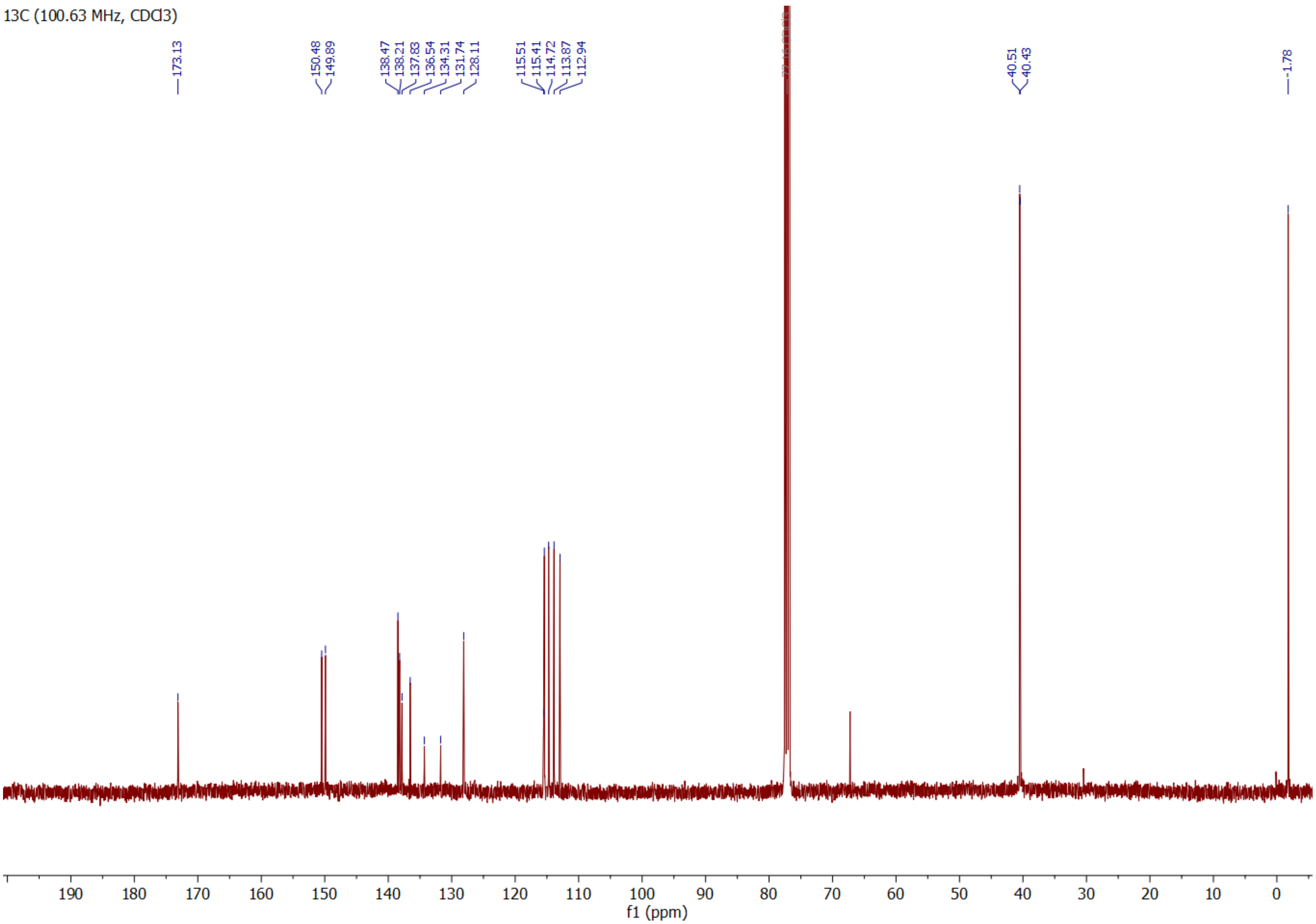

**Compound S3 $^1$H**

1H (400.15 MHz, CD3CN)

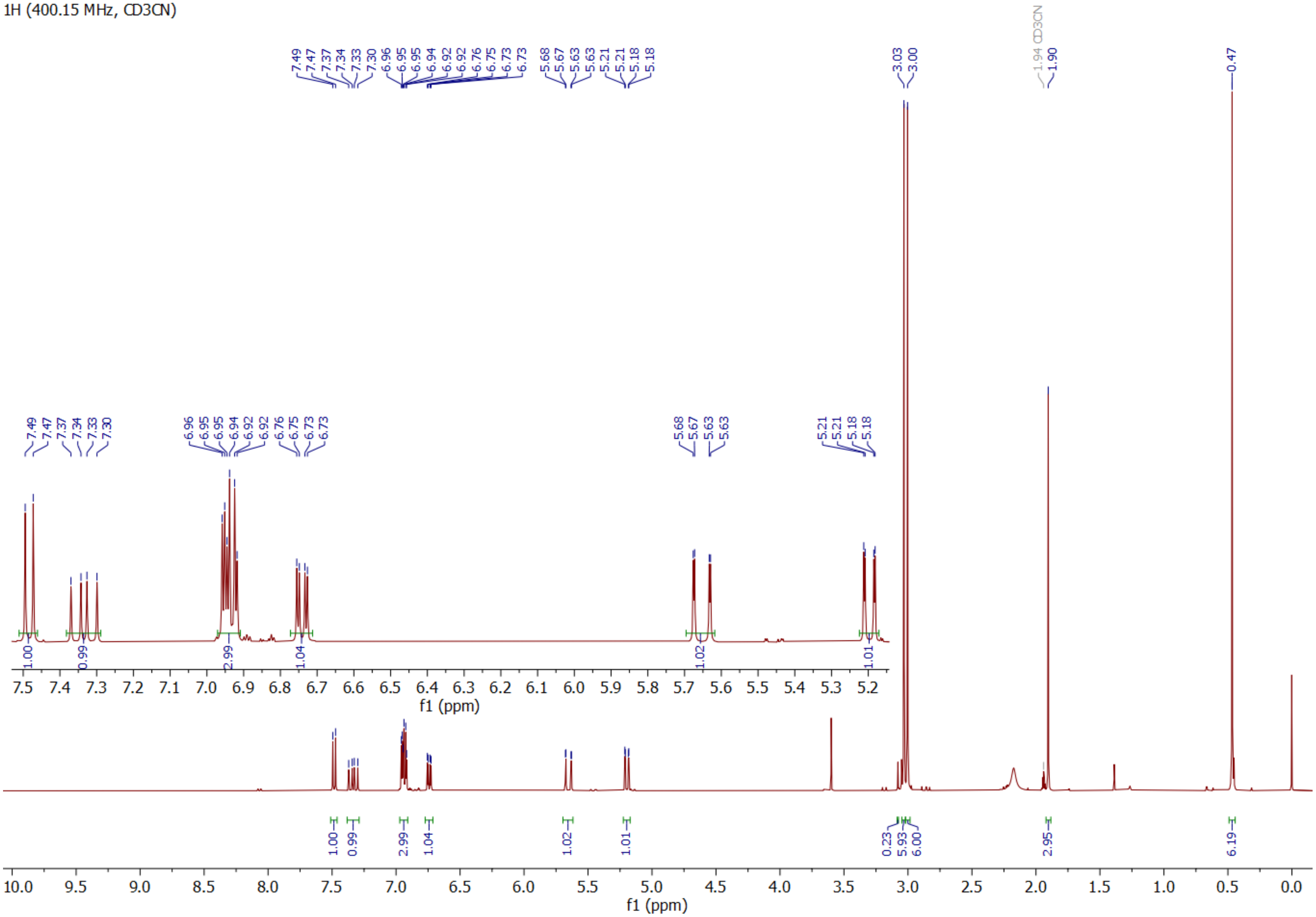

**Compound S3 $^{13}$C**

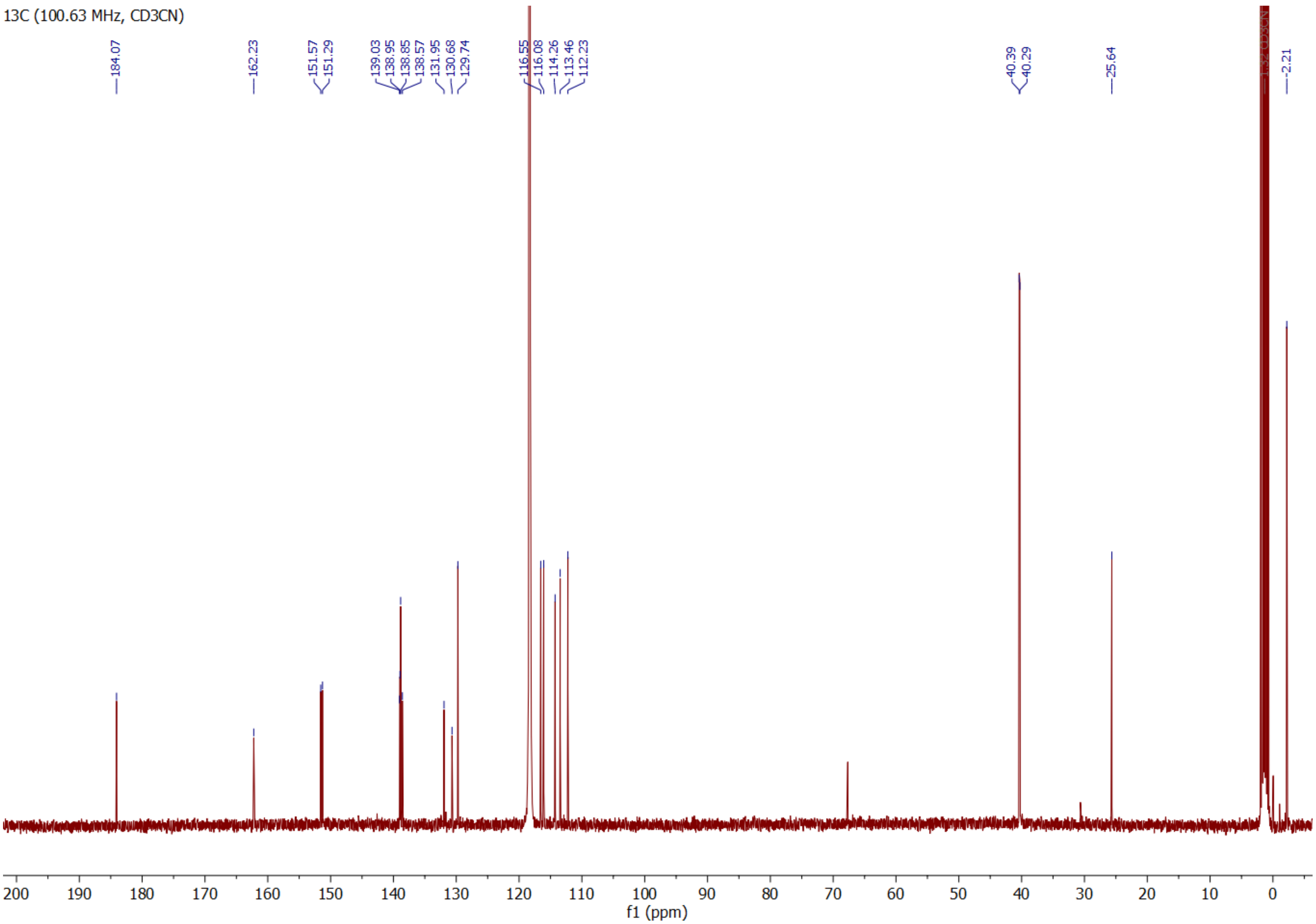

**Compound 1-CF* ¹H**

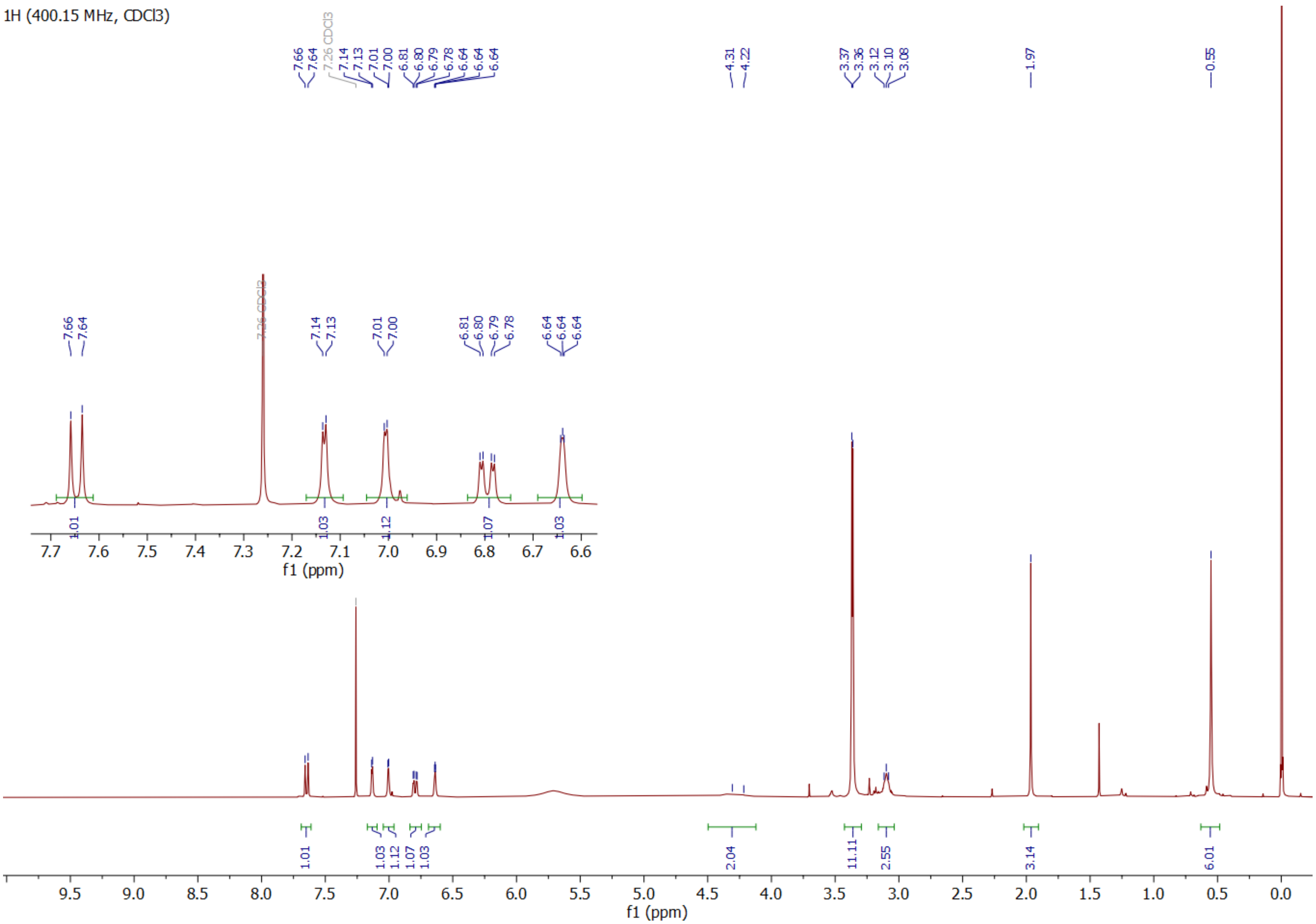

**Compound 1-CF* gHSQCad**

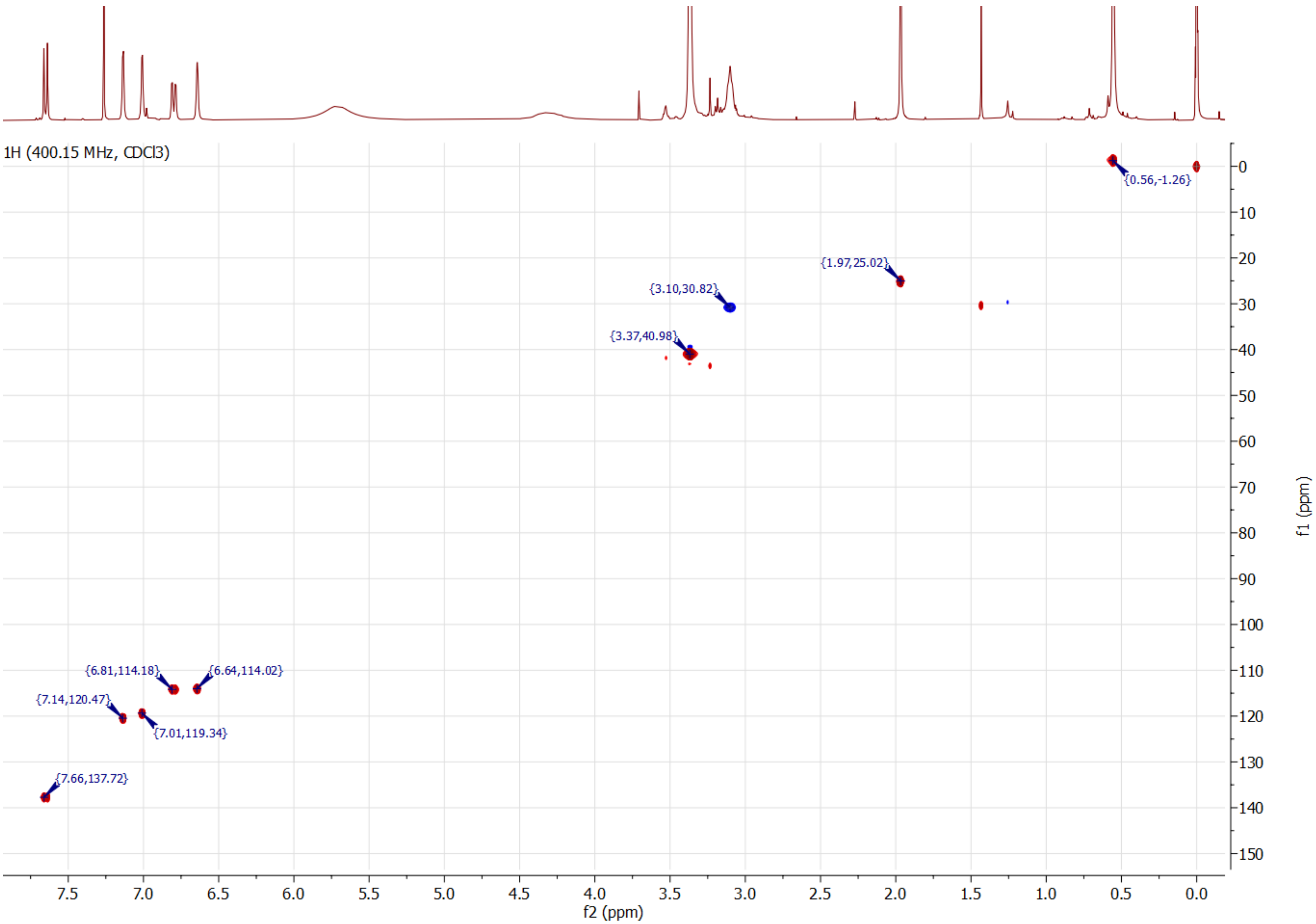

**Compound S4 ¹H**

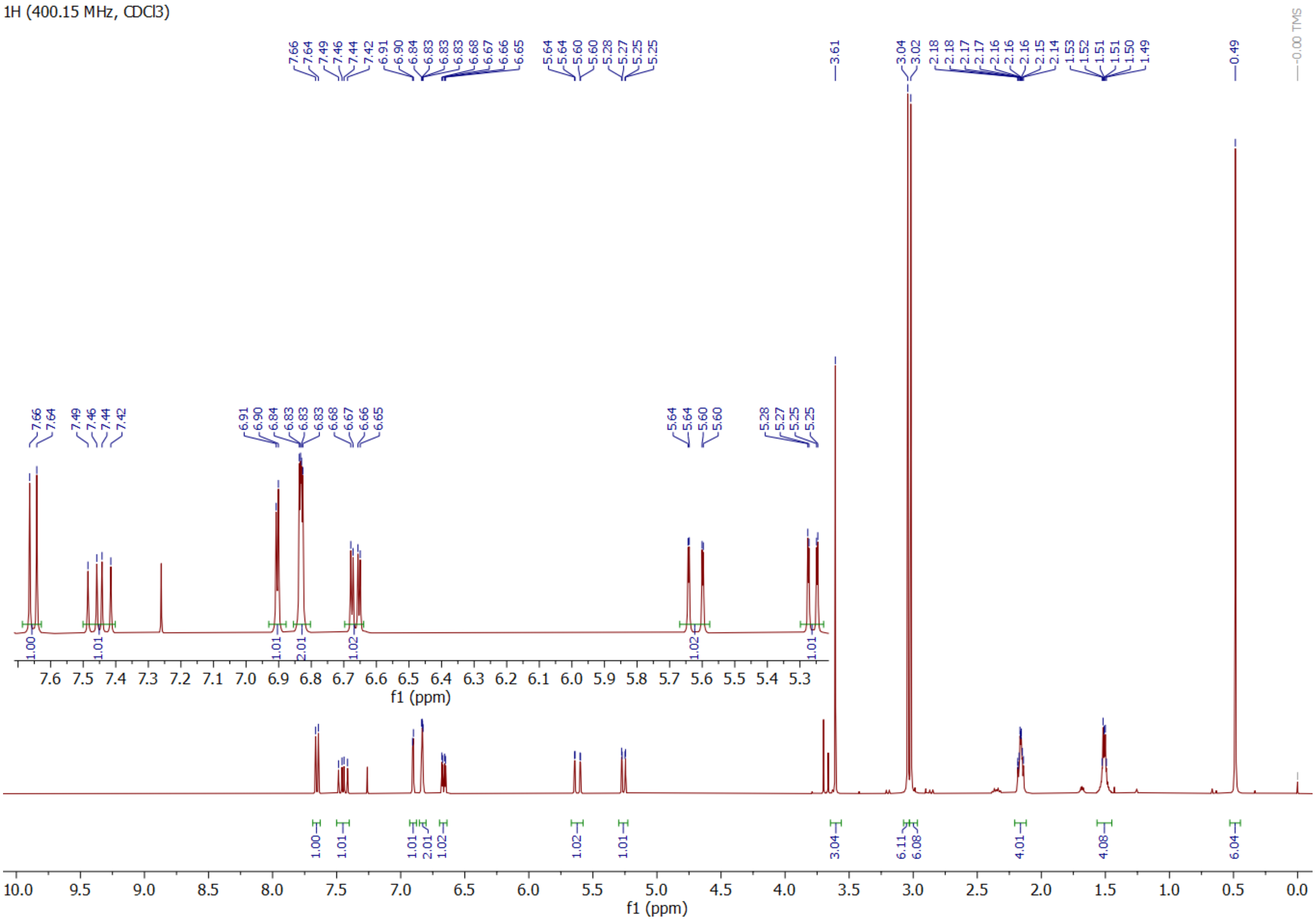

**Compound S4 $^{13}C$**

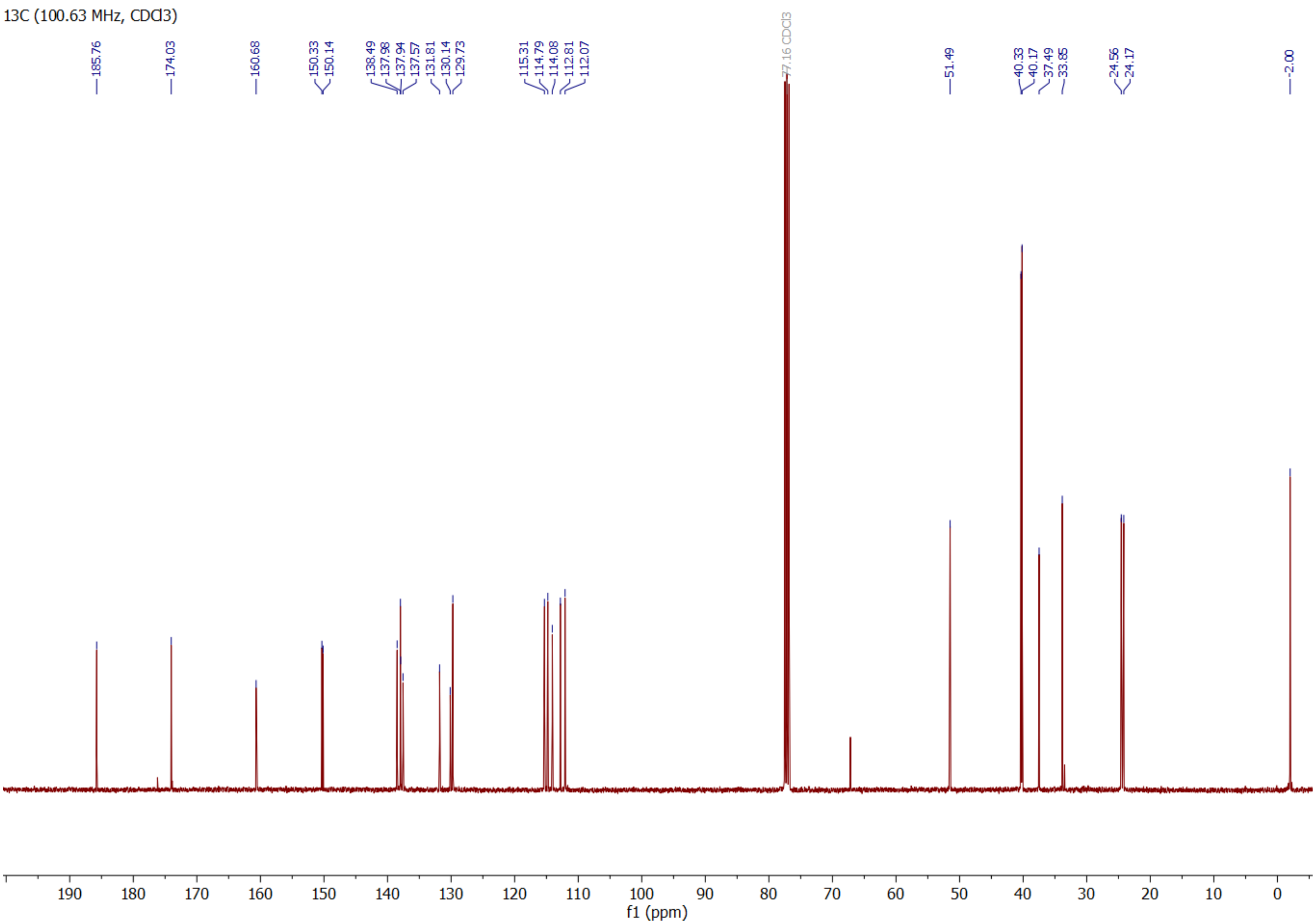

**Compound 1 $^{1}$H**

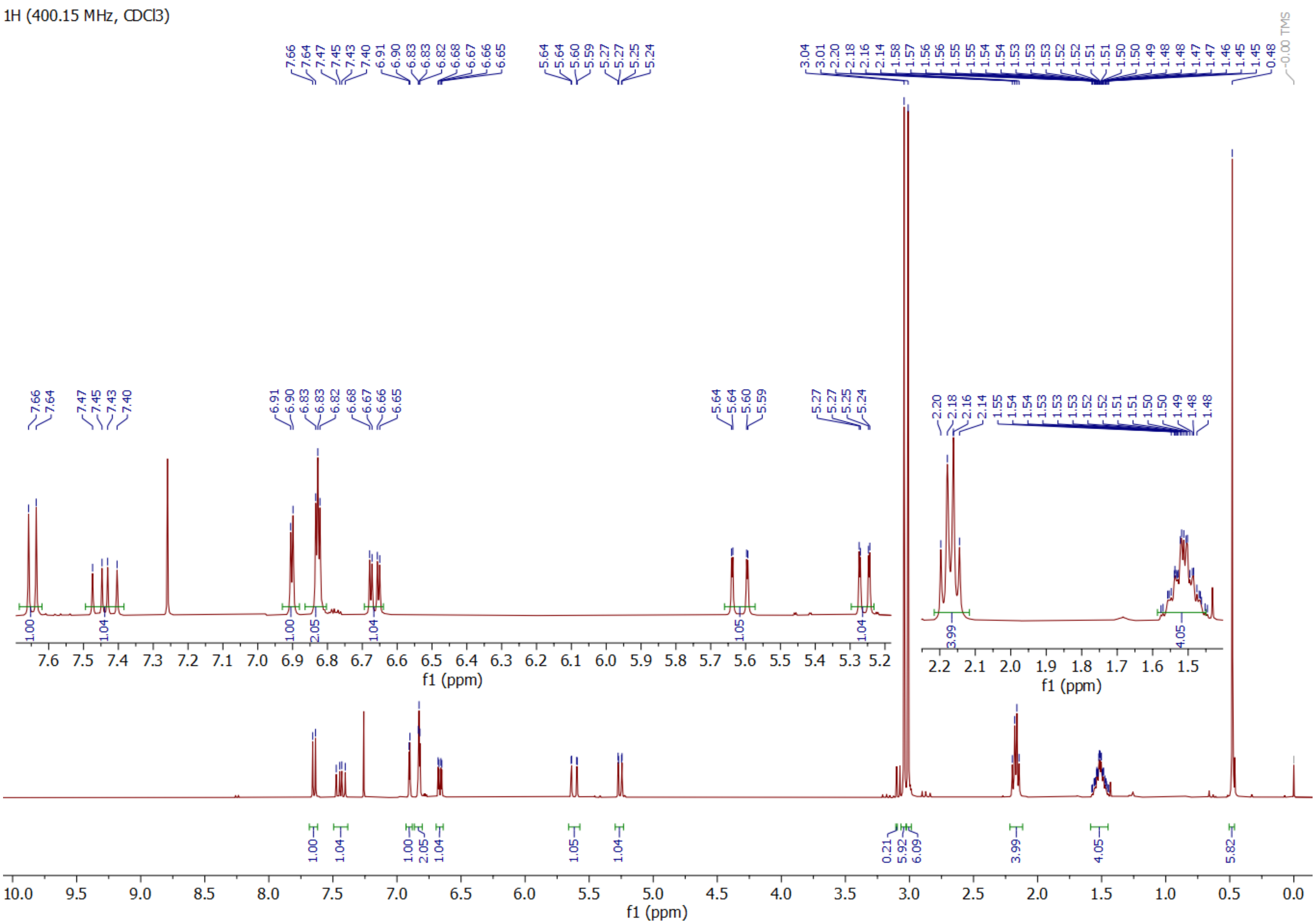

**Compound 1 $^{13}C$**

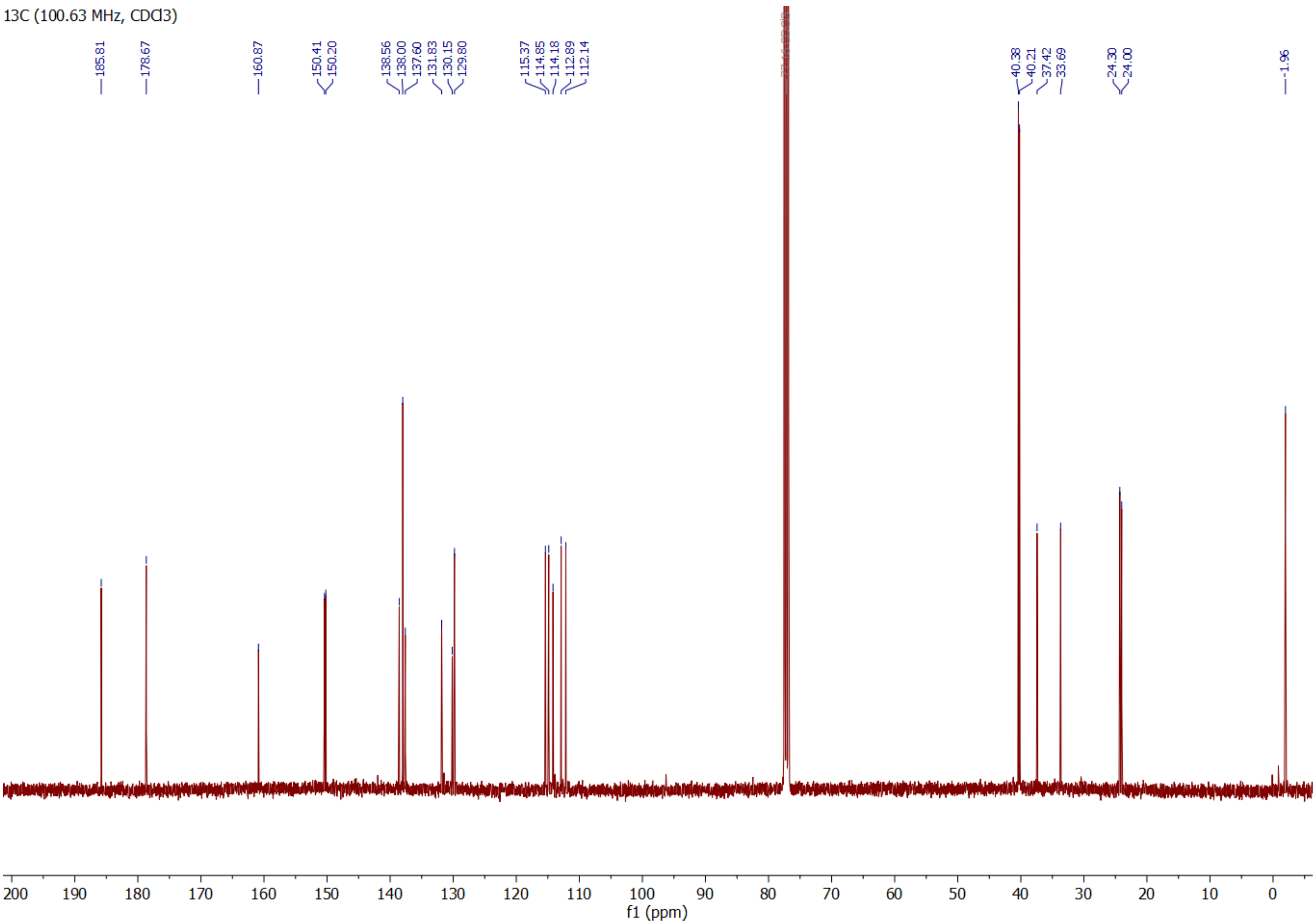

**Compound S6 $^{1}$H**

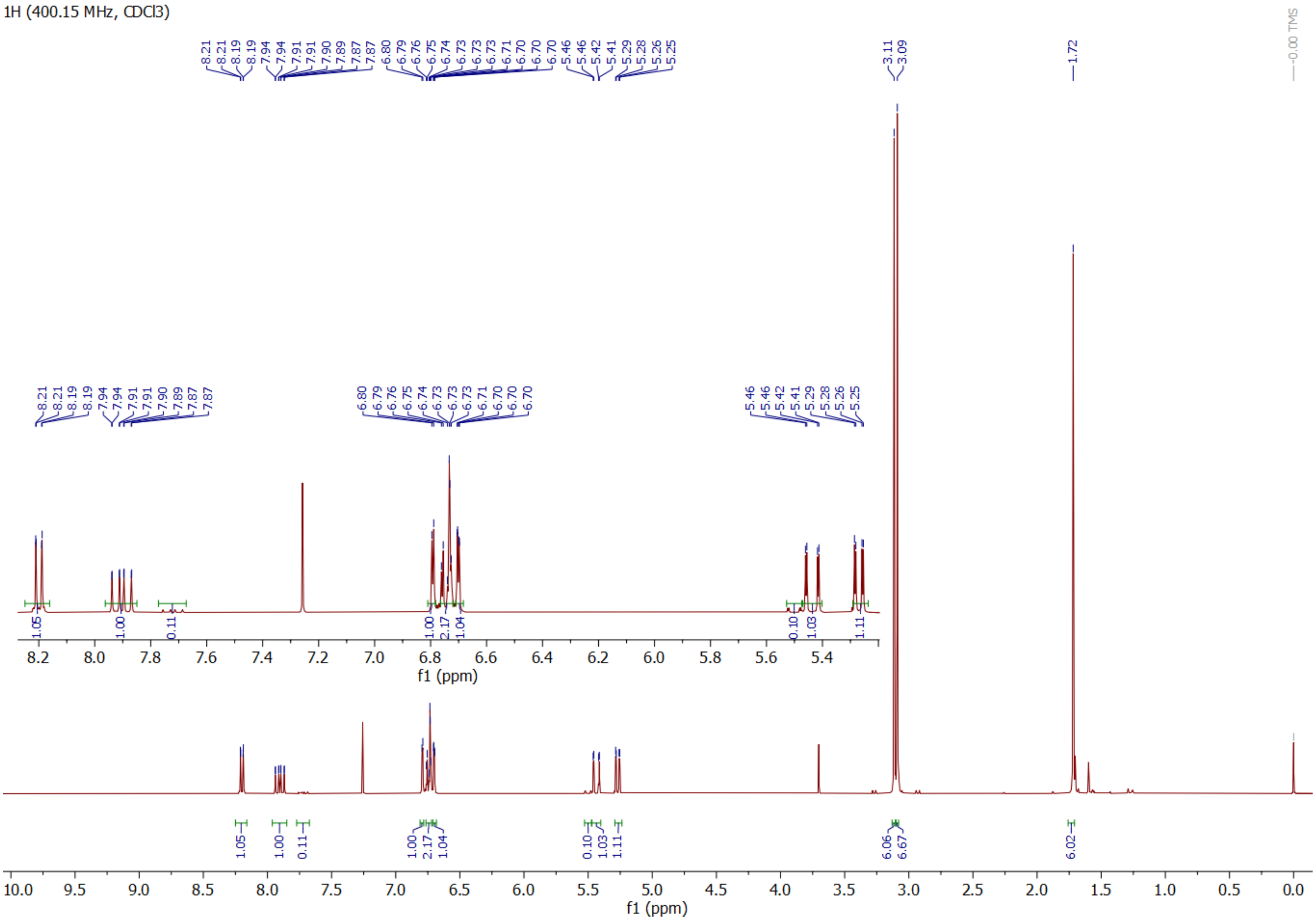

**Compound S6 $^{13}C$**

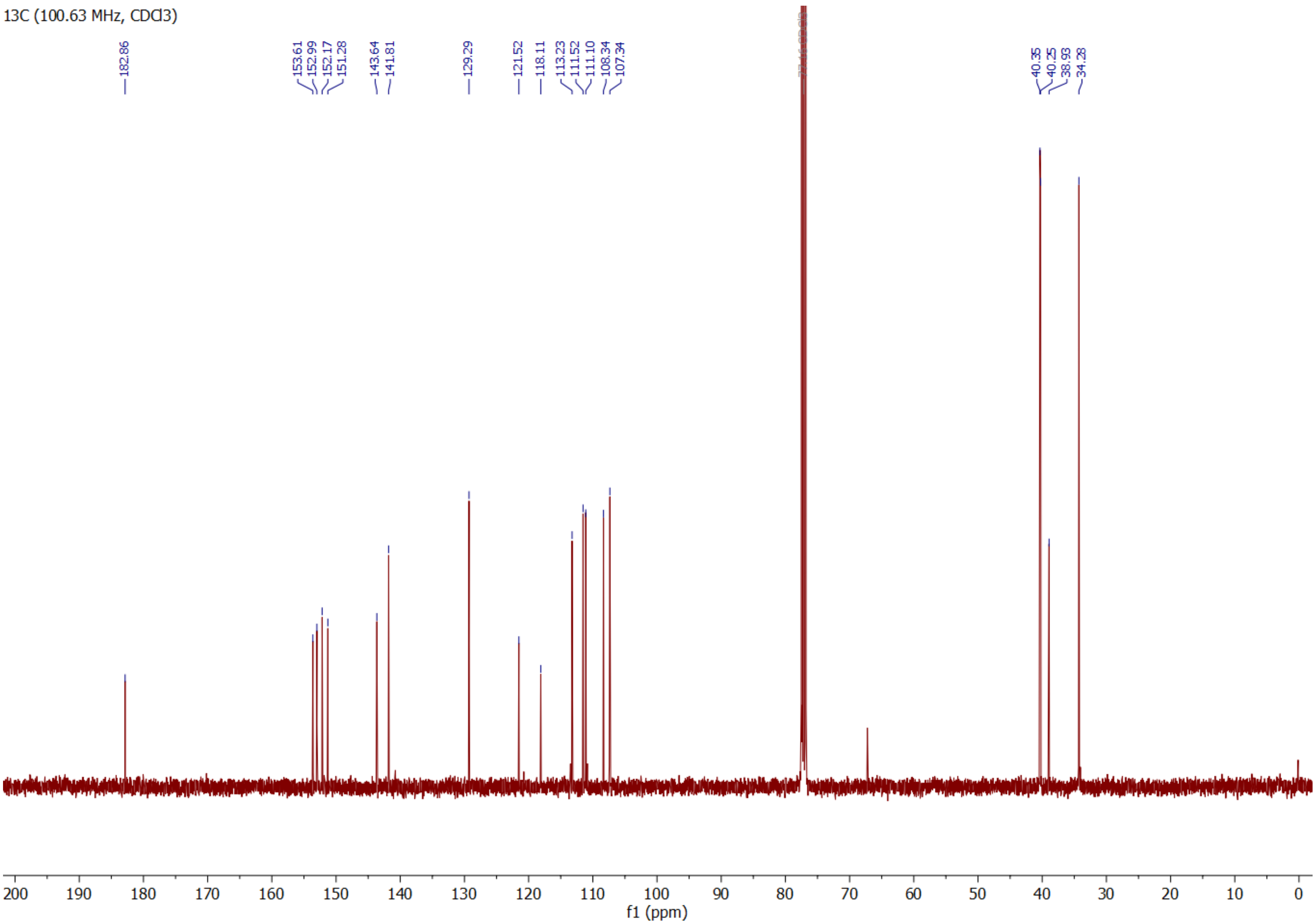

**Compound S7 $^{1}$H**

1H (400.15 MHz, CDCl3)

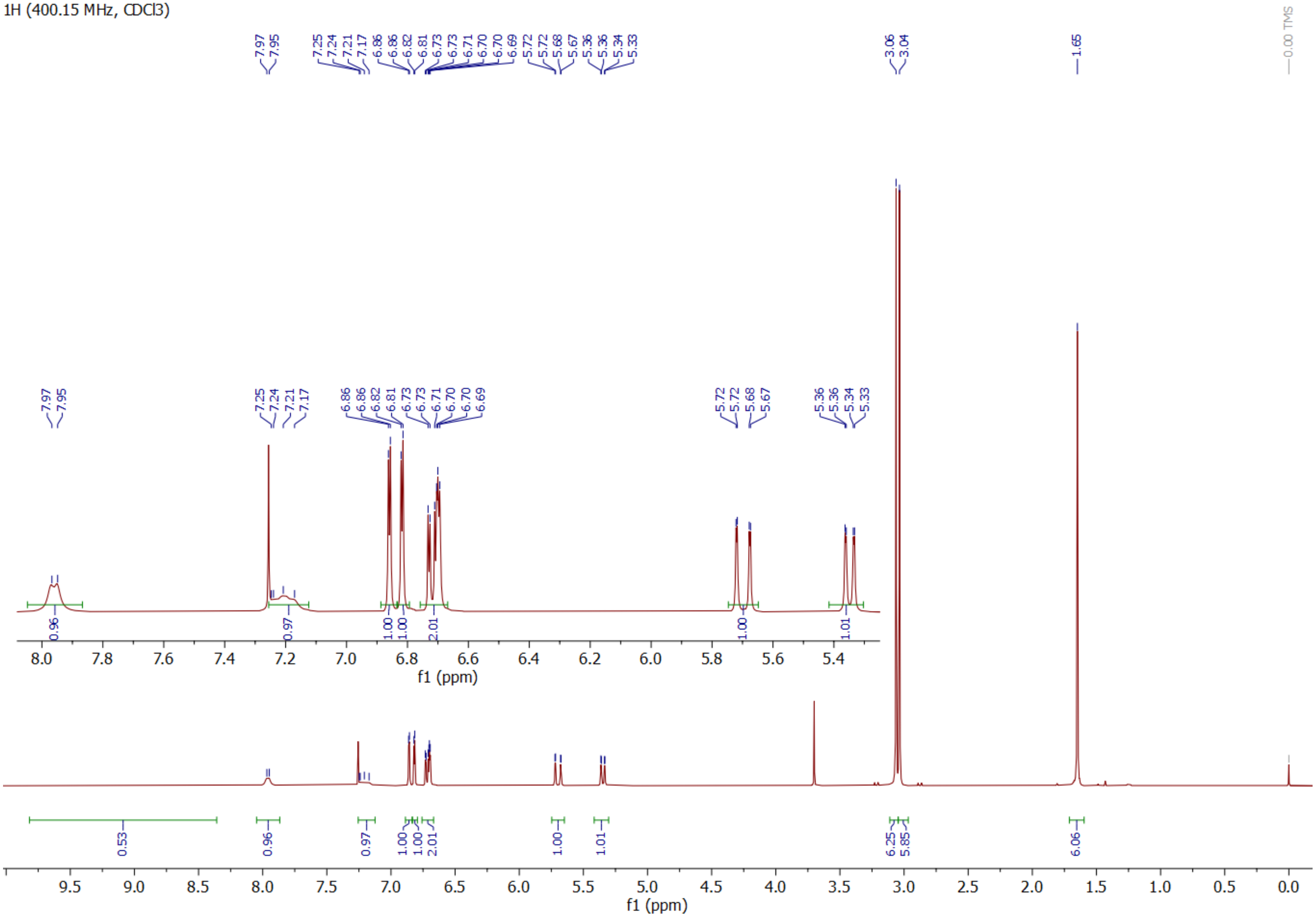

**Compound S7 $^{13}C$**

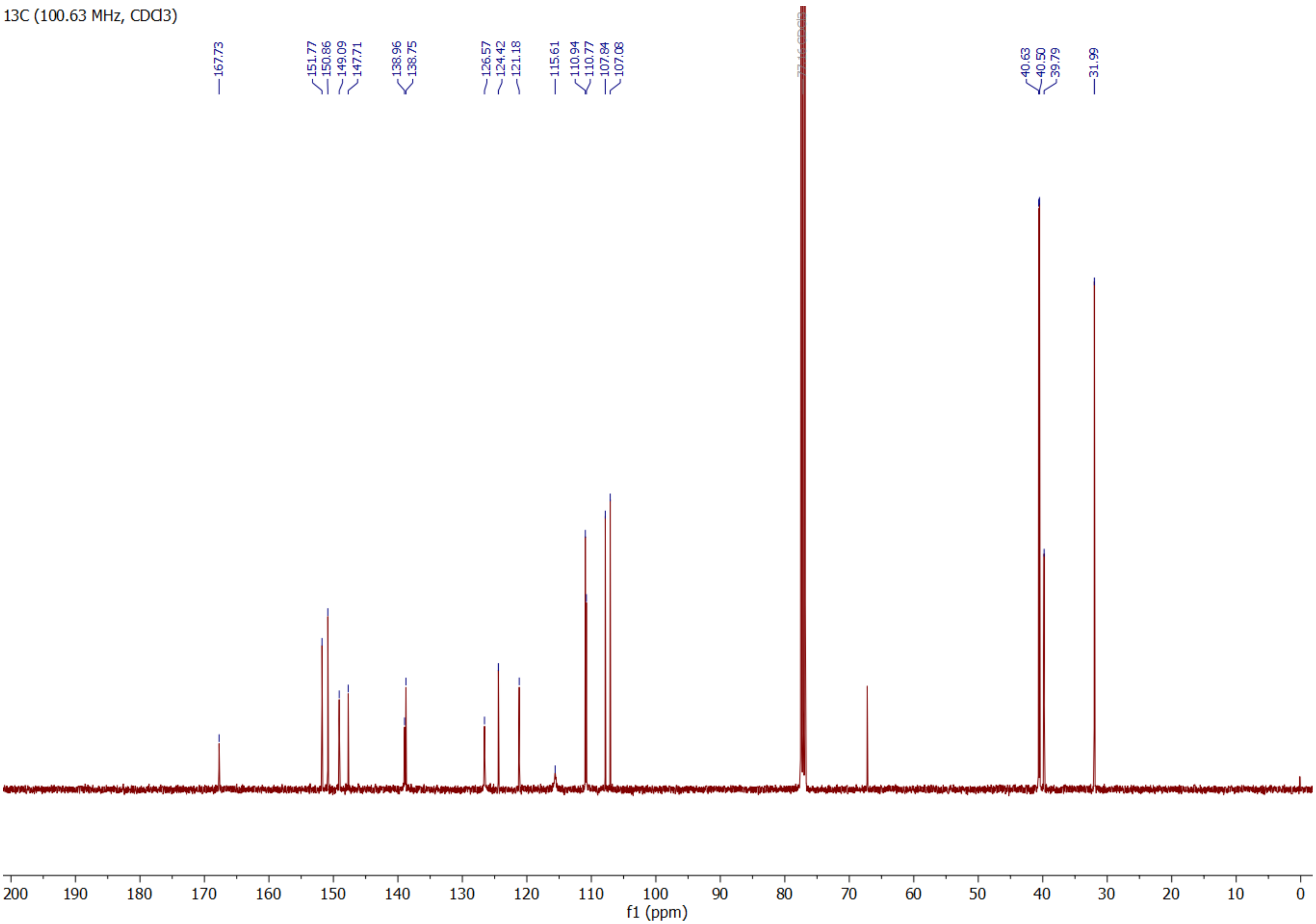

**Compound S8 $^{1}$H**

1H (400.15 MHz, CDCl3)

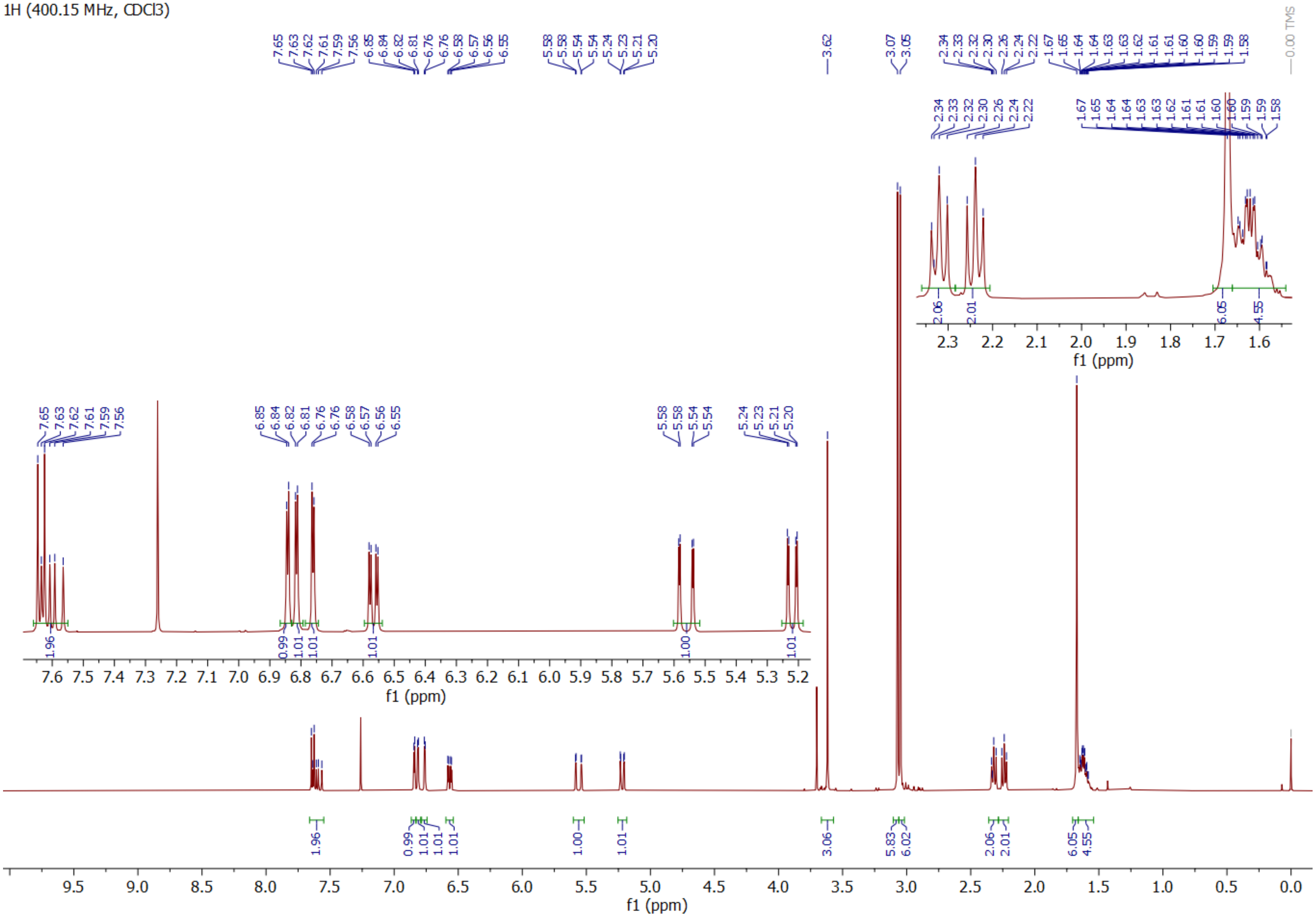

**Compound S8 $^{13}C$**

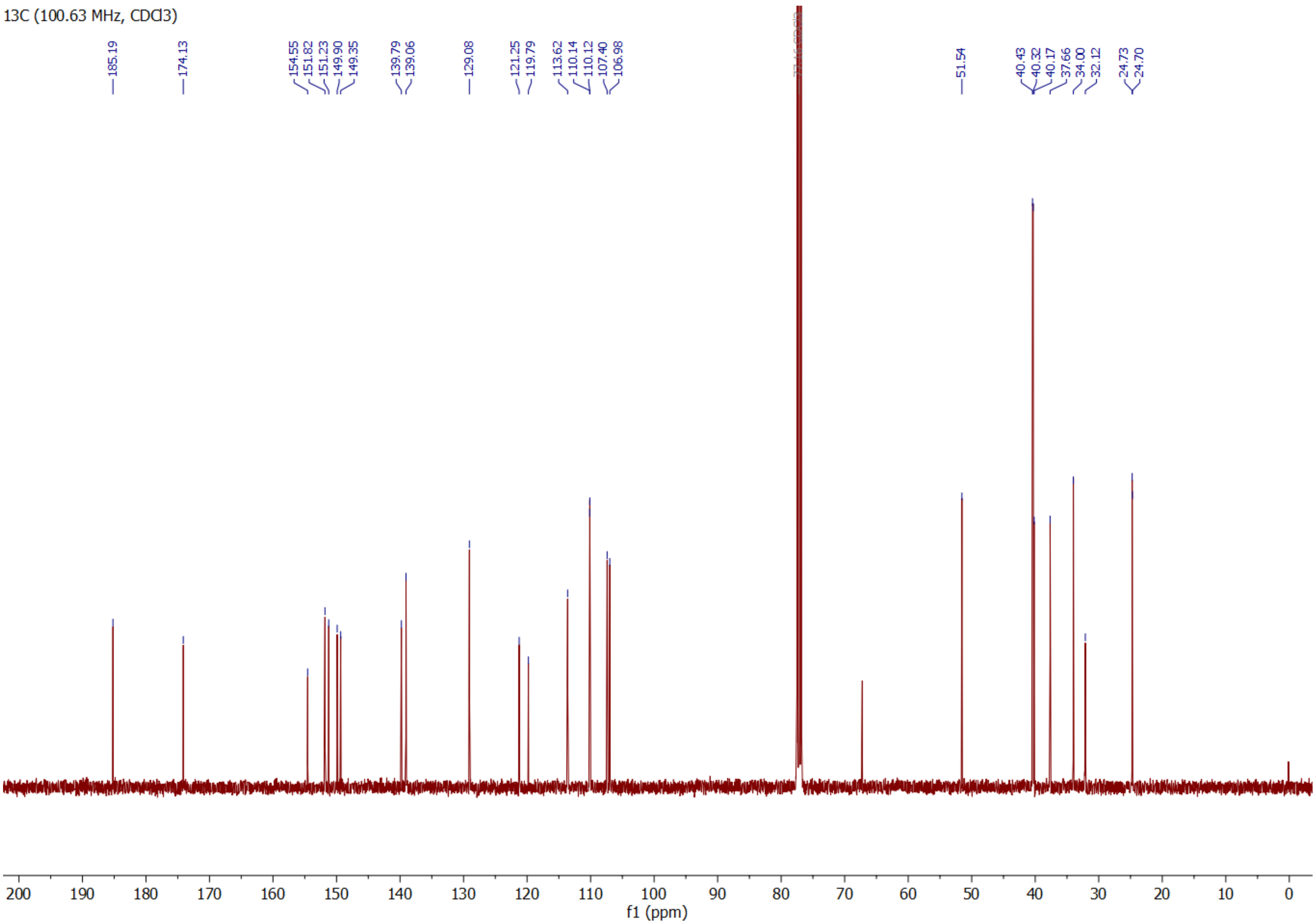

**Compound 2 $^{1}$H**

1H (400.15 MHz, CD3CN)

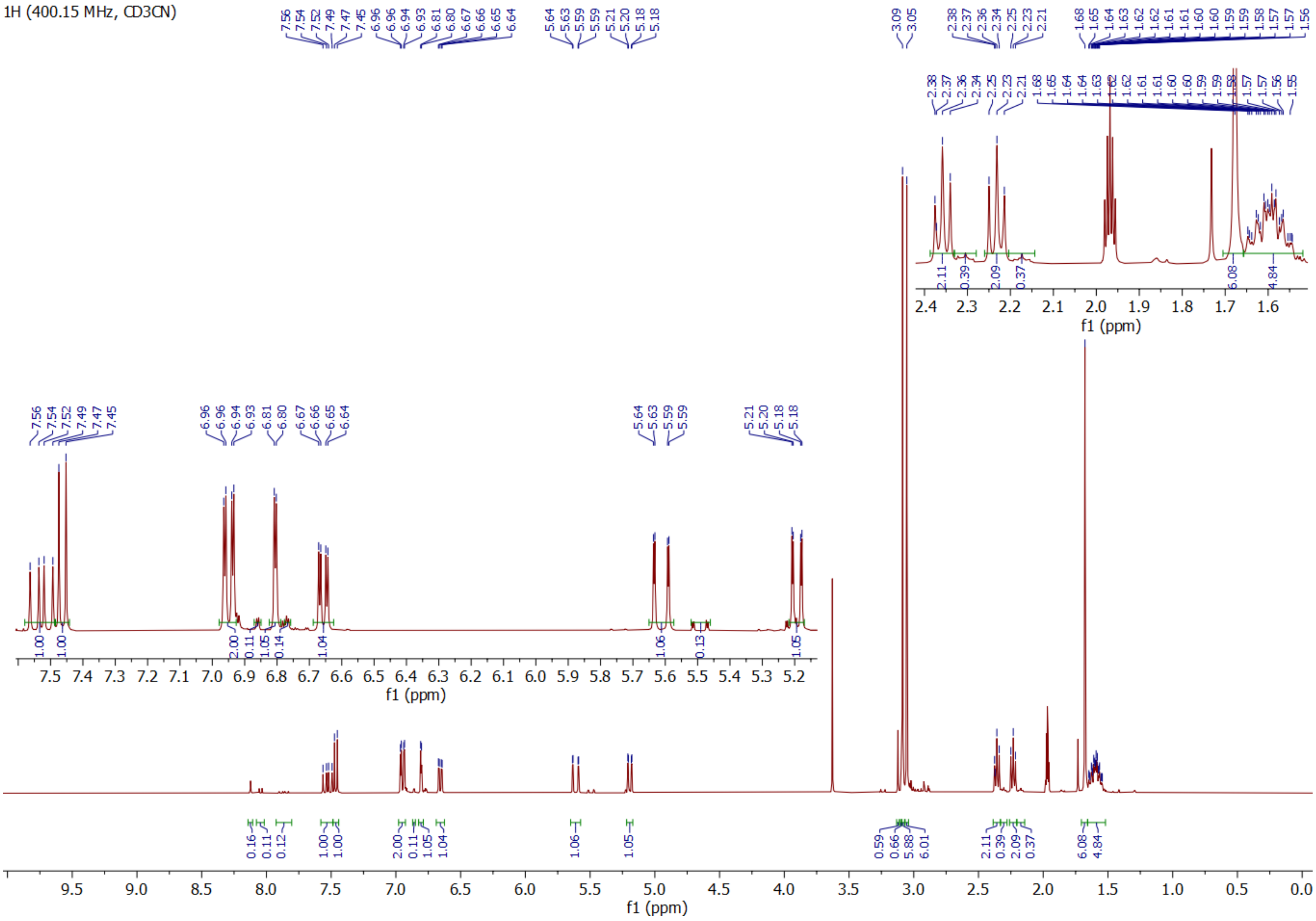

**Compound 2 $^{13}C$**

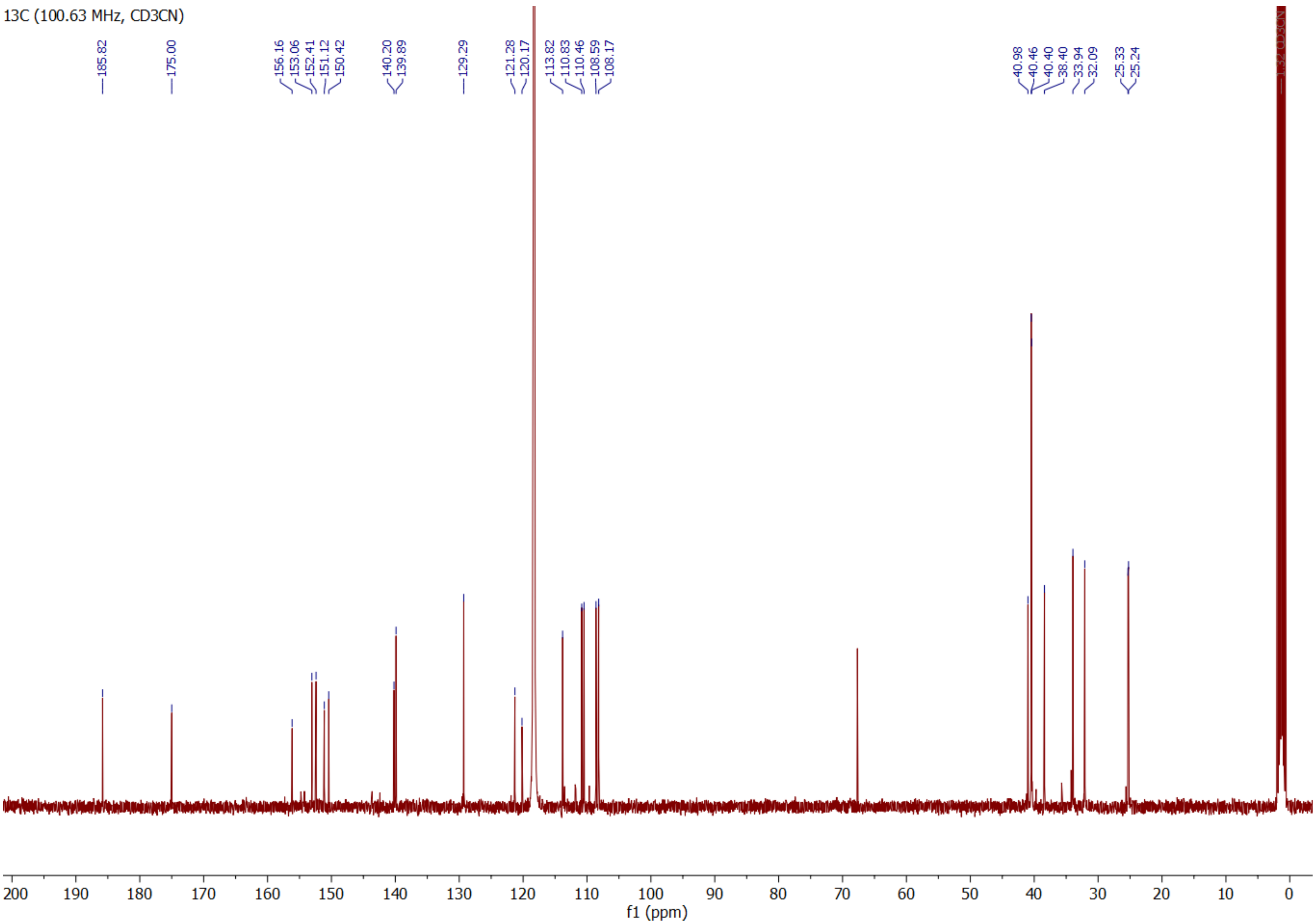

**Compound 3 $^1$H**

1H (400.15 MHz, DMSO)

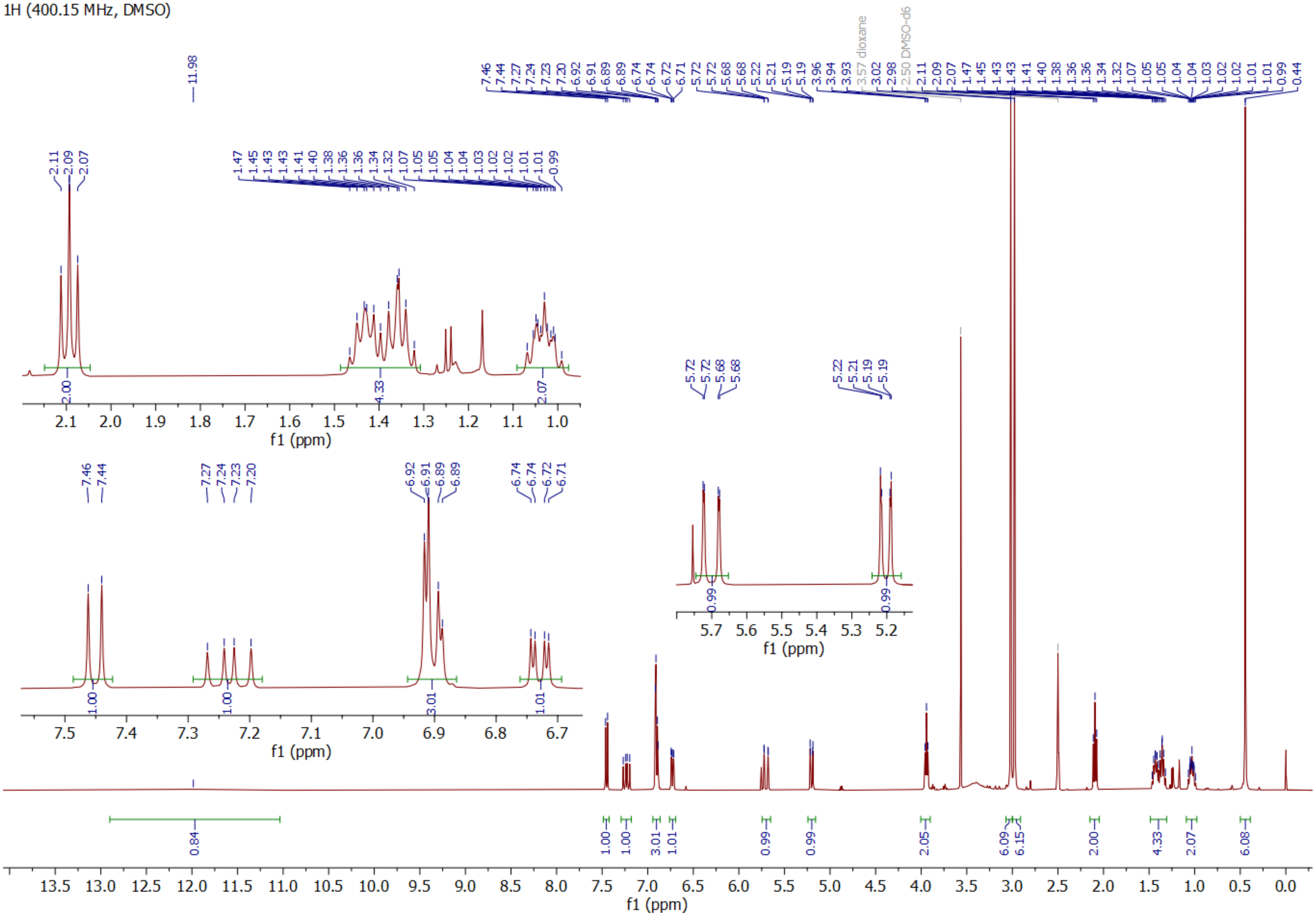

**Compound 3 $^{13}C$**

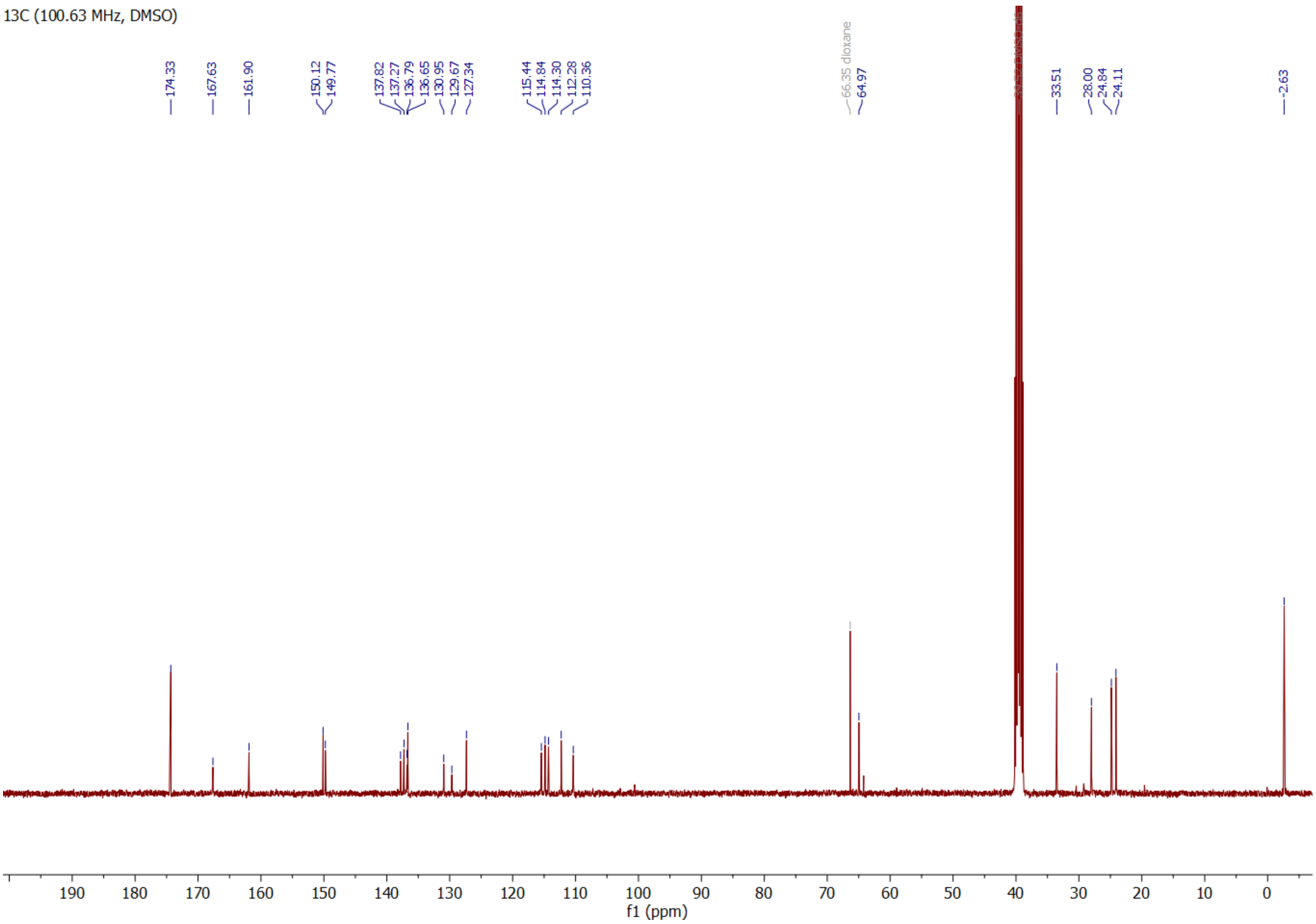

**Compound 4 $^{1}$H**

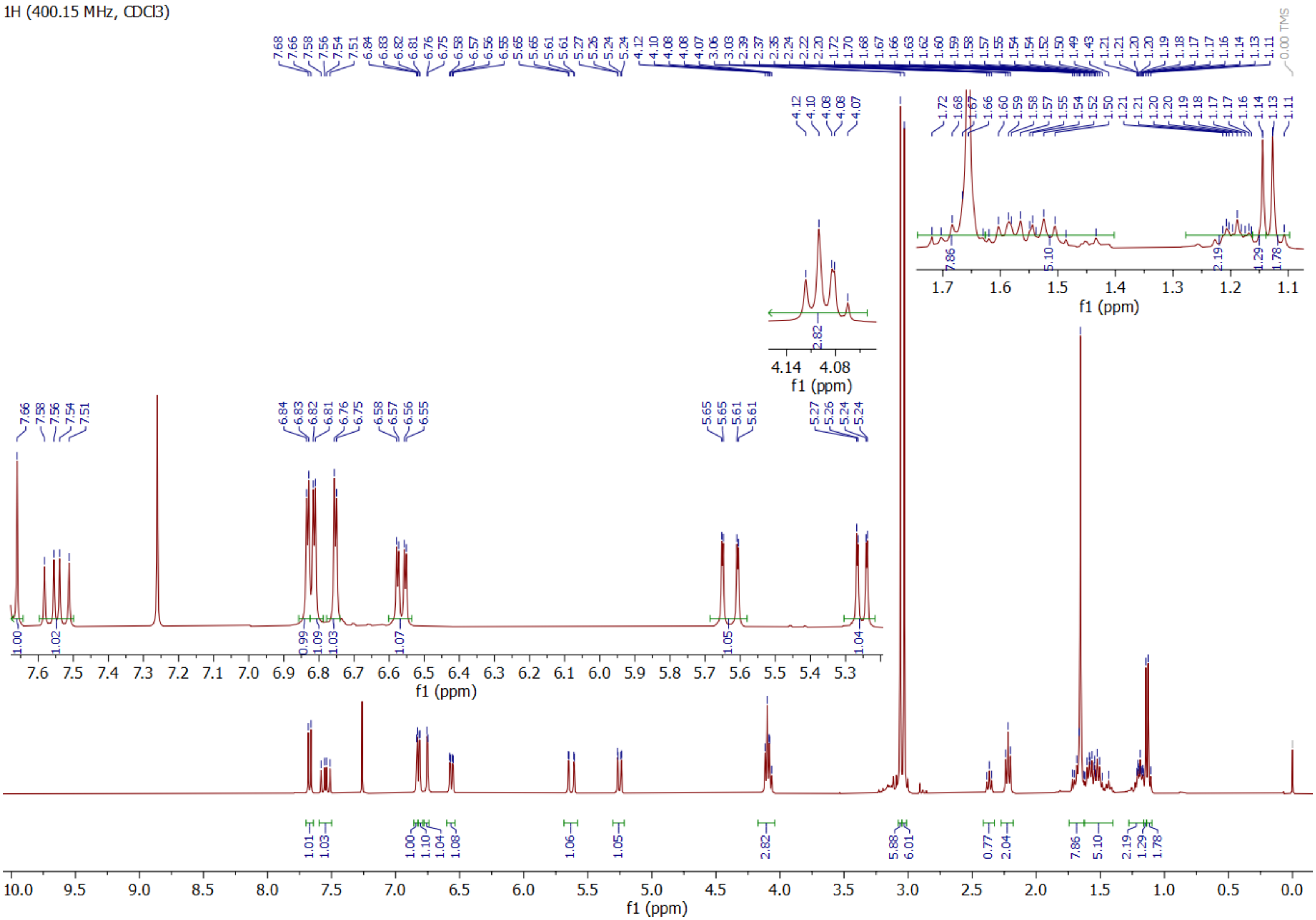

**Compound 4 $^{13}C$**

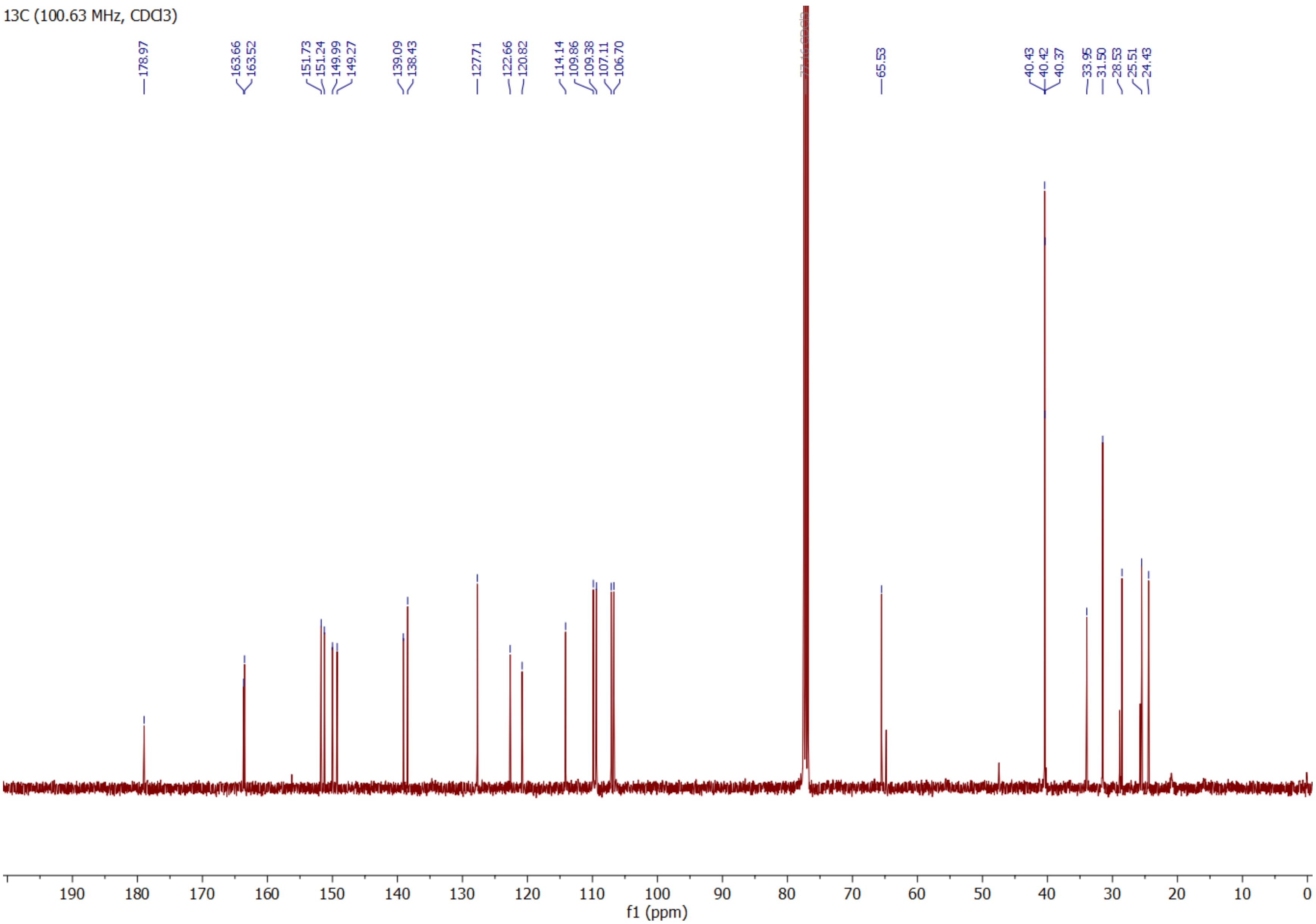

**Compound 1-Maleimide $^1$H**

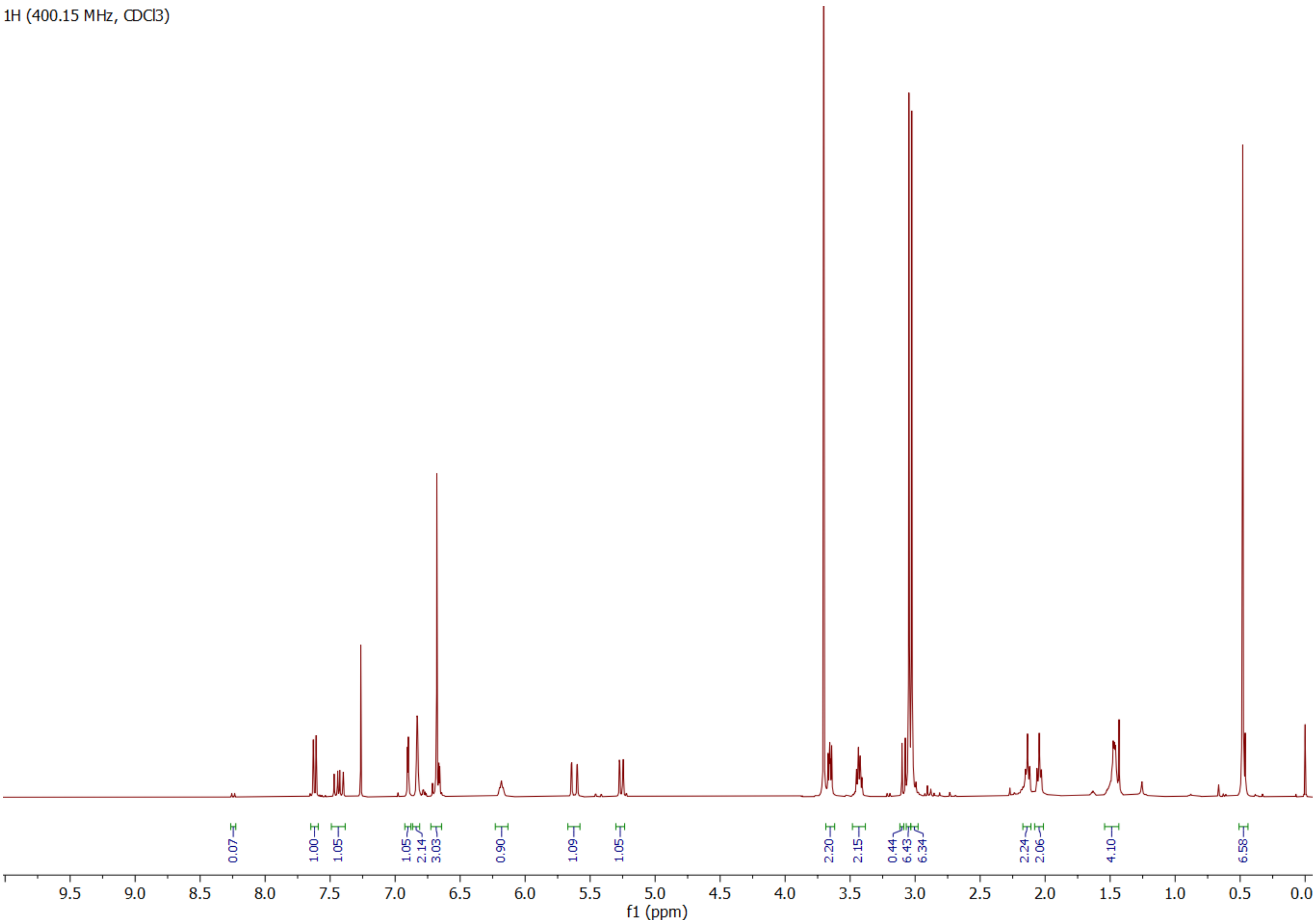

**Compound 1-Maleimide $^{13}C$**

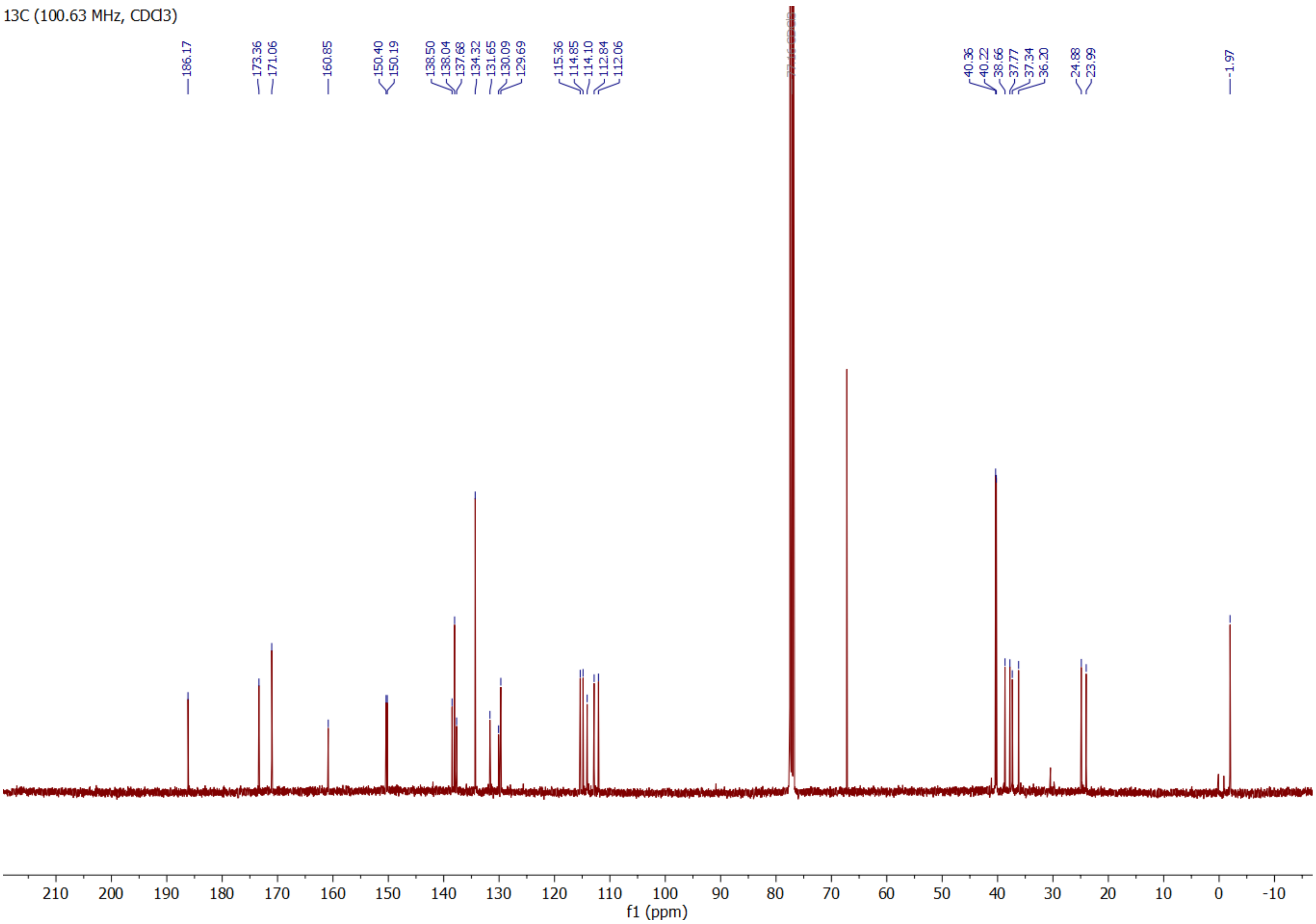

**Compound 3-Halo $^1$H**

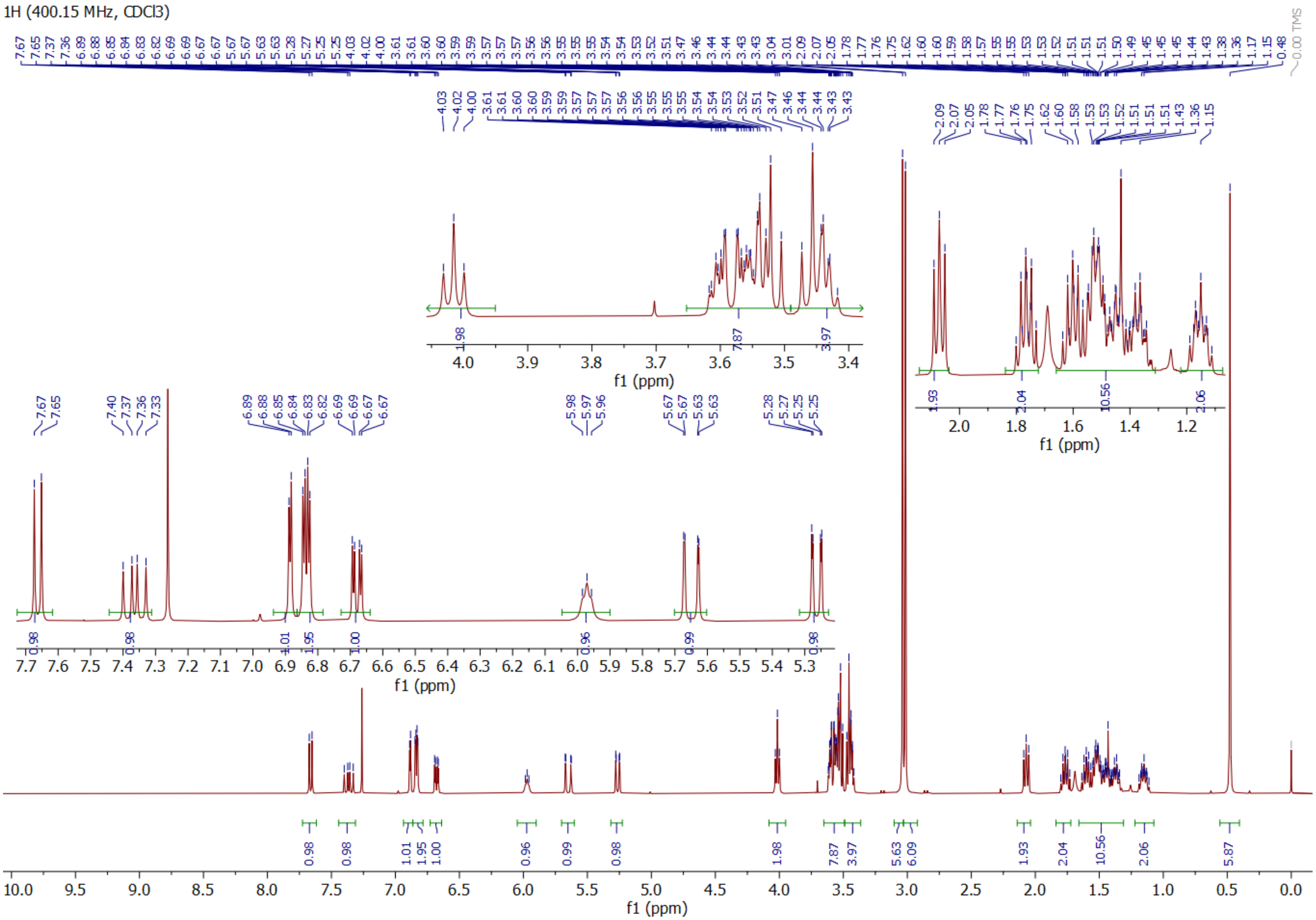

**Compound 3-Halo $^{13}C$**

13C (100.63 MHz, CDCl3)

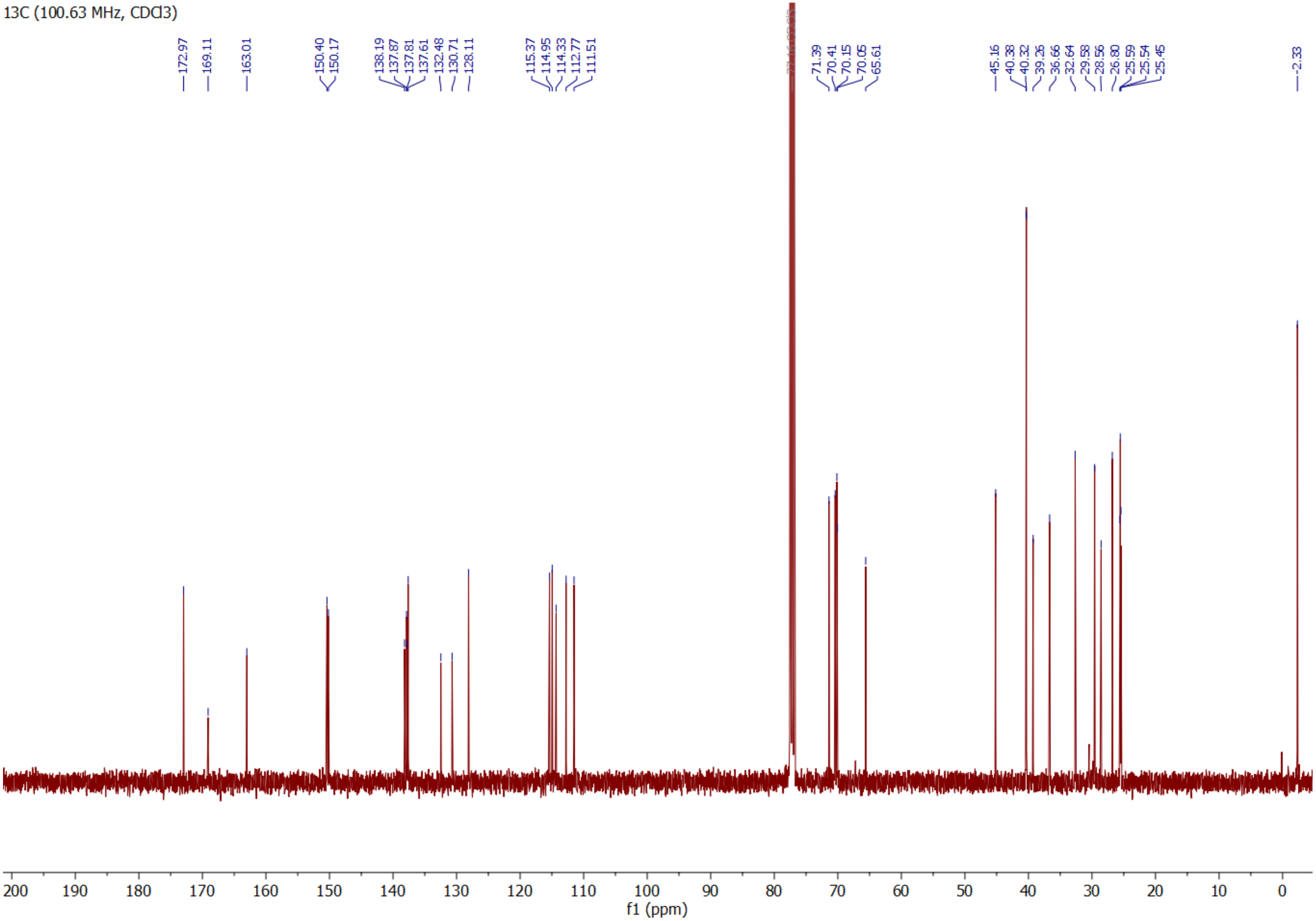

Compound S13 $^{1}$H

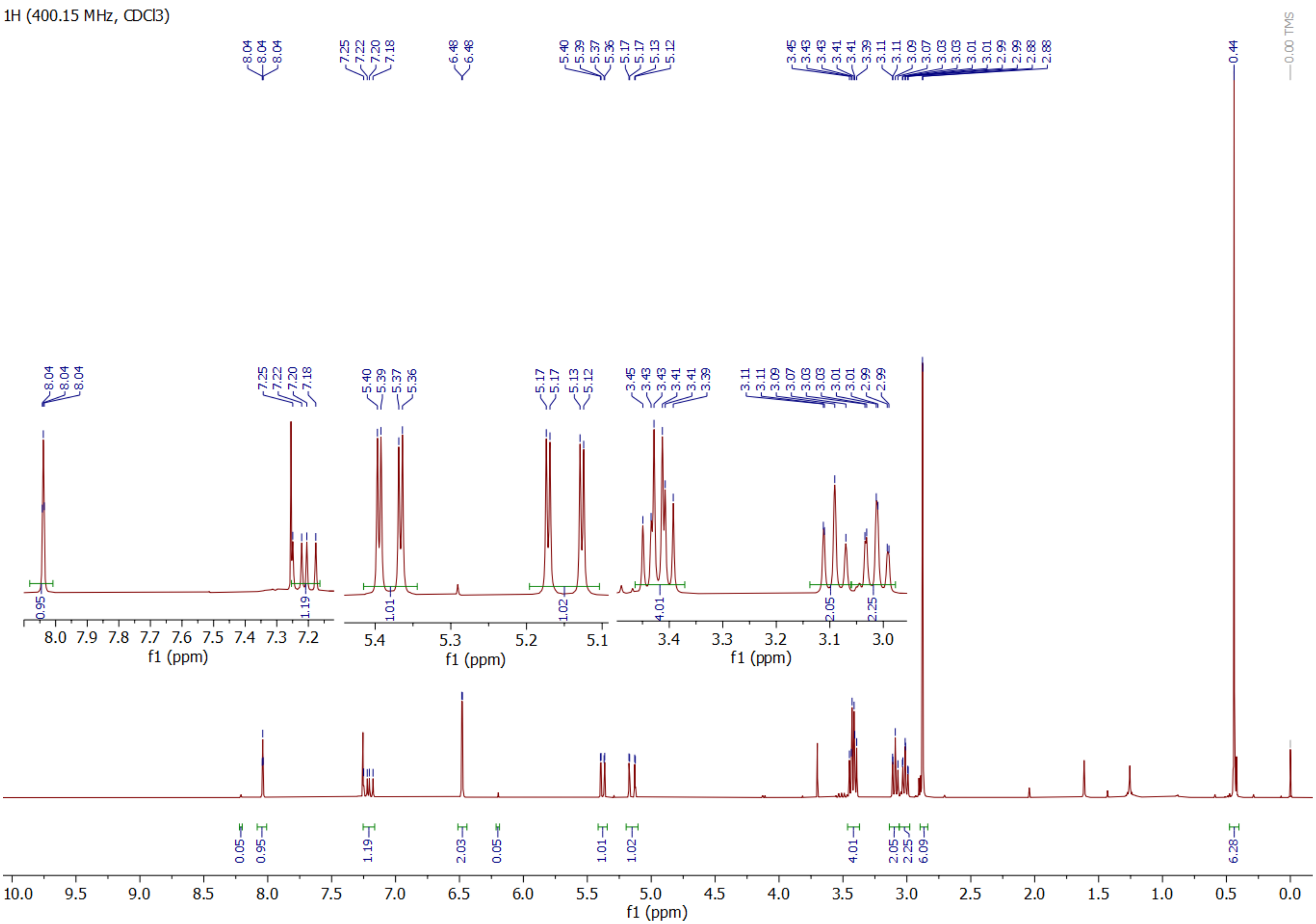

**Compound S13 $^{13}C$**

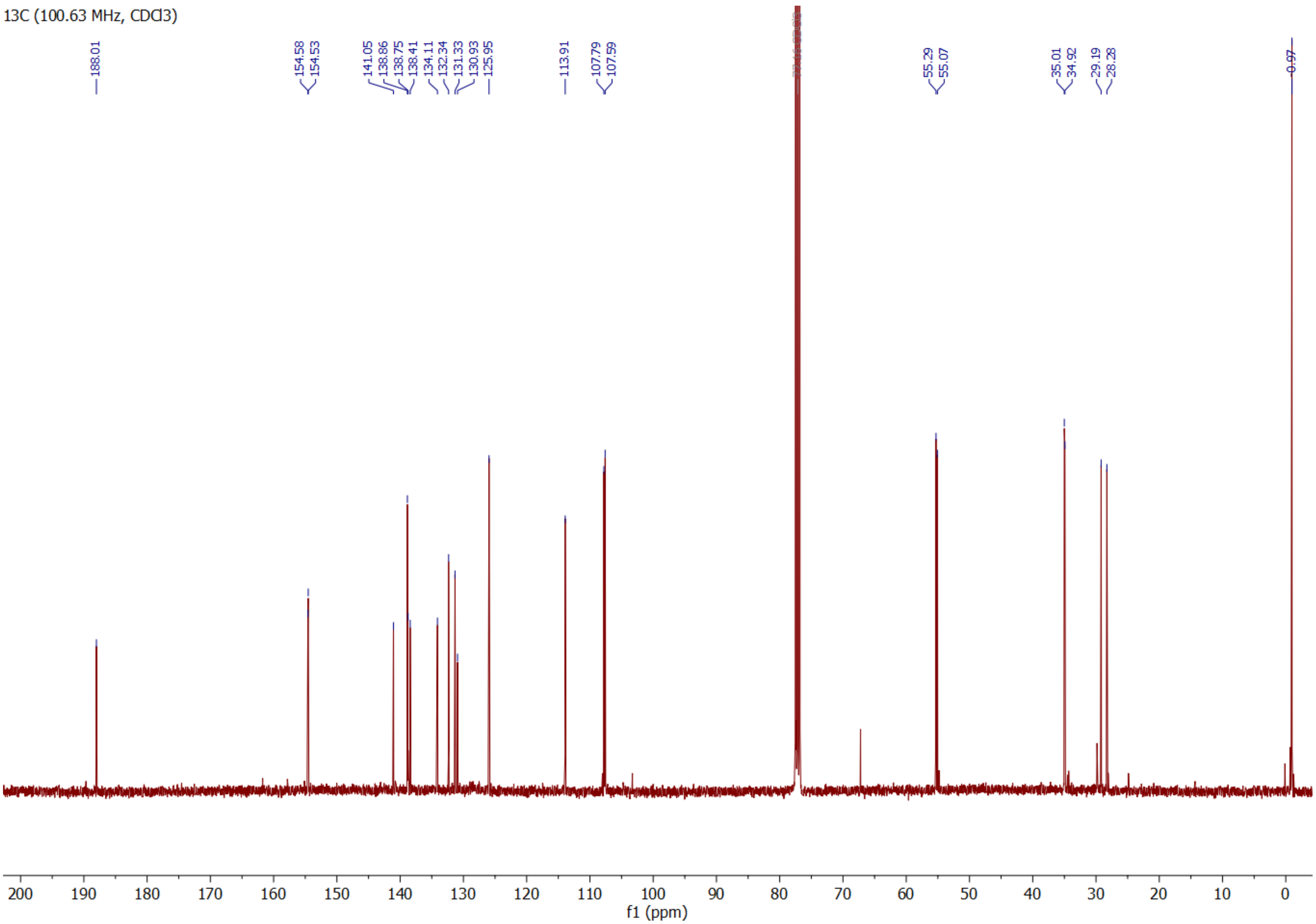

**Compound S14 $^{1}$H**

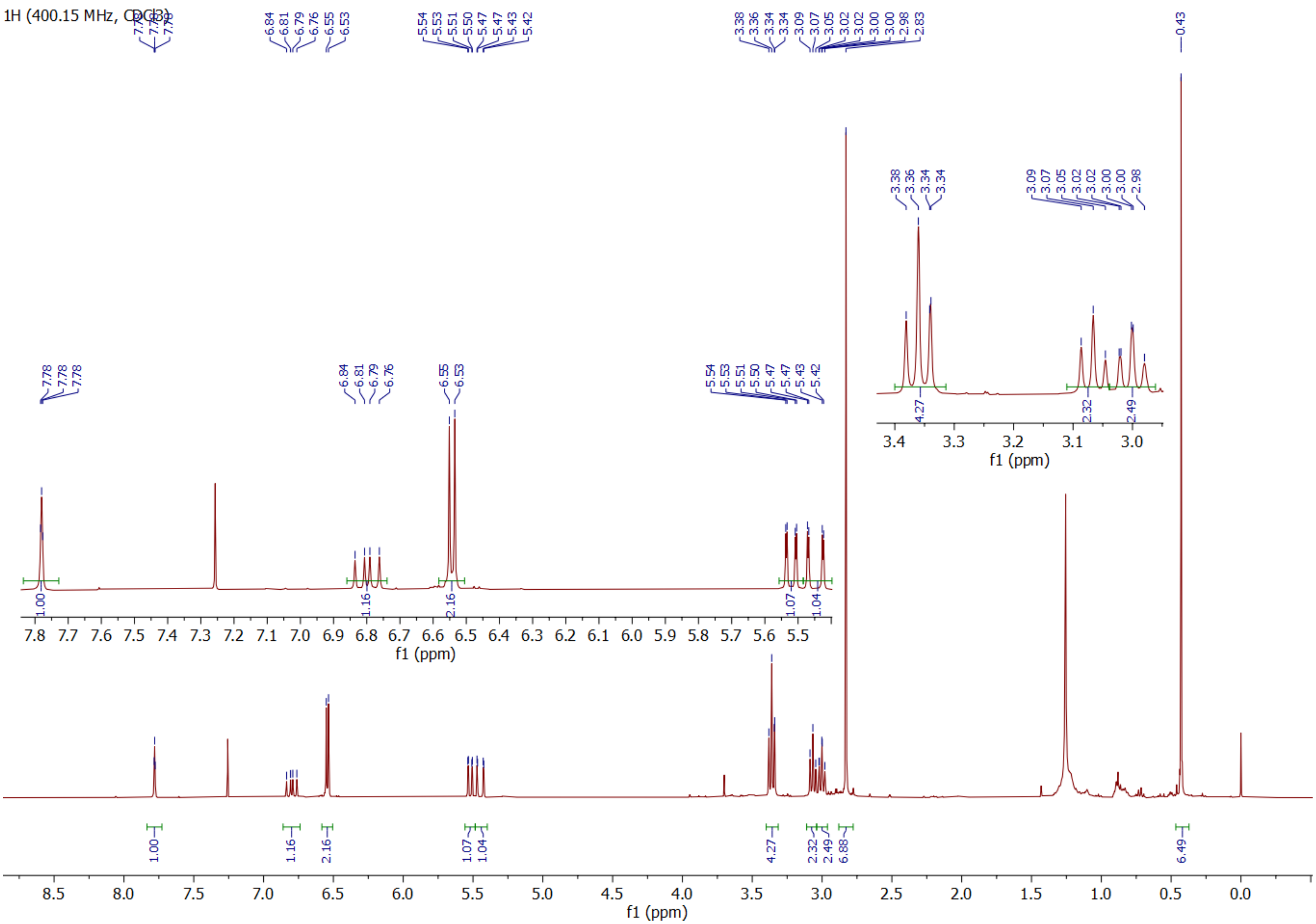

**Compound S14 $^{13}$C**

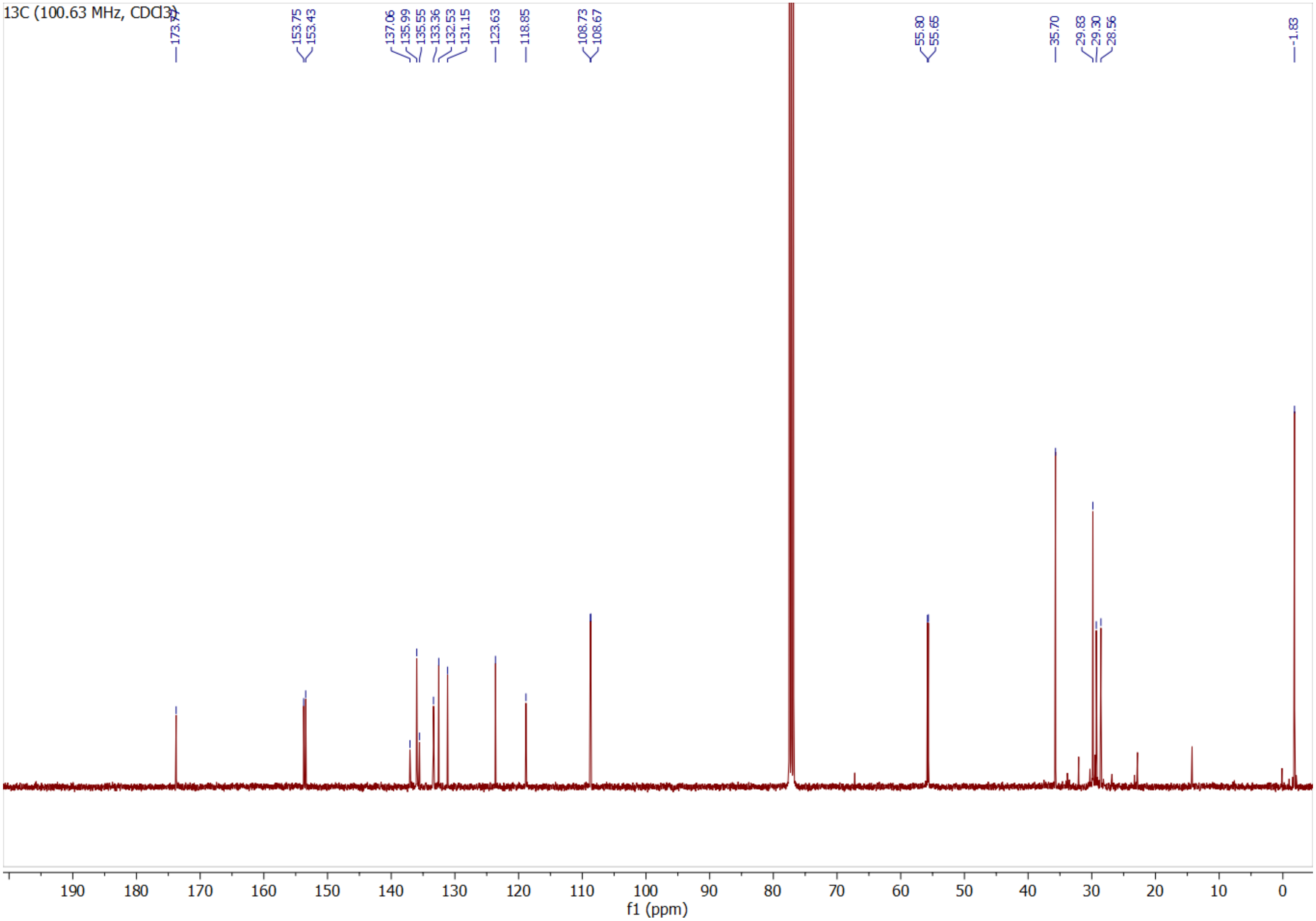

**Compound 5 $^{1}$H**

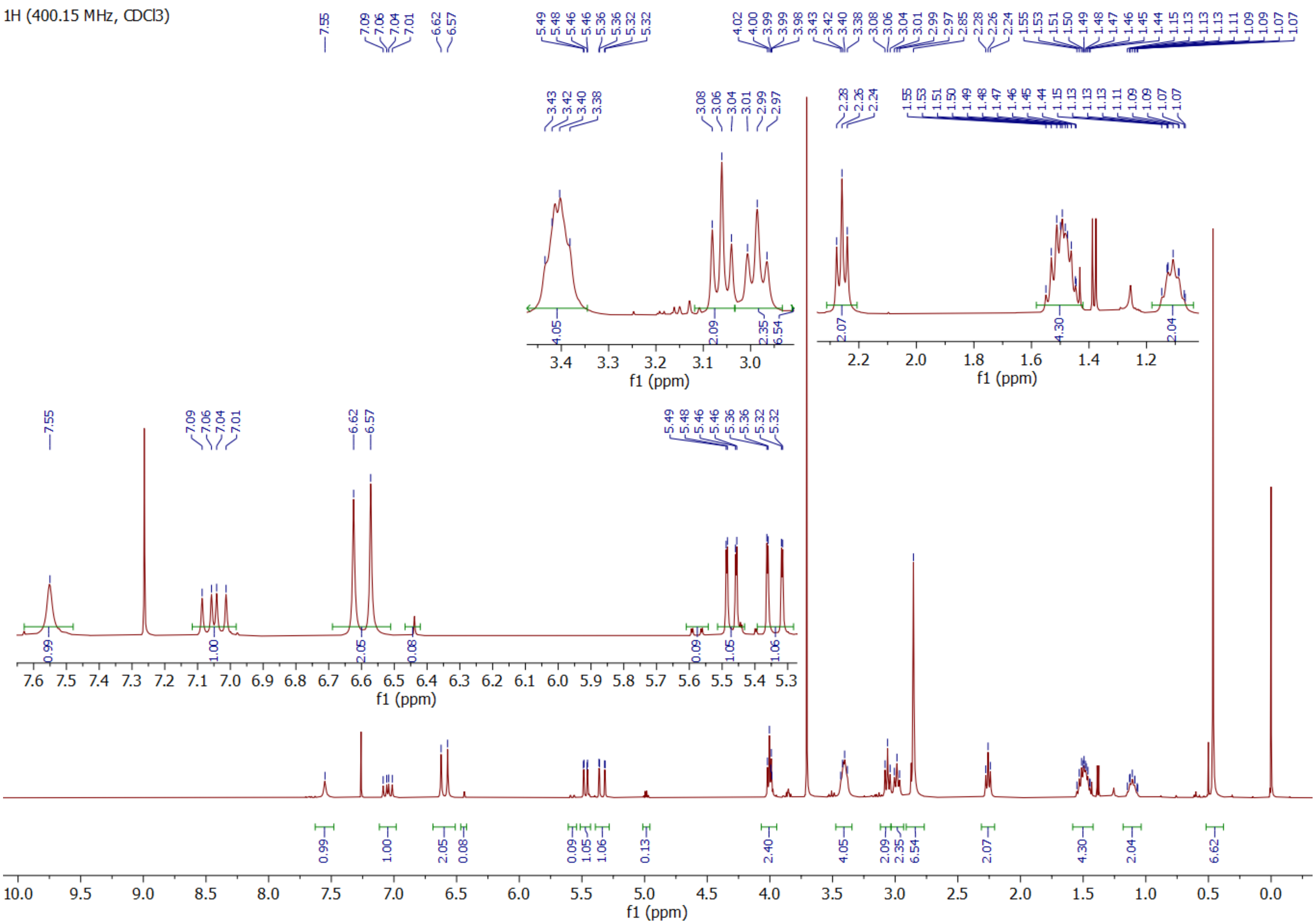

### Compound 5 gHSQCad

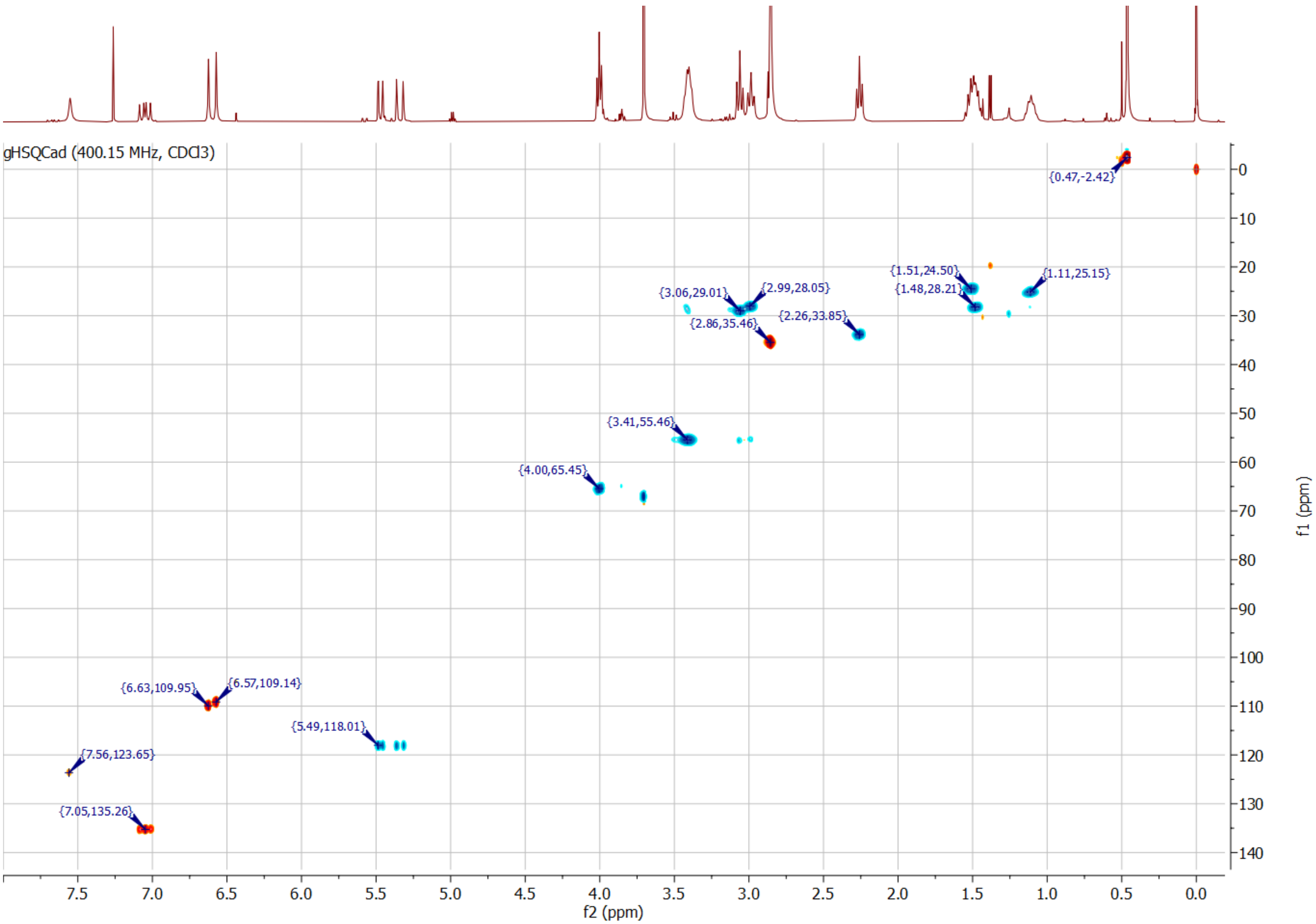

**Compound 5-Halo 1H**

1H (400.15 MHz, CDCl3)

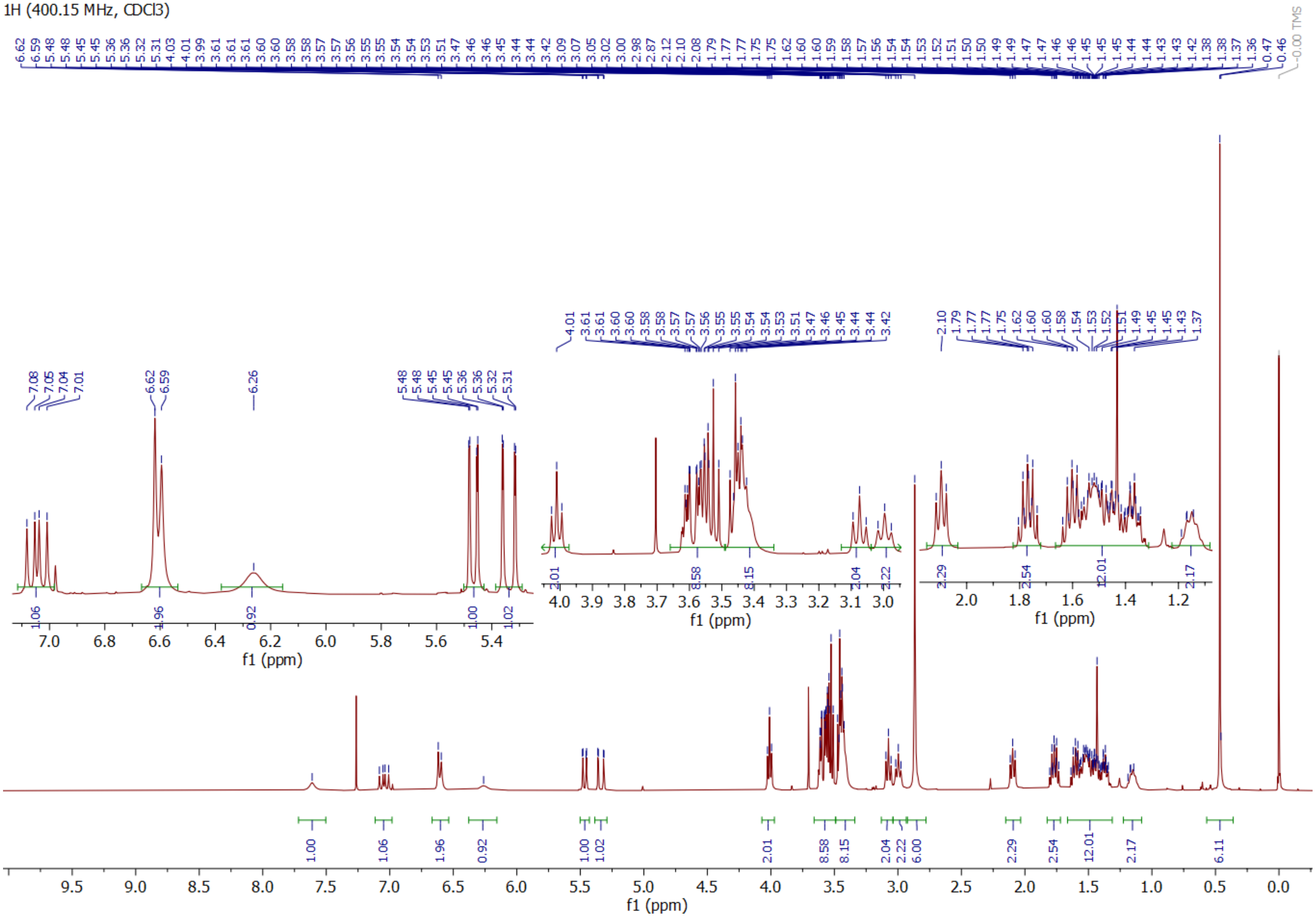

**Compound 5-Halo gHSQCad**

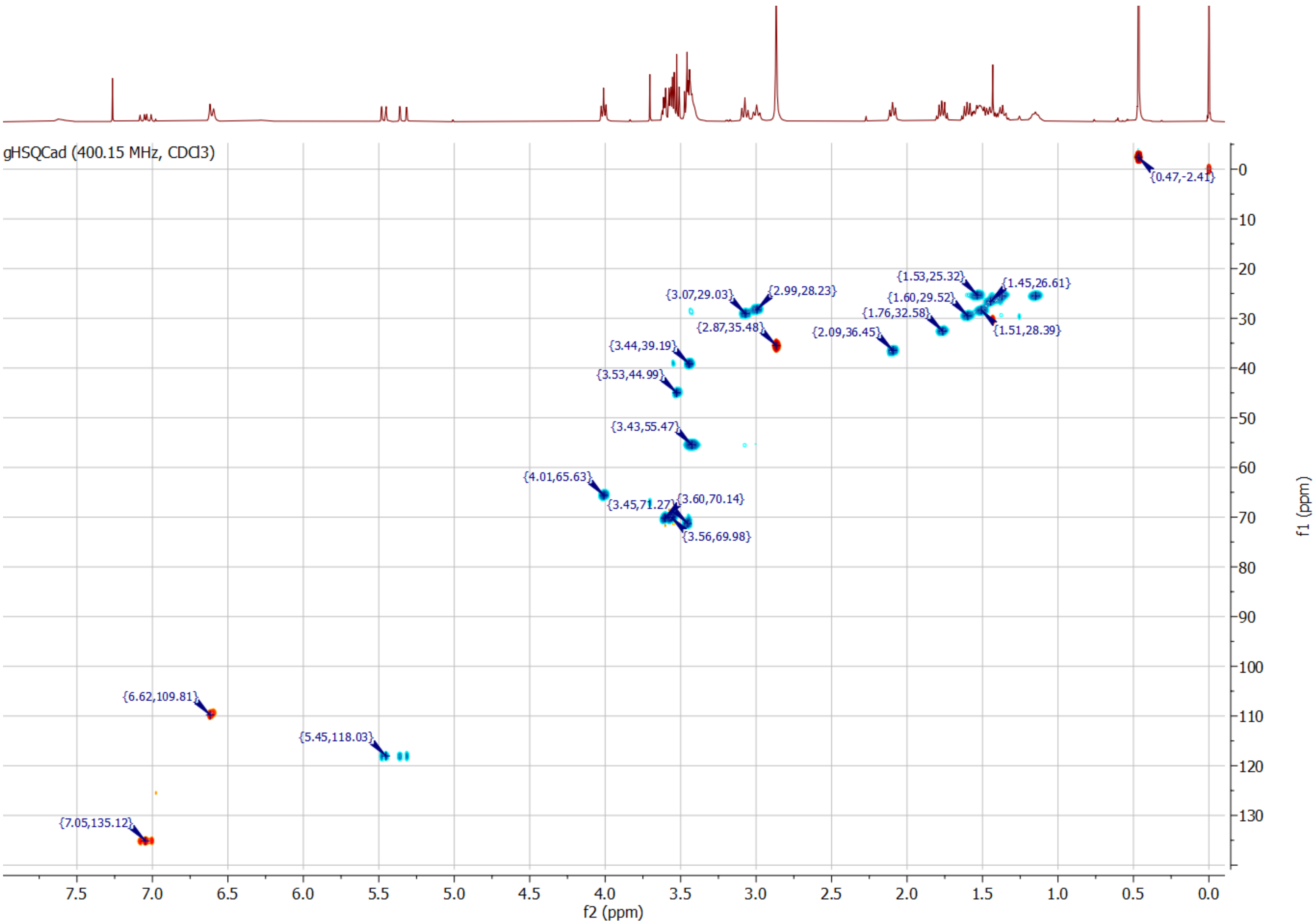
